# A pangenomic approach reveals the sources of genetic variation fueling the rapid radiation of Capuchino Seedeaters

**DOI:** 10.1101/2025.03.19.644153

**Authors:** María Recuerda, Simón Kraemer, Jonas R. R. Rosoni, Márcio Repenning, Melanie Browne, Juan Francisco Cataudela, Adrián S. Di Giacomo, Cecilia Kopuchian, Leonardo Campagna

## Abstract

The search for the genetic basis of phenotypes has primarily focused on single nucleotide polymorphisms, often overlooking structural variants (SVs). SVs can significantly affect gene function, but detecting and characterizing them is challenging, even with long-read sequencing. Moreover, traditional single-reference methods can fail to capture many genetic variants. Using long-reads, we generated a Capuchino Seedeater (*Sporophila*) pangenome, including 16 individuals from seven species, to investigate how SVs contribute to species and coloration differences. Leveraging this pangenome, we mapped short-read data from 127 individuals to perform FST scans and genome-wide association studies. Species divergence primarily arises from SNPs and indels (< 50 bp) in non-coding regions of melanin-related genes, as larger SVs rarely overlap with divergence peaks. One exception was a 55 bp deletion near the *OCA2* and *HERC2* genes, associated with feather pheomelanin content. These findings support the hypothesis that the reshuffling of small regulatory alleles, rather than larger species-specific mutations, accelerated plumage evolution leading to prezygotic isolation in Capuchinos.

**Teaser:** SNPs and indels primarily shape species differences and plumage diversity in Capuchino Seedeaters.

## Introduction

Mutations, the raw material on which evolutionary forces act, exist in different forms with unique properties. Single nucleotide polymorphisms (SNPs), small insertion/deletions (indels), and different types of structural variants (SVs) —encompassing insertions, deletions, duplications, inversions or translocations generally larger than 50 bp— differ in aspects such as their prevalence in the genome and their overall size (i.e., the number of nucleotide bases involved in the variant) (*1*). These diverse mutation types may also influence the evolutionary process in different ways. For example, because recombination within inversions is suppressed, when inversions involve multiple genes, these genes can co-evolve into what is known as a supergene, which can shape complex phenotypes (*2*). In contrast, the influence of multiple SNPs on a given phenotype can be broken down by recombination, hindering the ability of such variants to collectively shape a trait unless recombination and/or gene flow are suppressed.

Nevertheless, questions related to the evolutionary significance of SVs, like if larger variants generally lead to more complex evolutionary changes, remain unanswered (*1*). Additionally, the nature of the mutational process and the surrounding genomic context, can further shape the type of genetic variation available for evolution. For example, a copy number mutation in a repetitive region of the genome may be more likely (through replication slippage) than a point mutation (*3*). Transposable elements (TEs), mobile segments of DNA that can copy themselves and integrate in different parts of the genome, contribute to generating mutations and shaping genome evolution (*4*). TE-derived mutations are not necessarily random, as TEs can preferentially integrate in certain areas of the genome and are more prevalent in certain genomes versus others (*5*, *6*). Therefore, the rate of genetic change may depend on the type of genetic variants involved, and the availability of mutations will partially determine the pace of the evolutionary process. Rapid speciation may be fueled by the availability of novel genetic variation, a process that can be further accelerated by gene flow and recombination. Like mutations, these mechanisms can also introduce genetic variants into different genomic backgrounds, providing new genomic variation for evolution to act on (*7*). It is generally unknown whether certain types of mutations can more rapidly lead to the evolution of new traits and species, although there is a growing literature on the evolutionary importance of chromosomal inversions (*1*, *8*).

There are several methodological reasons why SNPs are the most common type of genetic marker used in genomic studies of non-model organisms (*1*, *9*). First, the prevalence of short-read sequencing technologies complicates the assembly of complex repetitive regions of the genome, as these reads typically do not span such regions, leading to incorrect assemblies (*10*). Consequently, repetitive areas like those rich in TEs are either incorrectly assembled or broken into many small scaffolds, resulting in these types of genomic regions (and mutations) being underrepresented in genomic studies. Second, it is a common practice to assemble a single reference genome, either from the focal study species or a closely related one, and subsequently conduct population-level whole-genome re-sequencing of a larger number of individuals that are mapped back to this reference genome (*11*). This approach will miss SVs that are absent from the reference genome, as variants in a given population or species that are not represented in the reference (for example an insertion) will be lost in the alignment process (*12*). Finally, although small indel mutations can still be recovered from short-read sequencing data, most of the bioinformatics pipelines and population genomic software for downstream analyses are based on SNP data, on which many researchers tend to focus (*13*). Thus, the growing number of genomic studies on non-model organisms has primarily focused on SNPs in non-repetitive regions of the genome to assess patterns of genetic variation. We comparatively know far less about how other types of mutations contribute to evolution, especially in non-model systems.

The use of long-read sequencing technologies can produce higher quality reference genomes by spanning repetitive regions and thus improving genomic assemblies (*10*). Moreover, the use of these technologies with approaches that leverage the combination of several reference genomes into a pangenome, can capture a more complete representation of the genomic variation in a species or population (*14*). Ideally, a pangenome represents the full spectrum of variation in a species, capturing the different types of genetic variants which can be present in a single individual, population, or the entire sample. The use of pangenomes has the potential to help mitigate the bias against genetic variants that have been traditionally harder to detect in evolutionary studies of non-model organisms, offering a more comprehensive view of genetic variation, including rare and population-specific SVs.

In this study we focus on a rapid radiation of 12 bird species in the genus *Sporophila* known as the Capuchino Seedeaters (*15*, *16*). Capuchinos differ primarily in adult male plumage and vocalizations, traits that in these species mediate assortative mating, yet show low genome-wide genetic differentiation (FST ∼ 0.008; (*17*)). While coloration differences are inherited genetically, song evolution is a mostly cultural process in songbirds (but see (*18*)). Male coloration differences between Capuchino species follow a modular pattern, with distinct patches (e.g., throat, belly, cap) consistently varying in a series of colors (e.g., black, cinnamon, white). For example, the Dark-throated Seedeater (*Sporophila ruficollis*), the Tawny-bellied Seedeater (*Sporophila hypoxantha*), and the Marsh Seedeater (*Sporophila palustris*) differ by having black, cinnamon or white throats, respectively. Despite the overall genomic homogeneity, previous studies have identified a small number of narrow genomic regions with elevated differentiation, many of which are near genes involved in the melanogenesis pathway (*17*, *19*, *20*), and have undergone selective sweeps (*21*). Genetic changes in these regions containing melanogenesis genes are strongly associated with variation in the composition of melanin pigment types and their deposition across different body parts in the Capuchinos (*20*).

The genetic variants which are candidates for controlling plumage coloration are predominantly non-coding SNPs near otherwise conserved pigmentation genes (*17*, *20*). These non-coding regions are in some cases conserved across more distantly related species, suggesting they could serve important regulatory functions (*17*). The outlier regions are repeatedly involved in the divergence between different Capuchinos and generally do not contain species-specific variants, but rather have shared haplotypes among species in unique combinations across the different divergence peaks (*17*, *19*). For example, the Iberá Seedeater (*Sporophila iberaensis*) and *S. ruficollis*, both with black throats, share genotypes near the *TYRP1* gene, yet differ in a genomic region close to the *HERC2* and *OCA2* genes, which is in turn also shared between *S. iberaensis* and other Capuchinos (*19*). The unique combinations of genotypes across multiple outlier regions may underlie the emergence of novel coloration phenotypes (*7*, *19*). Taken together, these findings suggest that the sharing and reshuffling of regulatory alleles at pigmentation genes (e.g., (*22*)) may have been the engine behind the generation of novel plumage patterns. These phenotypic differences function in mate-recognition, leading to the establishment and maintenance of species boundaries early in the speciation process (*19*). Additionally, the Z sex chromosome plays a disproportionate role in species differences (*17*), potentially contributing to rapid evolution, as has been described in other systems (*23*).

However, these findings are based on genomic studies that employed a single reference genome from a *S. hypoxantha* male sampled in the Esteros del Iberá, Argentina, which was primarily assembled using short-read sequences (*17*). Moreover, the FST outlier scans and genome-wide association studies (GWAS) were conducted exclusively using SNPs. It is therefore possible that the variation in non-coding SNPs near melanogenesis genes in the Capuchinos is accompanied by other, yet undetected genetic changes, such as species-specific SVs (perhaps generated by transposable element activity) absent in the *S. hypoxantha* individual used to assemble the reference genome. A preliminary analysis using long-read sequences to compare a single Pearly- bellied Seedeater (*Sporophila pileata*) individual to the *S. hypoxantha* reference genome found ∼500 SVs between these two individuals, yet only four inversions (∼450 bp) were located within the areas of genomic divergence (*17*). This result shows that SVs may be present in at least some divergence peaks, but their prevalence and level of differentiation across species remain unknown.

Here we aim to assess the relative contribution of different types of mutations to the evolution of Capuchinos, with the goal of achieving a better understanding of the genomic changes promoting rapid speciation. To this end, we assembled a pangenome from 16 individual reference genomes generated *de novo* through high-coverage Pacific Biosciences long-read sequencing. This Capuchino pangenome combined information from males and females of the seven species present in the area showing the highest sympatry in this group, the Esteros del Iberá in the Province of Corrientes, Argentina (*17*, *19*). We subsequently combined this pangenome with information from previously published and new short-read whole genome resequencing data for all Capuchinos, obtaining genotypes for these individuals for SNPs, indels and SVs. We used this information in FST outlier scans and coloration GWAS to ask how the different types of markers contribute to species divergence. We find a high level of synteny among the Capuchino genomes and that the differences in the previously identified divergence peaks are primarily shaped by SNPs and small indels (< 50 bp), mainly in non-coding regions which in some cases coincide with annotated TEs. Although we can detect larger SVs, these tend to segregate at low frequencies, and generally do not associate to divergence peaks. Our study strengthens the hypothesis that the shuffling of regulatory alleles between Capuchinos has promoted the rapid evolution of plumage traits, which lead to prezygotic reproductive isolation early in the speciation process.

## Results

### High similarity among reference genome assemblies from seven Capuchino species

We recovered two genome assemblies per diploid individual, a higher quality primary haplotype and an alternate one. The primary haplotypes were longer (1.14 Gb vs. 1.09 Gb in average length) and more contiguous than the alternate ones, containing approximately one-third as many scaffolds (315 vs. 1,075), a tenfold higher average N50 (31 Mb vs. 2.9 Mb), and an eightfold smaller L50 (13 vs 112 scaffolds) (Figure 1A). We did not observe large differences in these statistics across species, and the primary assemblies showed high synteny when compared visually (Figure 1B). The proportion and distribution of different types of repetitive and transposable elements (TEs) were very similar across primary assemblies and species, encompassing around 16% of the genome, and dominated by ∼11% retrotransposons (Table S1, Figure 1C). Gene content showed an average of 14,666 ± 46 (SD) genes per assembly, with little variation across individuals and species (Table S2). Gene completeness was similar between the primary assemblies, alternate assemblies, and gene annotations across species, with the primary assemblies being slightly more complete (96.5% single-copy orthologs) (Tables S3, S4; Figure 1D). Regarding gene identity, when we add genomes sequentially to our analysis, we reach a plateau at 15,788 unique genes, suggesting we can capture a greater number of genes by including multiple genomes (Figure 1E). Our analysis of gene presence/absence variation (PAV) using the annotations from the different assemblies initially suggested there are several genes present uniquely in each species (between 25 for *S. pileata* and 100 for *S. iberaensis*) (Figure S1, S2). However, upon further analysis we believe this is related to the quality of the assemblies and annotations, and not true missing genes, as a BLAST search found fragments or complete gene sequences for these putatively missing genes in the assemblies where they were initially thought to be missing (Table S5).

**Figure 1.**
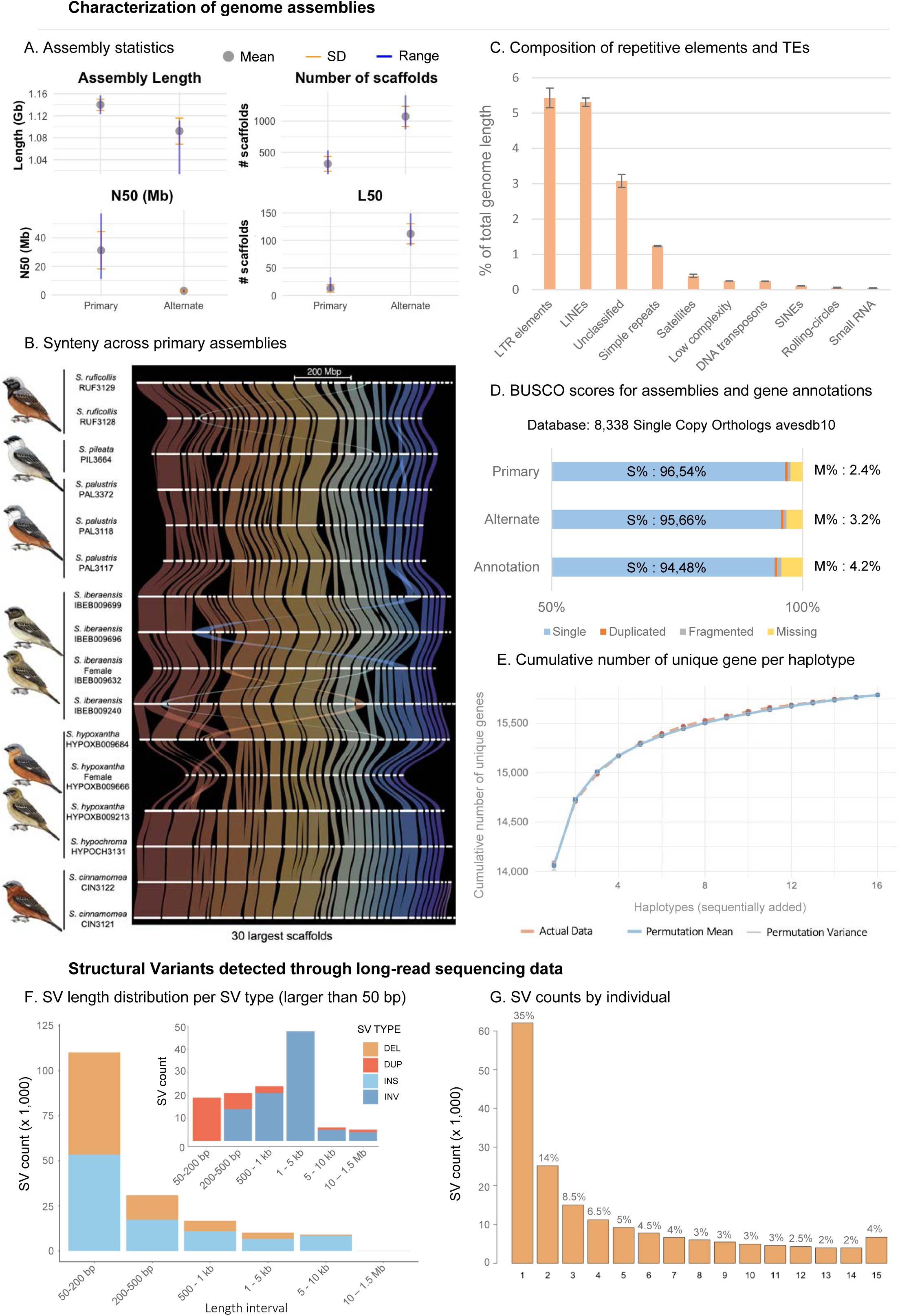
Genome assemblies and statistics for SVs called from long-read data. **A)** Length and contiguity statistics for the primary and alternate assemblies. **B)** Synteny representation of the 30 longest scaffolds across all primary assemblies generated using GENESPACE. **C)** Composition of transposable elements (TEs) in the primary assemblies. **D)** Evaluation of gene annotations through BUSCO analyses. Mean BUSCO scores for all primary and alternate assemblies, as well as the annotations for primary assemblies. **E)** Cumulative unique gene count per haplotype using the annotations, showing actual data (red dots, dashed line) and the expected curve based on 1,000 permutations (blue dots, solid line), with variation depicted by the grey bars. **F and G)** SV statistics from long-read sequencing data. **F)** SV length distribution per SV type, based on results from Sniffles2 supported by three SV callers (Sniffles2, PBSV, and SVIM- asm). **G)** SV counts per individual, showing the number of structural variants (SVs) present in 1 to 15 individuals (the reference genome HYPOXB009684 is not included), with most SVs found in only a single individual.

### The landscape of structural variation and the Capuchino Seedeater pangenome

Long-read data enabled us to identify an average of ∼55K structural variants (SVs > 50 bp) per individual genome that were consistently supported by the three SV-calling methods that we employed. When variants from all individuals were merged, this resulted in a total of 177,246 SVs. Insertions (54.5%) and deletions (45.5%) were most common, while inversions and duplications were comparatively rare (≤0.05% each) (Figure 1F). Smaller insertions, deletions, and duplications were the most frequent, while inversions were more prevalent in the 200 bp–5 kb range and absent in the smallest interval (Figure 1F). Approximately 35% of the variants were private, found in a single individual, while only 4% were shared by all individuals (Figure 1G). For example, among the 92 inversions that we detected, only one was supported by all individuals. Given that we found a total of ∼180 K SVs by comparing each individual to a reference genome, we sought to leverage all genomes collectively to uncover additional SVs and their shared patterns by constructing a pangenome. The pangenome length, measured in the total number of unique base pairs recovered from all individuals, is 1.5 Gb, with the core genome shared by every individual comprising 66% of this length and accounting for 48% of the total nodes (Figure 2A, S3). The pangenome recovered 59.2 million variants, with 7.6 times more SNPs than SVs (Figure 2B). Nearly half of the variants (45%) were rare, appearing only once as the alternate allele among the 16 individuals. Within SVs, 96.5% are shorter than 50 bp (i.e., indels; Figure 2B). To understand the patterns of differentiation of these markers among the different Capuchino species, we combined the pangenome with short-read sequencing data from a larger sample of birds. We therefore used the pangenome as a reference to map short-read data, resulting in a dataset of 127 individuals across 10 species, and obtained genotypes, including SVs, for this larger sample. This approach enabled us to perform GWAS and FST outlier scans to compare the patterns of genomic differentiation observed from the different types of markers (e.g., SNPs, indels, SVs).

**Figure 2.**
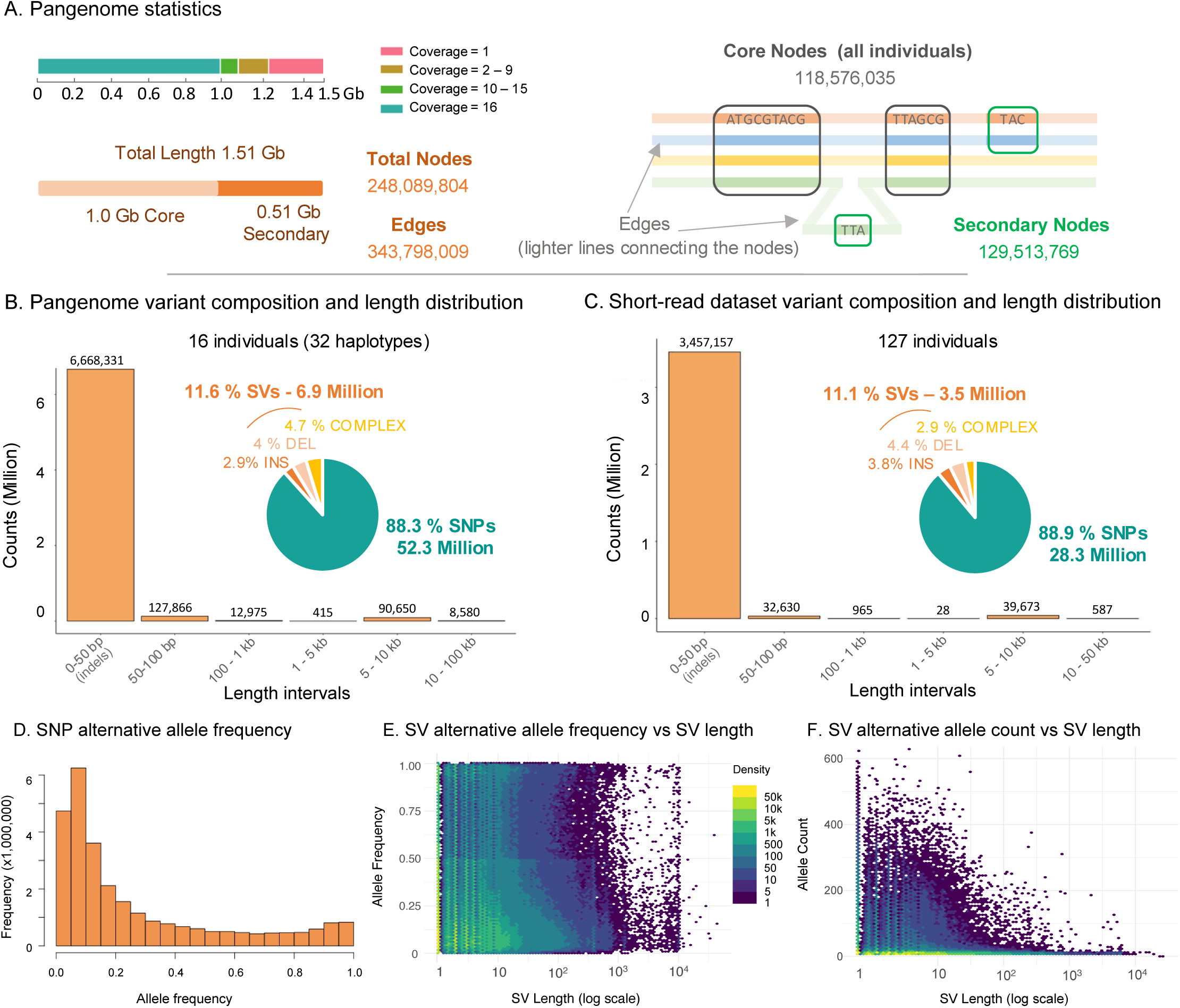
The Capuchino pangenome and variant genotyping using short-read data. **A)** Overview of statistics describing the pangenome, including the shared sequence length (Gb) across varying numbers of haplotypes as computed with Panacus, total length, number of nodes and edges, and the lengths and counts of core and secondary genome nodes. **B)** Length distribution and composition of SVs in the pangenome, highlighting total numbers and percentages of SNPs and SVs. SVs are further categorized into insertions, deletions, and complex rearrangements. **C)** Length distribution and composition of variants genotyped for 127 individuals from short-read data mapped to the pangenome, showing total numbers and percentages of SNPs and different types of SVs. **D)** Alternative allele frequency distribution for SNPs identified from short-read data mapped to the pangenome (see Figure S5A for the equivalent plot for SVs). **E)** Relationship between SV length (in log scale) and alternative allele count, and **F)** alternative allele frequency. In both **E** and **F** the density of data points is represented by a color gradient, with purple and yellow indicating lower and higher densities, respectively.

After genotyping the larger sample of individuals, we recovered ∼35.5 million variants, which after filtering (31.8 million) is slightly over half of the variants in the pangenome. Regardless, the proportions between SNPs and SVs remained similar, with 8 times more SNPs than SVs (28.3 vs 3.5 million), and only 2.7% of variants exceeding 50 bp (Figure 2C). We recovered ∼5.9 million variants in this larger dataset that were invariant in the pangenome, likely due to including more species and individuals per species. However, ∼29.6 million variants present in the pangenome were lost during short-read mapping and genotype calling. This is primarily due to the limitations of short-read data in resolving more complex, repetitive regions of the genome, which are lost during mapping. The mean mapping quality (± SD) of short-reads in regions present in the pangenome but missing in the short-read dataset was 3.86 ± 6.53, significantly lower than 55.3 ± 7.64 in regions present in the short-read dataset (Wilcoxon rank sum test: W = 127, p < 2.2e-16). Most of these problematic areas are located at the ends of scaffolds, enriched in repetitive and transposable elements (TEs). They often also show lower coverage in the pangenome dataset, suggesting they are inherently hard to resolve across sequencing platforms, but cannot be recovered with our short-read data (Figure S4).

To conduct GWAS and outlier analyses, we filtered the short-read dataset so that SVs were present in at least 80% of individuals (discarding variants in fewer than 102 individuals). Both the SNP and SV alternative allele frequencies were characterized by a higher abundance of loci with low-frequency alleles, with a gradual decline in abundance toward intermediate and high frequencies (Figure 2D, 2E, Figure S5A). The lower count in the lowest bin of allele frequencies in both the SNP and SV distributions is due to the missing data and allele count filters (Figure S5B and C). The overall density of SVs declined with their increasing length, indicating that longer SVs are less common (Figure 2E, 2F). Additionally, we found an inverse relationship between SV length and alternative allele frequency and count, with shorter SVs having higher allele counts (Figure 2F). The alternative allele frequency distribution demonstrated that low-frequency variants are prevalent across all SV lengths, particularly at shorter lengths (Figure 2E).

### Outlier genomic regions associated to plumage coloration and species differences

We conducted genome-wide association analyses across six different body parts using the concentration of eumelanin or pheomelanin in the feathers from these plumage patches as phenotypes. To explore the contribution of different types of markers to pigmentation, we partitioned the data set by variant type (e.g., SNPs, SVs). The GWAS revealed a total of seven strong outlier peaks repeatedly associated with eumelanin or pheomelanin concentration across the six plumage patches (Figure 3, Figures S6-S11). These peaks were consistently observed in different combinations depending on the plumage patch and pigment type. Five peaks were detected in both the SNP and SV datasets, encompassing the melanogenesis genes *OCA2/HERC2*, *ASIP*, *TYRP1* and *SLC45A2*, as well as the *AHCY* and *GPT2* genes, which are involved in amino acid degradation. The remaining two peaks were identified exclusively using the SNP dataset and did not contain annotated genes within them (Table 1, Table S6). We did not observe strong associations with eumelanin in the head and pheomelanin in the throat (Figure S8, S11). Most peaks were identified through SNPs, with only a few associated with indels (SVs < 50 bp). A single larger SV—a 55 bp deletion—was found within a peak associated with pheomelanin concentration in the belly plumage patch (Figure 3). Genetic variants within repetitive elements and annotated TEs recovered fewer peaks compared to variants outside these regions (Figure 3, Figures S6-S11). The GWAS results also identified 217 singletons (isolated SNPs and SVs, representing 24.5% of all significant GWAS hits) not included in the more prominent outlier peaks (Table S7).

**Figure 3.**
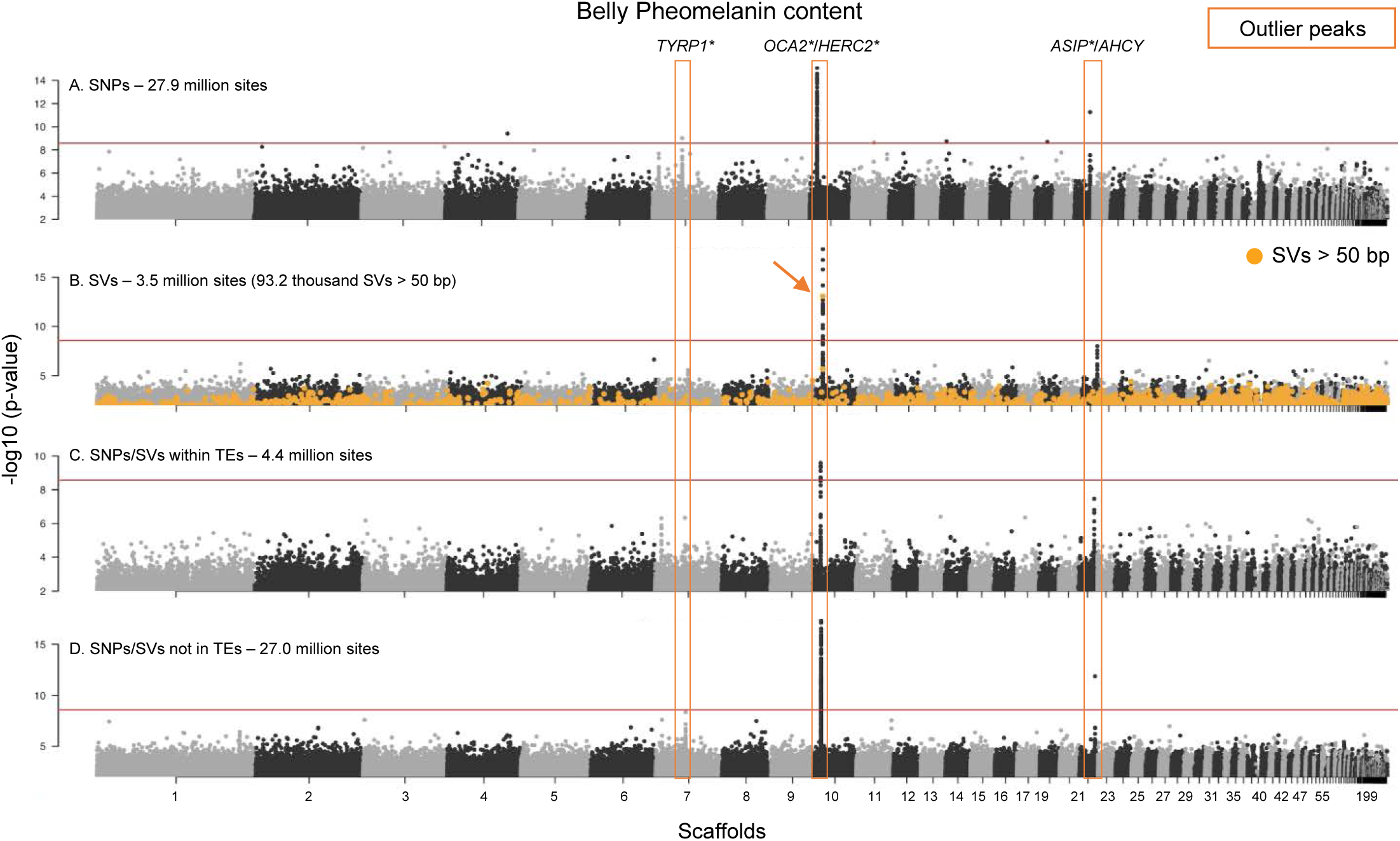
Genome wide association study for the pheomelanin content in the belly plumage patch. The analysis includes four datasets, displayed from top to bottom: **A)** SNPs, **B)** Structural variants including indels (< 50 bp) and long variants (in orange), **C)** Variants (SNPs and SVs) within repetitive and transposable elements (TEs), and **D)** Variants (SNPs and SVs) outside TEs. The y axis represents the -log10(p-value) obtained in the GWAS and the red line is the Bonferroni-corrected threshold of statistical significance (for all comparisons and variants), corresponding to a p-value of 2.65 *10^-9^. Scaffolds are ordered by decreasing size and represented in alternating black and gray. Peaks are highlighted with an orange rectangle, and known genes are labeled above the peaks. The genes marked with an asterisk (*) belong to the melanogenesis pathway. The peaks associated with pheomelanin content in the belly include *TRYP1* on scaffold 7, which is detected only by the SNPs dataset; *OCA2*/*HERC2* on scaffold 10, which is detected by all datasets, including the long SVs; and the peak containing the *ASIP* and *AHCY* genes on scaffold 21, which is detected by the SNPs and the variants outside the TEs datasets. The single large SV with a statistically significant association is marked with an arrow.

**Table 1.** GWAS and FST outlier peaks. Details for the 10 main outlier regions identified using both strategies, including peak coordinates (scaffold, end and length), the chromosomal location according to the Zebra finch genome, whether the peak was identified as a GWAS and/or FST r, the number and type of variants (SNPs and SVs) and their overlap with transposable elements (TEs) and coding regions, and the genes within peak. Additionally, the table provides the maximum, mean, and total length of the SVs within each peak. Peaks detected with both SNPs and re highlighted in bold, while those detected only with SNPs are in regular font. The peaks marked with an asterisk (*) are those that were also ered by the variants located within TEs. # In peak 1 we refer to the *OCA2*/*HERC2* gene pair which is involved in melanogenesis, yet the ic gene in the peak is *HERC2*. In Peak 3, the total length is shorter than the maximum length (§) because the longest variant is not fully ined within the peak—3,544 bp extend beyond its boundaries.

|  | Peak 1* | Peak 2 | Peak 3* | Peak 4* | Peak 5* | Peak 6 | Peak 7 | Peak 8* | Peak 9 | Peak 10 |
| --- | --- | --- | --- | --- | --- | --- | --- | --- | --- | --- |
| Scaffold | **10** | 14 | **21** | **7** | 9 | 41 | **7** | **3** | 7 | 7 |
| Chromosome | 10 | 11 | 20 | Z | 6 | Z | Z | 4 | Z | Z |
| Start coordinates (bp) | 6,190,001 | 13,950,001 | 13,545,422 | 25,260,001 | 15,010,001 | 108,180 | 5,260,234 | 9,950,001 | 5,560,001 | 5,730,001 |
| End coordinates (bp) | 6,250,000 | 13,990,000 | 13,700,000 | 25,320,000 | 15,030,000 | 112,141 | 5,272,125 | 9,990,000 | 5,580,000 | 5,750,000 |
| Peak length (bp) | 59,999 | 39,999 | 154,578 | 59,999 | 19,999 | 3,961 | 11,891 | 39,999 | 19,999 | 19,999 |
| GWAS/ F<sub>ST</sub> | GWAS/F<sub>ST</sub> | GWAS/F<sub>ST</sub> | GWAS/F<sub>ST</sub> | GWAS/F<sub>ST</sub> | GWAS/F<sub>ST</sub> | GWAS | GWAS | F<sub>ST</sub> | F<sub>ST</sub> | F<sub>ST</sub> |
| Gene1 | *HERC2* | *GPT2* | *ASIP* | *TYRP1* |  |  | *SLC45A2* | *ALB* |  | *SPEF2* |
| Gene2 | *OCA2*# | *CDCA9* | *AHCY* |  |  |  |  |  |  | *LOC112530520* |
| SNPs# | 1,375 | 1,161 | 2,683 | 1,402 | 335 | 104 | 133 | 269 | 180 | 269 |
| SVs# | 181 | 149 | 342 | 161 | 49 | 15 | 20 | 31 | 19 | 32 |
| SNPs# within TEs | 147 | 19 | 984 | 263 | 23 | 52 | 7 | 36 | 37 | 12 |
| SVs# within TEs | 19 | 4 | 153 | 40 | 3 | 10 | 1 | 12 | 4 | 6 |
| TEs# per peak | 25 | 8 | 122 | 39 | 8 | 4 | 5 | 31 | 12 | 8 |
| Length TEs | 5,149 | 602 | 54,332 | 10,665 | 973 | 1,543 | 611 | 4,665 | 3,869 | 610 |
| Length SVs within TEs | 217.55 | 6 | 3,594.4 | 196.6 | 17 | 25 | 1 | 79 | 415 | 78.8 |
| SNPs coding | 47 | 23 | 12 | 12 | 0 | 0 | 9 | 8 | 0 | 7 |
| SNPs coding F<sub>ST</sub> > 0.75 | 1 | 1 | 0 | 2 | 0 | 0 | 0 | 4 | 0 | 0 |
| SVs coding | 0 | 1 (1 bp INS CDCA9) | 0 | 0 | 0 | 0 | 0 | 0 | 0 | 0 |
| #SVs > 50bp | 3 | 1 | 4 | 1 | 0 | 0 | 0 | 0 | 1 | 0 |
| #SVs > 50bp F<sub>ST</sub> > 0.75 | 1 (DEL 55 bp) | 0 | 0 | 0 | 0 | 0 | 0 | 0 | 0 | 0 |
| Max length SVs peak (bp) | 146 | 64 | 7398§ | 97 | 15 | 10 | 9 | 43 | 389 | 42 |
| Mean length SVs peak (bp) | 11.9 | 10.6 | 126.7 | 9.1 | 2.8 | 2.8 | 2.7 | 4.3 | 24.0 | 8.9 |
| Total Length SVs (bp) | 951.2 | 507.2 | 4839.9 | 571.5 | 139.5 | 41.0 | 57.0 | 160.5 | 502.0 | 140.6 |

Similarly to the GWAS results, the FST scans revealed eight differentiation peaks repeatedly implicated in several comparisons involving several pairs of species. Four of these peaks were shared between SNPs and SVs. Three peaks included melanogenesis genes (as in the GWAS, except for the absence of the peak containing *SLC45A2*), and the fourth contained the *ALB* gene (Table 1, S6). The four remaining peaks were detected only with the SNP dataset, and two of these contained annotated genes (*SPEF2*, *GPT2* and *CDCA9*). As in the GWAS, we observed only a few strong outlier peaks per pairwise comparison (Figure 4, Figures S12-S25). Two comparisons (the Rufous-rumped Seedeater (*Sporophila hypochroma*) vs. *S. hypoxantha* and *S. hypochroma* vs. the Chestnut Seedeater (*Sporophila cinnamomea*)) lacked outlier windows or peaks, based on the criteria we established to focus on the strongest patterns, suggesting there are more subtle patterns of differentiation between these species (Figures S13, S14). Again, SNPs and variants outside repetitive elements and TEs recover most peaks (Table S8). Moreover, we did not find SVs longer than 50 bp within the peaks, except for the same 55 bp deletion on scaffold 10 detected in the GWAS, present in the *S. hypoxantha* vs. *S. iberaensis* comparison (Figure 4). There were 85 FST outlier singleton windows identified outside the main peaks (representing 12.9% of all outlier windows including those in peaks; Table S9), showing that there are areas of the genome with more subtle patterns of differentiation.

**Figure 4.**
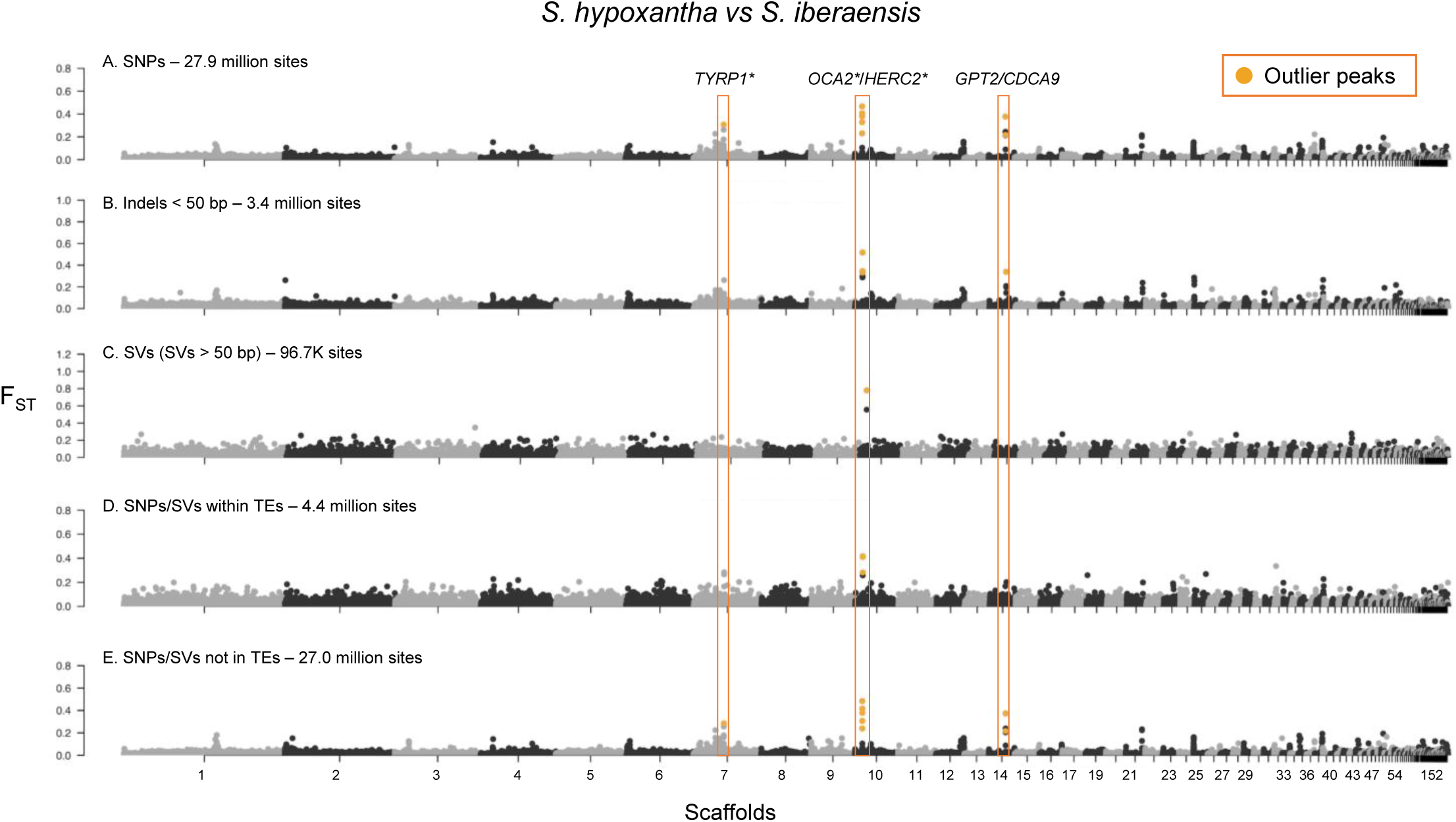
FST scan in 10 kb windows for the comparison between *S. hypoxantha* and *S. iberaensis*. The analysis includes five datasets, displayed from top to bottom: **A)** SNPs, **B)** Structural variants <5 0bp, **C)** Structural variants > 50bp, **D)** Variants (SNPs and SVs) within transposable elements (TEs), and **E)** Variants (SNPs and SVs) outside TEs. Orange dots mark outlier windows within identified differentiation peaks, defined as the top 0.1% of the FST distribution and containing at least one variant with FST > 0.75. Scaffolds are ordered by decreasing size and represented in alternating gray and black. Peaks are marked with an orange rectangle, and the known genes are labeled on top of the peaks. The genes marked with an asterisk (*) belong to the melanogenesis pathway. There are three peaks in this comparison: one containing *TYRP1* on scaffold 7, detected only by the SNP dataset; another containing *OCA2*/*HERC2* on scaffold 10, detected by all datasets; and the third containing *GPT2* and *CDCA9* on scaffold 14, detected by all datasets except the long SVs and the variants within TEs.

### Multiple sources of genetic variation in Capuchino Seedeaters

Across both the GWAS and FST outlier strategies, we detected a total of 10 peaks with an average length of 43 kb (range from ∼4 to ∼155 kb; Table 1, S6). Except for peak 10 (Table 1), the remaining regions had been identified previously as outliers using exclusively SNP markers and a single reference genome (*17*, *19*, *20*). Five peaks were shared between the GWAS and FST scans and constitute our strongest candidate regions. One was detected exclusively in the SNP dataset and did not contain annotated genes (peak 5), while the remaining four were identified in both the SNP and SV datasets (peaks 1-4) and included the melanogenesis genes *OCA2*/*HERC2*, *ASIP* and *TYRP* (Table 1, S6). Although three peaks did not contain annotated genes, they could contain regulatory loci influencing the expression of nearby genes (Table S6). Across both methods to identify outlier genomic regions, we observed consistent patterns of differentiation, with more peaks identified by SNPs and indels (preferentially outside of annotated repetitive regions and TEs), while SVs longer than 50 bp were generally absent within the peaks, except for the 55 bp deletion on scaffold 10. We found an additional 8 SVs with FST > 0.75 outside of the peaks (Table S10). Among these, two insertions in scaffolds 34 (∼207 bp) and 38 (∼73 bp) overlap with the intronic regions of the genes *SDBH* and *TPM4,* respectively, while the remaining SVs are located at distances ranging from 280 bp to ∼90 kb from the closest gene (Table S10). Both genes have been associated with reproductive traits in chickens, with *SDHB* serving as an indicator of sperm quality (*24*) and *TPM4* linked to follicle recruitment (*25*).

Finally, we employed a windowed-based PCA approach to search for large inversions that may have been missed by our pangenome. Although, this strategy found patterns consistent with a large inversion on the Z sex chromosome, it was not associated to species differences or any of the identified peaks (Figure S26).

Across outlier peaks, SNPs were more frequent and covered a greater proportion of bases than SVs, with the majority being non-coding (Figure S27A, Table 1). Repetitive elements and TEs covered between 1.4% and 37% of the peaks, with no clear pattern regarding the type of variant that overlapped these elements (Figure S27A, Table 1). The TE composition within the peaks was similar to the genome-wide pattern (Figure 1C), with a clear prevalence of retrotransposons, specifically LINES and LTRs (Figure S27B). Although the peaks did not contain extreme percentages of either SVs nor TEs compared to the genomic average, we observed greater variation in TE composition within the peaks compared to SV content (Figures S27C and S27D). Notably, half of these peaks are located on the Z sex chromosome (Table 1)—a pattern previously observed in this system (e.g. (*20*)) and others (*26*).

Only 1.5% (118 SNPs) of SNPs (irrespective of their level of differentiation) were coding variants, suggesting a minor role for coding differences overall. However, among the coding variants, eight SNPs with FST values above 0.75 (four of which were found in multiple comparisons) likely play an important role in driving color/species differentiation (Table S11). The Black-bellied Seedeater (*Sporophila melanogaster*) is involved in 9 out of the 13 comparisons and the genes affected were *TYRP1, ALB, GPT2* and *HERC2* (Table S11). We detected a single coding indel (a 1 bp insertion in *CDCA9*), but it was not highly differentiated among species (Table 1). In contrast, 11% of the variants within peaks were SVs with only 10 longer than 50 bp, of which 1 (the 55 bp non-coding deletion on scaffold 10 located 12.8 kb from the *HERC2* gene) had an FST value above 0.75 in the comparison between *S. hypoxantha* and *S. iberaensis* (Table 1, Figure S28A and S28B). In this species pair, most individuals from *S. iberaensis* are homozygous for the deletion (1/1), with some cases of heterozygosity, whereas *S. hypoxantha* individuals are predominantly homozygous without the deletion (0/0), although there are two 1/1 individuals (Figure S28C). This variant is not species-specific, as *S. palustris*, the Black-and-tawny Seedeater (*Sporophila nigrorufa*), and *S. hypochroma* exhibit genotypes similar to *S. hypoxantha* at this site, while the remaining five species share the deletion with *S. iberaensis*. The deletion is in high linkage disequilibrium (LD) with SNPs and indels within the peak, showing how different types of variants have the same genomic signal (Figure 5). We do however observe species-specific patterns when assessing variation across all peaks combined.

**Figure 5.**
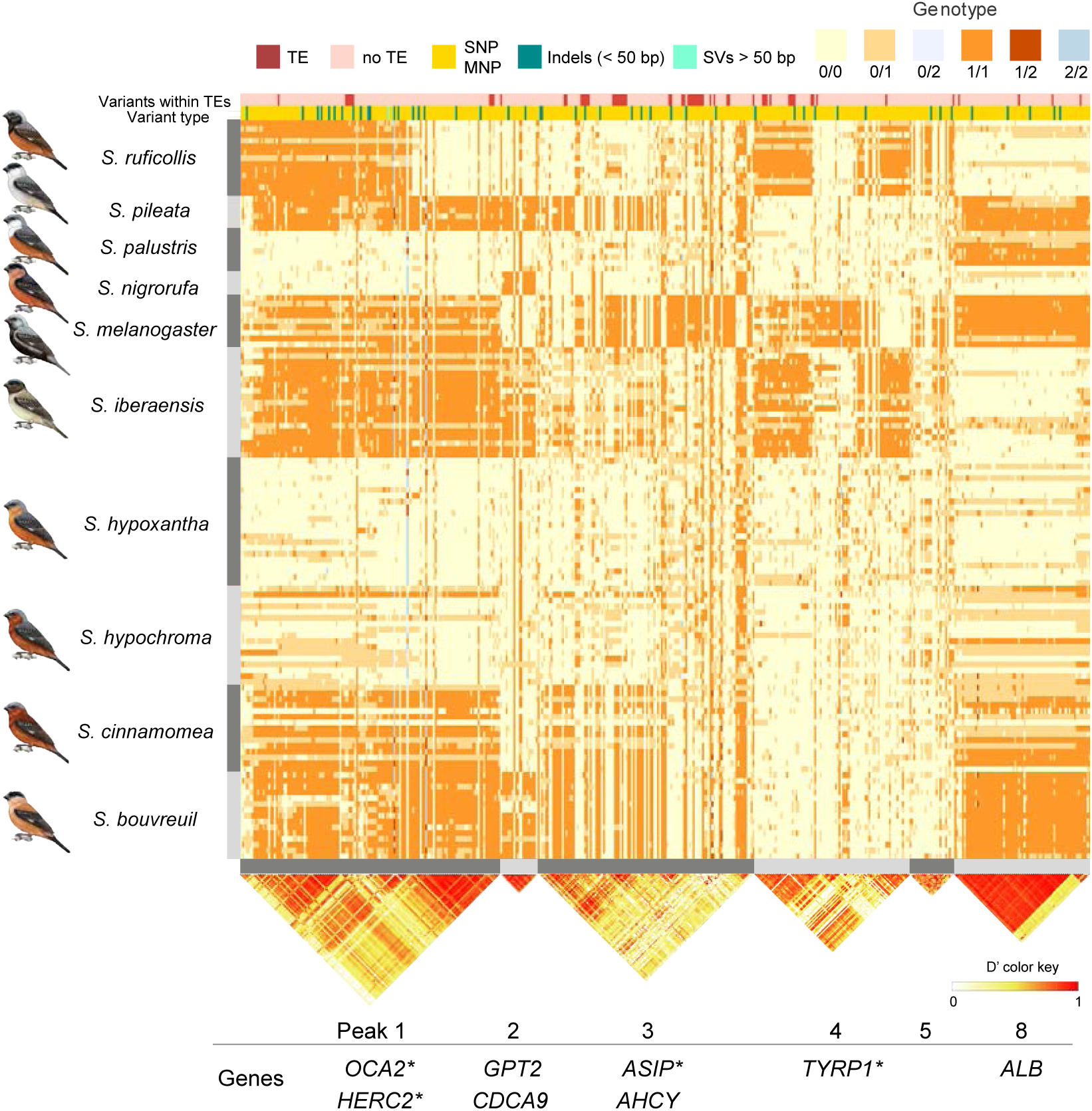
Capuchinos show species-specific patterns of genetic variation when comparing across all divergence peaks. Genotypes for 456 variants with no missing data (for visual simplicity) within peaks 1- 5 and 8 with FST values > 0.75 across populations, including the 55 bp deletion (marked in light blue in the “Variant type” track, see Figure S28C for details), obtained using short-read sequencing data mapped to the pangenome. Peaks 6 and 7 were excluded as they were only detected in the GWAS and do not have variants with FST > 0.75, while peaks 9 and 10 were excluded due to having very few variants, which unnecessarily complicate the plot. The peak IDs correspond to those in Table 1. Rows represent individuals from different populations, while columns correspond to genomic sites, grouped by peak. Variants overlapping transposable elements (TEs) are indicated at the top of the plot, with red denoting sites within TEs and white indicating sites outside of TEs. Variant types are also indicated, with SNPs and multi-nucleotide polymorphisms (MNPs) in yellow, indels (< 50 bp) in teal, and the 55 bp deletion (light blue). The genotypes are color-coded: the homozygous reference (0/0) is represented in yellow, the heterozygous genotypes (0/1, 0/2, 1/2) are represented in light orange, light blue, and dark red, respectively; and the homozygous alternate genotypes (1/1, 2/2) are represented in orange and blue, respectively. The presence of multiple alternate genotypes arises from indels and MNPs, where more than one alternative allele is present. Below each peak, the linkage disequilibrium (LD) pattern is displayed based on the D’ method, color-coded from yellow to red (0 to 1), with red indicating strong LD. Overall, most peaks exhibit high LD, suggesting that the variants within them are inherited in blocks. However, some peaks (e.g., 1, 3, and 4) show distinct LD blocks, indicating potential recombination events. Additionally, the genes within each peak are listed, with those marked by an asterisk (*) indicating genes that are part of the melanogenesis pathway.

The genotypes of variants (SNPs, indels and the 55 bp deletion) with FST > 0.75 within the peaks show clear genetic differentiation among species, with distinct clusters reflecting species-specific allele combinations in these highly differentiated regions (Figure 5). Among these regions, the two comparisons without significant peaks—*S. cinnamomea* vs. *S. hypochroma* and *S. hypoxantha* vs. *S. hypochroma*—emerge as the least differentiated species overall. However, some comparatively more subtle differences are still present in certain regions, such as peaks 1, 3 and 8, where *S. hypoxantha*, and *S. cinnamomea* predominantly exhibit (0/0) and (1/1) genotypes, respectively. Additionally, each comparison includes other differentiated regions that do not meet our peak thresholds. For example, the peak on scaffold 7, which contains the *SLC45A2* gene, includes 19 and 23 variants with FST > 0.5 in the *S. hypoxantha* vs. *S. hypochroma* and *S. hypochroma* vs. *S. cinnamomea* comparisons, respectively.

## Discussion

Our study provides a comprehensive view into the genetic changes driving rapid speciation in Capuchino Seedeaters by leveraging a pangenome built from 32 *de novo* genome assemblies. By integrating this genomic resource with short-read whole-genome sequencing data from ten species (including three previously underrepresented species: Copper Seedeater (*Sporophila bouvreuil*), *S. cinnamomea*, and *S*. *hypochroma*), we refined the characterization of genetic variants driving species differentiation, expanding the analysis beyond SNPs to include small insertions/deletions and other structural variants that have been omitted in past research. Previous studies on Capuchino Seedeaters have proposed that the reshuffling of small regulatory alleles has led to rapid plumage evolution, which coupled with differentiation in male song, contributes to prezygotic isolation and initiating speciation (*19*). These insights were derived from genomic studies based on SNP markers and a single (*S. hypoxantha*) reference genome. An alternative hypothesis to explain rapid speciation in the Capuchinos proposes the existence of species-specific structural variants near these outlier genes. These variants, potentially linked to TE activity, may drive differentiation but could have been overlooked due to methodological limitations. In this study we leverage our pangenome to distinguish these hypotheses.

The high-quality genome assemblies allowed us to improve on previous studies in various ways. First, by combining several genomes, we were able to generate a higher quality reference containing more genomic regions (approximately 1.5 Gb vs. 1.17 Gb, (*17*)), including areas that were not recovered (or are not present) in every individual. Additionally, the use of multiple genomes collectively captured a larger number of genes (15,788 vs. 14,667, (*17*)). We find that our genome assemblies are similar among the different species, and that it remains challenging to distinguish between sequences or genes that are truly missing from an individual from those that are absent due to limitations in the genome assembly process. We therefore recommend using various approaches to, for example, assess the presence and absence of genes. The pangenome shows SNPs are nearly 8 times more prevalent than SVs, with most SVs being shorter than 50 bp (indels), as seen in other systems (e.g.(*27*)). Insertions and deletions are the predominant variant type, while inversions and duplications were rare. SVs were most commonly found in a single or a few individuals at low frequencies. In comparison, two studies, one in the House Finch and one on *Aphelocoma* jays, detected between three and four times more structural variants larger than 50 bp than observed in our system (*28*, *29*). This difference may be due to the more recent diversification, and therefore higher overall genomic similarity in the Capuchinos (*17*, *19*). For instance, a large inversion detected in the House Finch is shared among all three species within the genus *Haemorhous* and is estimated to date back 10 million years (*28*), which represents a significantly longer timeframe compared to Capuchino divergence (*30*). The improved assembly of more complex or repetitive areas allowed us to annotate transposable elements, which show a comparatively minor contribution to patterns of species differentiation. We found that approximately 16% of the genome is composed of repetitive elements, with retrotransposons being most common. Although this TE content is slightly higher than previously reported for most birds (4.1% to 9.8% (*31*), with woodpeckers being an exception and reaching up to 31% (*32*)), TE detection nearly doubled in sparrows when using long-read versus short-read data (*33*). Overall, the pangenomic approach allowed us to recover regions of the genome (and variant types) that had been overlooked by previous studies in our system, and this methodological advance will likely also benefit other systems in the future.

The combination of the pangenome with short-read data failed to recover ∼29.6 million variants present in the pangenome. This reduction is due to the inherent limitations of short-read data in properly mapping to regions containing larger SVs and repetitive sequences (*34*), resulting in the loss of many variants at the scaffold ends and in other similarly highly repetitive regions.

Therefore, we may still be missing important differences in such regions, which could be relevant to species differentiation. While we now have the pangenome as a reference containing those regions, genotyping a larger number of individuals with long-read data will be crucial for resolving species differences in those challenging areas. Additionally, here we rely on variant call files (VCFs) to represent SVs, yet capturing the complexity of SVs in this format remains challenging (*29*). Our FST and GWAS analyses treat these SVs as multi-allelic loci and may therefore overlook potential relationships among alleles within a locus.

Despite variation in the numbers of individuals and species studied, the GWAS analyses for pigment concentrations and the FST scans consistently revealed the strongest outlier regions containing melanogenesis genes (*HERC2*/*OCA2*, *ASIP*, *TYRP1*, and *SLC45A2*), corroborating previous findings (*17*, *19*, *20*). The current study also confirmed patterns that had been previously identified in the Capuchino radiation, like the prevalence of differences in non-coding variants and the enrichment of outlier peaks on the Z sex chromosome. As seen before (*19*), species show unique combinations of genotypes across the different main outlier peaks, although other regions dispersed across the genome with more subtle patterns of differentiation also exist. Moreover, we find that differentiation levels vary among species, suggesting that the amount of gene flow and/or the number and extent of the genomic regions necessary to generate phenotypic differences varies depending on the species. The differentiated regions of the genome show a prevalence of SNPs over small SVs and an absence of large SVs (with a single exception near the *HERC2* and *OCA2* genes). The outlier regions did not differ significantly from the rest of the genome in terms of their SV or TE content. The different types of variants within outlier peaks (SNPs, indels) are generally found in high LD, likely acting collectively to shape phenotypes.

Future integration of chromatin accessibility and interaction data (such as ATAC-seq and Hi-C) with genomic variant analysis is needed to uncover regulatory networks and understand the functional significance of these non-coding regions, where over 90% of enhancers—key regulators of gene expression—are located (*35*). Importantly, by excluding species-specific SVs in outlier regions, we conclude that the reshuffling of regulatory alleles remains the most likely mechanism driving the rapid speciation of the Capuchinos.

While structural variants do not seem to have strongly shaped Capuchino Seedeater evolution, our results highlight the power of the pangenome framework for studying phenotypic evolution and speciation. SVs may play a more prominent role in systems with longer divergence times, but their full extent may currently be overlooked due to methodological limitations. Pangenomes are beginning to emerge in a wide range of organisms beyond plants and bacteria, as advances in sequencing technologies make these comprehensive genomic studies more feasible. These developments provide the opportunity to integrate all forms of genetic variation into the search for the genetic basis of phenotypes in non-model organisms (e.g. (*36–38*)), and promise to uncover previously hidden genetic contributions to phenotypic traits and adaptive evolution.

## Materials and Methods

### Sampling and sequencing

We generated a pangenome by selecting 16 individuals including between one and four individuals from seven highly sympatric species of Capuchino Seedeaters: *S. palustris* (3), *S. ruficollis* (2), *S. pileata* (1), *S. iberaensis* (4), *S. cinnamomea* (2), *S. hypoxantha* (3), and *S. hypochroma* (1). All individuals were males except for one *S. iberaensis and* one *S. hypoxantha* (Table S12). We extracted high molecular DNA from blood samples using the Zymo Research Quick-DNA HMW MagBead Kit, and one sample originated from a phenol/chloroform extraction of muscle tissue. We assessed the concentration of recovered DNA using the Qubit Broad Range Assay Kit (Thermo Fisher Scientific, Waltham, MA) and analyzed the size of the fragments with the Femto Pulse QC system (Cornell Biotechnology Resource Center Genomics Core). We used between 10 to 25 μg of high molecular weight DNA for Pacific Biosciences (PacBio) HiFi library preparation, which was conducted by the sequencing facilities. We subsequently sequenced all individuals using one PacBio HiFi Revio SMRT Cell per individual at the Novogene (Sacramento, CA) and Cornell Weill (New York, NY) sequencing centers. For the 16 Capuchino individuals sequenced using PacBio HiFi, the average sequencing yield was 72 Gb per sample (range 36-89 Gb) with a mean read length of 15,596 bp (range of 11,376 to 19,637 bp; Table S13).

Additionally, we used previous whole-genome resequencing (WGS) data from 121 individuals of 10 species (range between 3 and 28 individual per species; see Table S13 for details including Accession numbers), and generated new data for 41 additional individuals including one that was previously sequenced (for details see Table S14). The new sequences were obtained after extracting DNA from blood stored in ethanol, using the Qiagen DNeasy Blood and Tissue kit.

We prepared libraries using the New England BioLabs NENNext Ultra II FS DNA Library Prep Kit for Illumina with TrueSeq adapters. The sequencing was performed in two batches using an Illumina NovaSeq X – 2 x150 paired end lane from Novogene and one from the Biotechnology Resource Center (BRC) at the Cornell Genomics Facility (Table S14). The two sequencing batches yielded an average of 90.4 and 214.6 million raw reads, respectively (Table S14).

### Genome assemblies and annotations

We used the PacBio HiFi reads to generate *de novo* assemblies for each individual. First, we verified the absence of adapters and converted BAM files to FASTQ format using HiFiAdapterFilt v3.0.0 (*39*). Next, we produced primary and alternate assemblies (haplotype-resolved) with hifiasm v0.19.9 (*40*), followed by the removal of haplotigs and contig overlaps using purge_dups v1.2.6 (*41*). Genome size and heterozygosity were estimated using Jellyfish v2.3.0 (*42*) and GenomeScope v2.0 (*43*), employing a k-mer size of 21, and adjusting to 19 when the model failed to converge. GenomeScope predicted a similar heterozygosity and genome length across all assemblies, with an estimated heterozygosity of ∼1.2% and an initial estimated genome size of ∼0.99 Gb (Table S15, Figure S29). For the purged assemblies of the primary and alternate haplotypes, we assessed assembly metrics using assembly-stats v.1.0.1 (https://github.com/sanger-pathogens/assembly-stats) (Table S16), and Merqury plots and QV scores (Quality Value) obtained with Merqury v1.3. The QV score is a measure of accuracy in logarithmic scale, representing the number of errors per base in the assembly; higher QV scores indicate fewer errors and better assembly quality. A QV of 60 corresponds to an estimated error rate of 1 in 1 million bases. The Merqury plots (Figure S30) and QV scores (Table S16) further confirmed the high quality of the assemblies. Specifically, the mean QV scores (± standard deviation) were 61.4 (±1.2) and 63.2 (±1.0) for the primaries and alternate assemblies, respectively (Table S16). The quality and completeness of the assemblies was further evaluated with the Benchmarking Universal Single-Copy Orthologs (BUSCO v5.5.0) pipeline (*44*) using the Aves database (aves_odb10; Table S3).

We used the genome HYPOXB009684 as the reference for subsequent analyses requiring a single reference and for anchoring the pangenome to a coordinate system. We selected this genome because it belongs to the same species (*S. hypoxantha*) as the original reference genome (Genbank Assembly GCA_002167245.1) described in Campagna et al. (*17*), and because it has slightly higher contiguity among the three available assemblies from this species.

We identified and masked repetitive elements prior to genome annotation and pangenome construction. First, we built a custom repeat library for the Capuchino Seedeaters using RepeatModeler v2.0.1 (*45*) with default options. To complement this custom library and improve the classification of LINE and SINE elements, avian repeat families were retrieved from the Dfam 3.8 database (*46*) and merged with the custom Capuchino repeat library that we generated with RepeatModeler. Finally, the combined repeat library was applied to soft-mask repetitive regions of each Capuchino genome assemblies using RepeatMasker v4.0.7 with the RMBlast engine (*47*).

We carried out gene prediction using BRAKER3 (*48*), which combines protein evidence with *ab initio* prediction models. First, a comprehensive protein database was constructed by merging the OrthoDB vertebrate dataset with the proteome of the Zebra finch (*Taeniopygia guttata*), downloaded from UniProt ((*49*), Zebra finch, UniProt proteome UP000007754). The protein sequences were reformatted to ensure consistency across datasets, and the combined database was compressed for use. The masked genomes, generated using RepeatModeler, served as the input for BRAKER3 alongside the combined vertebrate and Zebra finch protein dataset. We executed BRAKER3 specifying the AUGUSTUS *ab initio* model, protein alignment via ProtHint, and diamond for sequence similarity search, resulting in gene predictions output in GFF3 format. Additionally, for a homology-based gene annotation approach, we downloaded annotations for both the chicken (*Gallus gallus*; GCF_000002315.6_GRCg6a) and the Zebra finch (GCF_003957565.2_bTaeGut1.4). First, the annotations were reformatted using AGAT v1.2.0 (*50*) to ensure compatibility, and the new GFF files were then used in the GeMoMa v1.9 (*51*) pipeline for further refinement of gene models. Using the chicken and Zebra finch reference genomes, GeMoMa initially predicted around 51,380 ± 2,078 (SD) genes. Finally, we used GeMoMa’s GAF tool to combine the genes predicted from both reference genome annotations (i.e. chicken and Zebra finch), applying a filter criterion that required the start codon to be ’M’, the stop codon to be ’*’, and the score per amino acid to be greater than or equal to 0.75. After combining the predicted genes from both annotations and filtering, this resulted in an average of 14,666 genes per annotation (Table S2).

### Synteny among assemblies and Gene Presence Absence Variation (PAV) analysis

We used the GENESPACE v1.3.1 (*52*) R package in R version 4.2.3 (*53*) with default parameters to infer and visualize synteny blocks among the thirty longest scaffolds from all the primary assemblies. GENESPACE combines OrthoFinder and MCScanX to identify orthologous and paralogous genes across multiple genomes and detect syntenic blocks to identify collinear regions among genomes.

We combined two complementary approaches to assess gene-level variation among assemblies. First, we extracted gene lists from each GFF file and compared gene presence and absence across individuals, identifying 13,573 common genes among all primary assemblies. In parallel, we used Pangene v1.1 (*54*), which leverages miniprot to align protein-coding exons from all genes, extracted from the primary assembly annotations (GFF files), to all the remaining assemblies.

We ran Pangene with default parameters, setting the “outs” option to 0.99 to ensure that only the best alignments with a score of at least 99% for the protein were included. This process generated a pangenome graph in GFA format and identified 11,760 genes common among all primary assemblies. Both approaches provided lists of genes unique to each individual, which we then compared to identify overlaps and find the most robust presence-absence variation (PAV) candidates which were consistent across both methods. We compared the gene lists comparisons in R and represented the results using Venn Diagrams and Upset plots using the R libraries ggVennDiagram v1.5.2 (*55*) and UpSetR v1.4.0 (*56*). We combined the individual lists of genes per species for the Venn Diagram plots. Both methods found a similar number of unique genes per species, except for *S. palustris*, where the Pangene method identified 3.5 times more unique genes (Figure S1, S2). However, the identity of these genes varied between methods, indicating that each method missed some genes recovered by the alternative approach. This underscores the challenges of mapping protein sequences to genomes and accurately annotating all genes across our datasets. The overlapping genes between both methods, which we considered strong PAV candidates, were retrieved from the specific annotations and then aligned using BLAST v2.16.0 (*57*) against all genomes to confirm their presence or absence. We focused exclusively on the primary assemblies, as alternate assemblies tend to be more fragmented and prone to missing genes that may still be present in the primary assemblies. Only 13 genes (Table S5) were detected to be uniquely present in certain species by both methods, and after aligning these using BLAST against all genomes, we were able to consistently recover them (Table S5). This suggests that differences in thresholds and alignment methods can result in some genes being overlooked by specific approaches. Therefore, we do not have strong evidence for genes that are present or absent in certain species, although not all genes are represented in every individual annotation.

### Identifying and genotyping SVs using long read data

To characterize SVs longer than 50 bp with our PacBio HiFi reads, we employed three SV calling methods: PBSV v2.6.2 (*58*) Sniffles v2.2 (*59*) and SVIM-asm v.1.0.3 (*60*). For PBSV, we first aligned each sample’s reads to the HYPOXB009684 reference genome using pbmm2 v17.0 align with the settings “sort” and “--preset-ccs”. Signatures of structural variation were identified with pbsv discover, incorporating a BED file of repetitive regions via the “--tandem-repeats” option to improve SV detection. Finally, we detected SVs per individual using pbsv call with default settings, specifying “--ccs” and “--preserve-non-acgt”. For Sniffles, we mapped the PacBio HiFi reads from each individual to the reference genome (HYPOXB009684) using minimap2 v2.28, then called SVs per sample using default parameters. For SVIM-asm, we aligned the primary and alternate assemblies of each sample to the HYPOXB009684 reference genome with minimap2, using the options “-a -x asm5 --cs -r2k”. Variants were then called for each sample with svim-asm diploid and default parameters. We sorted and indexed all resulting variant call files (VCF) using BCFtools v1.20 (*61*). PBSV detected the highest number (∼244K) of SVs, which is around 2.2 and 2.5 times more than Sniffles2 and SVIM-asm, respectively (Table S17). To consolidate results, we used SURVIVOR v1.0.7 merge (*62*) to combine the three VCFs per sample generated by the different SV callers, retaining only calls detected by all three methods and within a 1 kb distance of each other as the same event (using the options 1000 3 1 1 0 1). When two homologous variants have slight sequence differences, breakpoint-based approaches like SURVIVOR can be too conservative (*63*). The 1 kb threshold to determine the coincidence of SVs from the three different calling methods has been used in other SV analyses (e.g. (*64*, *65*)), and using a smaller threshold, such as 500 bp, resulted in minimal changes to the number of recovered variants. SURVIVOR identified a similar number of SVs for each sample supported by the three SV callers (∼55K) (Table S17). We next combined the SVs retained by SURVIVOR for each individual into a single VCF file, retaining all the calls longer than 50 bp, merging those within a 1 kb distance and not taking into account the DNA strand of the SV (using the options 1000 1 1 0 0 50). We extracted the SV type and length and classified them in length intervals (50-200 bp, 200-500 bp, 500-1 kb, 1-5 kb, 5-10 kb and 10-15.5 Mb).

### Pangenome graph construction and variant decomposition

We used the Cactus Pangenome pipeline v2.8.0 (*66*) to generate the Capuchino pangenome. Briefly, minigraph v0.20 is used to generate a Graphical Fragment Assembly (GFA) graph from the purged and masked HiFi primary and alternate assemblies for the 16 individuals (i.e., 32 haplotypes). Then the assemblies are mapped back to the graph using minigraph. These mapped assemblies are the input for Cactus to generate a new graph that includes variants of all sizes obtaining the pangenome graph as output in different formats including giraffe, gfa, gbz and vg. We calculated statistics from the pangenome graph in GFA format using Panacus v.0.2.3 (*67*), including pangenome length, as well as the core and accessory genome lengths. The core genome was defined as the regions shared by all the haplotypes. Additionally, we used Panacus to describe the pangenome features in terms of the number of nodes and edges that are expected on average when the haplotypes are added sequentially. We annotated the variant types from the VCF file obtained from the pangenome using vcf-annotate “--fill-type” from the VCFtools v0.1.16 library (*68*). We classified variants as SNPs and multi-nucleotide polymorphisms (MNPs) if all alternate alleles were either SNPs, MNPs, or a combination of both. Insertions and deletions were identified when all alternate alleles supported those variant types, while variants were categorized as complex when alternate alleles were a mix of variant types or specifically marked as complex by vcf-annotate.

### Mapping short-read data to the pangenome

We used the vg toolkit v1.53.0 (*69*) for pangenome-based variant calling and genotyping. We mapped short-read data from 161 individuals using vg giraffe (*70*). The whole genome re- sequencing data was first processed to remove adapters and filtered using AdapterRemoval v2.1.1 (*71*). We set the minimum length to 75 bp, the minimum Phred quality to 10 and merged overlapping paired-end reads. All the output files (e.g., truncated or merged reads) were mapped separately against the pangenome graph and the resulting GAM files were then merged for every individual. Using vg pack, we computed the read support from the GAM files, ignoring sites with mapping and base quality below 5 (“-Q 5”). With the pack files, a VCF file was generated per individual using vg call (*72*). Because some of the data from previous studies had lower coverage, we retained only individuals with an average depth of coverage greater than 4X resulting in a final dataset with 127 individuals (Table S18). Finally, the VCF files were indexed and combined into a single VCF using BCFtools v1.20 (*61*) index and merge, respectively. This VCF file was filtered to retain sites with a mean depth between 4X and 50X, missing data below 0.8, and any locus with a non-reference allele count (not necessarily the same allele) of at least 4 (“--non-ref-ac-any 4”). The resulting VCF file was divided to generate five datasets: 1) SNPs and MNPs (referred to as the SNP dataset); 2) short SVs (SVs < 50 bp) also referred to as indels; 3) SVs longer than 50 bp; 4) SNPs and SVs combined, within annotated repetitive elements and TEs (referred to as variants within TEs); and 5) SNPs and SVs outside annotated repetitive elements and TEs (variants outside TEs). Then the SNP dataset was furthered filtered by minor allele count 4 (“--mac 4”). For the GWAS analysis we generated a dataset with all SVs and colored the resulting plots according to their length (greater or smaller than 50 bp). All the mapping steps were performed using GNU Parallel v20170522 (*73*). We calculated the length of a variant as the average length of all the alternate alleles at that site. We calculated the allele frequencies for the SNP and SV datasets (combining both indels and long SVs) and the allele count for the SV dataset using VCFtools (*68*) and plotted the histogram of the alternative allele. For SVs with more than one alternative allele, we tested both calculating the average frequency of all alternative alleles and using only the first alternative allele, and the distributions were nearly identical. We show the results from the first approach. We also visualized the relationship between SV length and both allele frequency and allele count using 2D-hexbin density plots in ggplot2 v3.5.1 (*74*). We log-transformed SV lengths, applied hexagonal binning with 100 bins, and used color intensity to reflect data density.

Due to the lower number of variants recovered after mapping short-read data against the pangenome compared to the variants present in the pangenome, we studied the patterns of mapping quality. To calculate the mean mapping quality of the short-read data aligned to the pangenome, we first used the vg surject command from the vg toolkit to generate a BAM file from the pangenome graph and the GAM alignments from five individuals (see Table S18 for details). We then used SAMtools v1.20 (*61*)to extract the mapping quality values per site from the resulting BAM file. To assess whether mapping quality differs between regions with low and normal mean depth of coverage, the averaged and binned data (of 5 individuals in 50 kb windows) were divided into two groups based on the average coverage: one group with coverage less than 1 and the other with coverage greater than or equal to 1 (≥1). Given the imbalance in sample sizes between these groups, we performed undersampling by randomly selecting a subset of observations from the larger group (≥1) to match the sample size of the smaller group (<1).

The mapping quality values in both groups did not follow a normal distribution according to a Shapiro-Wilk test. Therefore, we applied the non-parametric Wilcoxon rank-sum test to compare the distribution of mapping quality between the two groups.

We used the dataset derived from mapping short-read data to the pangenome to scan the genome for large inversions using the local PCA method implemented in the lostruct package in R (*75*). This method detects outlier regions of the genome by computing PCAs for SNP windows and calculates the distance among all windows. Finally, multidimensional scaling (MDS) is used to display the distance among all windows and identify those with anomalous patterns. We applied the method to scaffolds larger than 1 Mb in window sizes of 1,000 SNPs, and used the R package SNPRelate v1.36.1 (*76*) to compute and display PCAs from the scaffolds of interest.

### GWAS and FST scans using SNPs and SVs

We conducted genome-wide association studies (GWAS) with four datasets (SVs grouped together irrespective of size), including the 127 individuals from 10 species (see Table S18 for details) using PLINK v2 (*77*). *Sporophila minuta* and *S. castaneiventris* were not included in this study as they show higher genetic differentiation from the remaining 10 species, known as the southern Capuchino Seedeaters (*15*, *16*). First, we filtered out SVs with more than 254 alleles in all datasets, resulting from multiple alternate alleles, as PLINK cannot process these (excluding 3,444 sites for the SVs dataset, and 1,103 and 2,341 for the datasets including SNPs and SVs within and outside TEs, respectively) (Table S19). We used the mean pigment concentrations (eumelanin and pheomelanin) per species across six plumage patches as our phenotype, as described in (*20*), leading to 12 different GWAS. The first 10 principal components from a Principal Components Analysis were used to control for population structure, which in the Capuchinos is low (*17*). Moreover, to control for multiple comparisons, we applied the Bonferroni correction by adjusting the significance threshold based on the total number of observations (SNPs+SVs) and multiplying by the number of GWAS (12), resulting in p ≤ 2.65 x10⁻⁹ as a cutoff for statistical significance. We combined outliers into peaks when there were more than 5 hits close to each other (< 50 kb), while isolated outliers are listed in Supplementary Materials (Table S7).

We ran FST scans with VCFtools (*68*) on all five original datasets per site and in 10 kb windows (Table S19). For the FST comparisons we only included six out of the seven species with more than eight individuals in our dataset (*S. cinnamomea*, *S. iberaensis*, *S. hypoxantha*, *S. hypochroma*, *S. melanogaster and S. ruficollis*), excluding *S. bouvreuil* because it shows an overall higher level of differentiation (*30*), resulting in a total of 95 individuals across 15 pairwise comparisons. Excluding *S. bouvreuil* also allowed us to both reduce the number of pairwise comparisons and focus on the species with the lowest background level of differentiation, making the FST peak analyses more manageable. Because of the large number of comparisons, we decided to define FST outlier peaks in a way that focuses on the most highly differentiated genomic regions between species. We defined outlier windows as those with weighted FST in the top 0.1% of the FST distribution and containing at least one individual variant with FST > 0.75. We combined consecutive outlier windows into peaks, and report the remaining outlier windows in Supplementary Materials (Table S9). Other regions of the genome may show more subtle patterns that are still relevant to shaping phenotypic differences.

Due to PLINK’s limitation of performing GWAS with sites containing more than 254 alternate alleles, 3,444 SVs were excluded, 99.7% of which were long SVs (Table S19). However, these variants were included in the FST scans, and none showed an FST > 0.75. Therefore, although we are not directly testing their association with pigment concentrations, assuming these regions are being correctly assembled, it is complicated to assess if they are playing a significant role in species differentiation. Once we narrowed down the main outlier peaks, we compared their TE and SV content per kb to the average genomic distribution. We performed a permutation test to assess statistical significance by sampling 1 kb windows in the 30 longest scaffolds, excluding the first and last 50 kb to create genomic TE and SV distributions. By comparing the observed values in the peaks to this distribution, we could determine whether the observed TE or SV content per kb differed from what would be expected by chance from the genome. The observed values did not fall within the bottom or top 2.5% of the null distribution, and we thus concluded that the SV and TE content within the peaks did not differ significantly from the genome-wide distribution. Additionally, to visualize patterns of linkage disequilibrium (LD) within the peaks, we used LDBlockShow v1.40, (*78*) to generate LD plots based on the D’ method from the variants within each peak with FST ≥0.75 and without any missing data.

## Acknowledgments

We thank the members of the Fuller Evolutionary Biology laboratory group (particularly B. Butcher and Irby Lovette) for lab and computational support. We thank the volunteers for essential help with fieldwork. We also thank the government agencies of Brazil (ICMBio, MMA, and SISBIO) and Argentina (SAyDS, Dirección de Recursos Naturales Corrientes, Dirección de Parques y Reservas de la Provincia de Corrientes, Administración de Parques Nacionales) for providing the necessary research authorizations and collection permits that facilitated our work (63556-12, Brazil, and IF-2018-53280705-APN-DRNEA#APNAC, IF-2022-133544133-APN-DRNEA#APNAC, Argentina). Samples for this study were exported through permits CE-2023- 49830382-APN-DNBI#MAD, and CE-2024-83310820-APN-SSAM#JGM.

## Author contributions

Conceptualization: LC, MR Methodology: MR, LC

Investigation: MR, LC, SK, JRRR, MR, MB, JFC, ASDG, CK.

Visualization: MR Writing—original draft: MR, LC Writing—review & editing: MR, LC

## Data and materials availability

All data needed to evaluate the conclusions in the paper are present in the paper and/or the Supplementary Materials. Additional data related to this paper may be requested from the authors. Genomic data have been archived in GenBank (BioProject ID PRJNA382416). The genome assemblies, generated from PacBio HiFi data, are available under BioProjects PRJNA1223491–PRJNA1223508 and PRJNA1223510–PRJNA1223523. All the accession numbers are provided in the Supplementary Materials.

