## Supplementary Material for "A pangenomic approach reveals the sources of genetic variation fueling the rapid radiation of Capuchino Seedeaters"

### Supplementary Tables

**Table S1. Transposable element composition for each assembly.** All primary assemblies, irrespective of species, have approximately 16% of the genome covered by TEs, with retroelements being the most common element type. The table includes GC content (%), percentages of retroelements (SINEs, LINEs, and LTRs), DNA transposons, rolling circles, unclassified elements (Uncl.), small RNA, satellites, simple repeats, low complexity repeats, and the total percentage of the genome covered by transposable elements (TEs) irrespective of class. It also provides the mean and standard deviation across all assemblies for each TE category.

| Sample | Species | Sex | %GC | Retro. Total | Retro. SINEs | Retro. LINEs | Retro. LTR | DNA Transp. | Rolling-circles | Uncl. | Small RNA | Satellites | Simple repeats | Low Complex. | Total % genome |
|---|---|---|---|---|---|---|---|---|---|---|---|---|---|---|---|
| CIN3121 | *S. cinnamomea* | Male | 42.9 | 11.0 | 0.11 | 5.4 | 5.5 | 0.24 | 0.05 | 3.1 | 0.04 | 0.38 | 1.3 | 0.25 | 16.4 |
| CIN3122 | *S. cinnamomea* | Male | 42.9 | 11.2 | 0.11 | 5.4 | 5.6 | 0.24 | 0.06 | 3.1 | 0.05 | 0.43 | 1.2 | 0.26 | 16.6 |
| CRO3131 | *S. hypochroma* | Male | 43.0 | 11.4 | 0.11 | 5.5 | 5.8 | 0.23 | 0.06 | 3.4 | 0.04 | 0.49 | 1.3 | 0.25 | 17.1 |
| HYPOXB009213 | *S. hypoxantha* | Male | 42.8 | 10.5 | 0.11 | 5.2 | 5.2 | 0.24 | 0.07 | 3.0 | 0.04 | 0.31 | 1.3 | 0.25 | 15.7 |
| HYPOXB009666 | *S. hypoxantha* | Female | 43.0 | 11.2 | 0.11 | 5.3 | 5.9 | 0.24 | 0.07 | 3.2 | 0.05 | 0.39 | 1.2 | 0.25 | 16.7 |
| HYPOXB009684 | *S. hypoxantha* | Male | 42.9 | 10.9 | 0.11 | 5.3 | 5.5 | 0.24 | 0.07 | 3.2 | 0.05 | 0.40 | 1.2 | 0.25 | 16.3 |
| IBEB009240 | *S. iberaensis* | Male | 43.0 | 11.2 | 0.11 | 5.5 | 5.7 | 0.23 | 0.05 | 3.2 | 0.04 | 0.40 | 1.3 | 0.25 | 16.6 |
| IBEB009632 | *S. iberaensis* | Female | 42.8 | 10.4 | 0.11 | 5.2 | 5.0 | 0.24 | 0.07 | 2.8 | 0.04 | 0.34 | 1.2 | 0.25 | 15.3 |
| IBEB009696 | *S. iberaensis* | Male | 43.0 | 11.2 | 0.11 | 5.5 | 5.6 | 0.24 | 0.08 | 3.2 | 0.05 | 0.45 | 1.2 | 0.25 | 16.7 |
| IBEB009699 | *S. iberaensis* | Male | 43.0 | 11.1 | 0.11 | 5.4 | 5.6 | 0.24 | 0.07 | 3.3 | 0.04 | 0.42 | 1.3 | 0.25 | 16.7 |
| PAL3117 | *S. palustris* | Male | 42.9 | 10.8 | 0.11 | 5.4 | 5.4 | 0.24 | 0.05 | 3.0 | 0.05 | 0.40 | 1.2 | 0.25 | 16.1 |
| PAL3118 | *S. palustris* | Male | 42.8 | 10.4 | 0.11 | 5.2 | 5.1 | 0.24 | 0.04 | 2.9 | 0.05 | 0.39 | 1.2 | 0.25 | 15.5 |
| PAL3372 | *S. palustris* | Male | 42.8 | 10.3 | 0.11 | 5.1 | 5.1 | 0.24 | 0.05 | 2.9 | 0.04 | 0.37 | 1.2 | 0.25 | 15.4 |
| PIL3664 | *S. pileata* | Male | 42.8 | 10.3 | 0.11 | 5.2 | 5.0 | 0.24 | 0.05 | 2.8 | 0.04 | 0.34 | 1.2 | 0.25 | 15.2 |
| RUF3128 | *S. ruficollis* | Male | 42.9 | 10.5 | 0.11 | 5.2 | 5.2 | 0.24 | 0.04 | 2.9 | 0.05 | 0.35 | 1.2 | 0.25 | 15.6 |
| RUF3129 | *S. ruficollis* | Male | 42.9 | 11.1 | 0.11 | 5.4 | 5.6 | 0.24 | 0.06 | 3.1 | 0.04 | 0.43 | 1.3 | 0.25 | 16.5 |
| MEAN | | | 42.9 | 10.8 | 0.11 | 5.3 | 5.4 | 0.24 | 0.06 | 3.1 | 0.04 | 0.39 | 1.2 | 0.25 | 16.2 |
| SD | | | 0.07 | 0.38 | 0.00 | 0.12 | 0.28 | 0.00 | 0.01 | 0.18 | 0.01 | 0.05 | 0.02 | 0.00 | 0.60 |

**Table S2. Gene annotation for each reference assembly.** Number of genes recovered by GeMoMa before and after combining and purging the genes annotated using the chicken and Zebra finch annotations. We used GeMoMa's GAF tool to combine and purge the genes from both annotations, applying filtering criteria that required the start codon to be 'M', the stop codon to be '*', and the score per amino acid to be greater than or equal to 0.75. The initial annotations contained on average 51,380 genes, which after filtering resulted in around 14,666. Female individuals are marked with an asterisk (*) in the species column. The table shows that the number of annotated genes across species is fairly consistent, particularly after filtering, suggesting that gene annotations are comparable across species.

| Annotation | Species | GeMoMa initial gene number | Final gene number |
|---|---|---|---|
| CIN3121 | *S. cinnamomea* | 52,025 | 14,665 |
| CIN3122 | *S. cinnamomea* | 53,884 | 14,690 |
| CRO3131 | *S. hypochroma* | 54,676 | 14,689 |
| HYPOXB009213 | *S. hypoxantha* | 49,980 | 14,659 |
| HYPOXB009666 | *S. hypoxantha** | 49,973 | 14,709 |
| HYPOXB009684 | *S. hypoxantha* | 52,235 | 14,720 |
| IBEB009240 | *S. iberaensis* | 53,486 | 14,682 |
| IBEB009632 | *S. iberaensis** | 49,877 | 14,532 |
| IBEB009696 | *S. iberaensis* | 50,554 | 14,707 |
| IBEB009699 | *S. iberaensis* | 54,706 | 14,711 |
| PAL3117 | *S. palustris* | 48,843 | 14,638 |
| PAL3118 | *S. palustris* | 49,501 | 14,665 |
| PAL3372 | *S. palustris* | 51,200 | 14,653 |
| PIL3664 | *S. pileata* | 49,148 | 14,611 |
| RUF3128 | *S. ruficollis* | 48,893 | 14,650 |
| RUF3129 | *S. ruficollis* | 53,104 | 14,669 |
| MEAN | | 51,380 | 14,666 |
| SD | | 2,078 | 46 |

**Table S3. BUSCO scores for the primary and alternate assemblies.** The BUSCO analysis was performed using the aves_db10 avian database, which includes 8,338 single-copy orthologs. The table reports the number and percentage of complete orthologs (C), subdivided into single-copy (S) and duplicated (D) orthologs, as well as those that are fragmented (F) and missing (M). BUSCO scores for both primary and alternate assemblies are consistently high across all species, with the percentage of complete orthologs generally exceeding 96%. The primary assemblies had a smaller proportion of missing and fragmented genes compared to the alternate assemblies, while duplicated genes were consistently around 0.5% across all assemblies. While the primary assemblies typically show slightly better scores than the alternate assemblies, both exhibit comparable quality and completeness in gene recovery and are also similar across species. PRI denotes the primary haplotype and ALT the alternate one.

| | C | C% | S | S% | D | D% | F | F% | M | M% |
|---|---|---|---|---|---|---|---|---|---|---|
| RUF3129_PRI | 8,090 | 97.0% | 8,056 | 96.6% | 34 | 0.4% | 45 | 0.5% | 203 | 2.5% |
| RUF3129_ALT | 8,061 | 96.7% | 8,018 | 96.2% | 43 | 0.4% | 44 | 0.5% | 233 | 2.8% |
| IBEB009240_PRI | 8,083 | 96.9% | 8,049 | 96.5% | 34 | 0.4% | 49 | 0.6% | 206 | 2.5% |
| IBEB009240_ALT | 8,039 | 96.4% | 7,998 | 95.9% | 41 | 0.5% | 52 | 0.6% | 247 | 3.0% |
| PAL3118_PRI | 8,091 | 97.1% | 8,060 | 96.7% | 31 | 0.4% | 44 | 0.5% | 203 | 2.4% |
| PAL3118_ALT | 8,053 | 96.6% | 8,012 | 96.1% | 41 | 0.5% | 47 | 0.6% | 238 | 2.8% |
| PAL3117_PRI | 8,086 | 96.9% | 8,057 | 96.6% | 29 | 0.3% | 47 | 0.6% | 205 | 2.5% |
| PAL3117_ALT | 8,062 | 96.7% | 8,028 | 96.3% | 34 | 0.4% | 51 | 0.6% | 225 | 2.7% |
| PAL3372_PRI | 8,095 | 97.1% | 8,059 | 96.7% | 36 | 0.4% | 49 | 0.6% | 194 | 2.3% |
| PAL3372_ALT | 8,064 | 96.7% | 8,033 | 96.3% | 31 | 0.4% | 48 | 0.6% | 226 | 2.7% |
| RUF3128_PRI | 8,081 | 96.9% | 8,041 | 96.4% | 40 | 0.5% | 46 | 0.6% | 211 | 2.5% |
| RUF3128_ALT | 8,050 | 96.5% | 8,008 | 96.0% | 42 | 0.5% | 60 | 0.7% | 228 | 2.8% |
| HYPOXB009213_PRI | 8,086 | 97.0% | 8,048 | 96.5% | 38 | 0.5% | 47 | 0.6% | 205 | 2.4% |
| HYPOXB009213_ALT | 8,068 | 96.8% | 8,026 | 96.3% | 42 | 0.5% | 52 | 0.6% | 218 | 2.6% |
| IBEB009632_PRI0 | 8,081 | 96.9% | 8,046 | 96.5% | 35 | 0.4% | 54 | 0.6% | 203 | 2.5% |
| IBEB009632_ALT0 | 7,973 | 95.6% | 7,925 | 95.0% | 48 | 0.6% | 61 | 0.7% | 304 | 3.7% |
| HYPOXB009666_PRI | 8,074 | 96.9% | 8,034 | 96.4% | 40 | 0.5% | 49 | 0.6% | 215 | 2.5% |
| HYPOXB009666_ALT | 7,669 | 91.9% | 7,632 | 91.5% | 37 | 0.4% | 58 | 0.7% | 611 | 7.4% |
| HYPOXB009684_PRI | 8,079 | 96.9% | 8,043 | 96.5% | 36 | 0.4% | 51 | 0.6% | 208 | 2.5% |
| HYPOXB009684_ALT | 8,060 | 96.6% | 8,016 | 96.1% | 44 | 0.5% | 42 | 0.5% | 236 | 2.9% |
| IBEB009696_PRI | 8,097 | 96.7% | 8,062 | 96.7% | 36 | 0.4% | 49 | 0.6% | 192 | 2.3% |
| IBEB009696_ALT | 8,037 | 96.4% | 8,001 | 96.0% | 36 | 0.4% | 54 | 0.6% | 247 | 3.0% |
| IBEB009699_PRI | 8,083 | 96.9% | 8,048 | 96.5% | 35 | 0.4% | 50 | 0.6% | 205 | 2.3% |
| IBEB009699_ALT | 8,006 | 96.1% | 7,960 | 95.5% | 46 | 0.6% | 62 | 0.7% | 270 | 3.2% |
| CIN3121_PRI | 8,083 | 97.0% | 8,045 | 96.5% | 38 | 0.5% | 49 | 0.6% | 206 | 2.4% |
| CIN3121_ALT | 8,069 | 96.8% | 8,027 | 96.3% | 42 | 0.5% | 52 | 0.6% | 217 | 2.6% |
| CIN3122_PRI | 8,093 | 97.0% | 8,058 | 96,6% | 35 | 0.4% | 43 | 0.5% | 202 | 2.5% |
| CIN3122_ALT | 8,019 | 96.2% | 7,970 | 95.6% | 49 | 0.6% | 49 | 0.6% | 270 | 3.2% |
| CRO3131_PRI | 8,080 | 96.9% | 8,044 | 96.5% | 36 | 0.4% | 48 | 0.6% | 210 | 2.5% |
| CRO3131_ALT | 8,079 | 96.9% | 8,040 | 96.4% | 39 | 0.5% | 44 | 0.5% | 215 | 2.6% |
| PIL3664_PRI | 8,089 | 97.0% | 8,049 | 96.5% | 40 | 0.5% | 46 | 0.6% | 203 | 2.4% |
| PIL3664_ALT | 7,974 | 95.6% | 7,938 | 95.2% | 36 | 0.4% | 69 | 0.8% | 295 | 3.6% |

**Table S4. BUSCO scores for the gene annotations.** The BUSCO analysis was performed using the 8,338 single-copy orthologs from the aves_db10 avian database. Details as in Table S3. BUSCO scores for the gene annotations are consistently high and similar among species.

| | C | C% | S | S% | D | D% | F | F% | M | M% |
|---|---|---|---|---|---|---|---|---|---|---|
| RUF3129 | 7,934 | 95.1% | 7,898 | 94.7% | 36 | 0.4% | 74 | 0.9% | 330 | 4.0% |
| IBEB009240 | 7,917 | 94.9% | 7,874 | 94.4% | 43 | 0.5% | 81 | 1.0% | 340 | 4.1% |
| PAL3118 | 7,918 | 94.5% | 7,882 | 94.5% | 36 | 0.4% | 79 | 0.9% | 341 | 4.2% |
| PAL3117 | 7,922 | 95.0% | 7,883 | 94.5% | 39 | 0.5% | 71 | 0.9% | 345 | 4.1% |
| PAL3372 | 7,934 | 95.2% | 7,895 | 94.7% | 39 | 0.5% | 73 | 0.9% | 331 | 3.9% |
| RUF3128 | 7,910 | 94.9% | 7,869 | 94.4% | 41 | 0.5% | 76 | 0.9% | 352 | 4.2% |
| HYPOXB009213 | 7,935 | 95.2% | 7,889 | 94.6% | 46 | 0.6% | 68 | 0.8% | 335 | 4.0% |
| HYPOXB009632 | 7,911 | 94.9% | 7,871 | 94.4% | 40 | 0.5% | 74 | 0.9% | 353 | 4.2% |
| HYPOXB009666 | 7,892 | 94.6% | 7,849 | 94.1% | 43 | 0.5% | 72 | 0.9% | 374 | 4.5% |
| HYPOXB009684 | 7,920 | 95.0% | 7,884 | 94.6% | 26 | 0.4% | 72 | 0.9% | 346 | 4.1% |
| IBEB009696 | 7,925 | 95.0% | 7,883 | 94.5% | 42 | 0.5% | 72 | 0.9% | 341 | 4.1% |
| IBEB009699 | 7,917 | 95.0% | 7,877 | 94.5% | 40 | 0.5% | 69 | 0.8% | 352 | 4.2% |
| CIN3121 | 7,926 | 95.0% | 7,926 | 94.5% | 43 | 0.5% | 70 | 0.8% | 342 | 4.2% |
| CIN3122 | 7,924 | 95.0% | 7,882 | 94.5% | 42 | 0.5% | 70 | 0.8% | 344 | 4.2% |
| CRO3131 | 7,924 | 95.0% | 7,889 | 94.6% | 35 | 0.4% | 67 | 0.8% | 347 | 4.2% |
| PIL3664 | 7,900 | 94.7% | 7,856 | 94.2% | 44 | 0.5% | 71 | 0.9% | 367 | 4.4% |

**Table S5. Summary of genes missing using both annotation and Pangene approaches.** This table shows the percentage of identical positions (BLAST identity) for the 13 genes identified as unique by both the annotation and Pangene approaches in specific species. The genes highlighted in green were detected as present exclusively in the species common to the highlighted columns by both approaches. The BLAST identity values in white represent those same genes in individuals where they were initially detected as missing. These results suggest that the genes identified as putatively unique in some assemblies are actually present in the remaining assemblies, as confirmed by the BLAST high sequence similarity (>95% in many cases). The analysis combines genes unique to individuals within the same species. Species abbreviations: *S. palustris* (PAL), *S. iberaensis* (IBE), *S. ruficollis* (RUF), and *S. hypoxantha* (HYPOX).

| Genes\Individual | HYPOXB009213 | HYPOXB009666 | HYPOXB009684 | RUF3128 | RUF3129 | IBEB009240 | IBEB009632 | IBEB009696 | IBEB009699 | PAL3117 | PAL3118 | PAL3372 |
|---|---|---|---|---|---|---|---|---|---|---|---|---|
| *LOC100224242* | 98.7 | 98.7 | 98.7 | 98.7 | 98.7 | 98.7 | 98.7 | 100 | 97.5 | 100 | 98.7 | 100 |
| *LOC101752242* | 100 | 100 | 100 | 100 | 100 | 100 | 100 | 100 | 100 | 100 | 100 | 100 |
| *LOC107054362* | 97.6 | 92.1 | 92.1 | 96.5 | 94.3 | 96.3 | 93.3 | 89.9 | 96.3 | 94.4 | 100 | 96.4 |
| *NECTIN1* | 100 | 100 | 100 | 100 | 100 | 100 | 100 | 100 | 100 | 100 | 100 | 100 |
| *LOC100224089* | 100 | 100 | 100 | 100 | 100 | 100 | 100 | 100 | 100 | 100 | 100 | 100 |
| *LOC112530980* | 67.6 | 58.0 | 48.5 | 81.4 | 52.8 | 63.2 | 67.2 | 54.6 | 64.9 | 84.4 | 48.3 | 53.4 |
| *ATP5A1Z* | 100 | 100 | 100 | 100 | 100 | 100 | 100 | 100 | 100 | 100 | 100 | 100 |
| *BEST4* | 80.6 | 80.6 | 80.6 | 96.4 | 96.4 | 98.1 | 100 | 100 | 100 | 100 | 100 | 98.2 |
| *CZH18ORF32* | 76.3 | 96.1 | 75.6 | 77.6 | 77.6 | 77.6 | 77.6 | 76.3 | 75.6 | 75.6 | 75.6 | 75.6 |
| *HINTW* | 79.2 | 98.2 | 79.2 | 79.2 | 79.2 | 79.2 | 79.2 | 79.2 | 79.2 | 79.2 | 79.2 | 79.2 |
| *MAPK4* | 89.7 | 100 | 74.2 | 95.2 | 95.2 | 95.2 | 92.9 | 95.2 | 95.2 | 90 | 95.2 | 95.2 |
| *SV2A* | 100 | 100 | 100 | 100 | 100 | 100 | 100 | 100 | 100 | 100 | 100 | 100 |
| *TREM-B2* | 87.9 | 88.6 | 90.9 | 90.1 | 95.8 | 89.4 | 95.8 | 93.3 | 90.2 | 89.4 | 88.7 | 96.3 |

**Table S6. GWAS and $F_{ST}$ outlier peaks.** For the 10 peaks that we detected, this table includes the scaffold number, start, end, and length of the peaks, their identification as GWAS and/or $F_{ST}$ outliers, genes within the peaks (or closest genes, highlighted in yellow with the distance to the peak specified in brackets), associated patches and pigments (eumelanin and pheomelanin), $F_{ST}$ comparisons with significant outliers (including the number of SNPs and SVs with $F_{ST}$ > 0.75 in brackets). Peaks, patches/pigments and comparisons detected with both SNPs and SVs are highlighted in purple, the ones detected only with SNPs and SVs are in green and blue, respectively. The orange peak represents one detected with both SNPs and SVs but classified as an $F_{ST}$ outlier in only one window (not considered a peak in our analysis) when using SVs. Bolded columns indicate peaks identified as outliers by both GWAS and $F_{ST}$ analyses. Asterisks (*) mark peaks also detected through variants within TEs. To determine the start and end of peaks detected in both the SNP and SV datasets, we used the lowest and highest coordinates identified from either marker type. We indicate the marker type that defined the boundary of the peak in green if they originate from the SNP dataset, and purple if they are identical in both.

| **Peak ID** | **Peak 1** | **Peak 2** | **Peak 3** | **Peak 4** | **Peak 5** | Peak 6 | Peak 7 | Peak 8 | Peak 9 | Peak 10 |
|---|---|---|---|---|---|---|---|---|---|---|
| **Scaffold** | **10*** | **14** | **21*** | **7*** | **9*** | 41 | 7 | 3* | 7 | 7 |
| **Start_peak** | **6,190,001** | **13,950,001** | **13,545,422** | **25,260,001** | **15,010,001** | 108,180 | 5,260,234 | 9,950,001 | 5,560,001 | 5,730,001 |
| **End_Peak** | **6,250,000** | **13,990,000** | **13,700,000** | **25,320,000** | **15,030,000** | 112,141 | 5,272,125 | 9,990,000 | 5,580,000 | 5,750,000 |
| **GWAS/$F_{ST}$** | **GWAS/ $F_{ST}$** | **GWAS/ $F_{ST}$** | **GWAS/ $F_{ST}$** | **GWAS/ $F_{ST}$** | **GWAS/ $F_{ST}$** | GWAS | GWAS | $F_{ST}$ | $F_{ST}$ | $F_{ST}$ |
| **Gene1** | ***HERC2*** | ***GPT2*** | ***ASIP*** | ***TYRP1*** | ***ZNF503* (20,255bp)** | *ELAC* (7,658bp) | *SLC45A2* | *ALB* | *PRLR* (21,985bp) | *SPEF2* |
| **Gene2** | | **CDCA9** | **AHCY** | | | | | | | *LOC112530520* |
| **GWAS 1** | **rump_eu** | **head_phe** | **belly_eu** | **rump_eu** | **rump_eu** | rump_eu | throat_eu | | | |
| **GWAS 2** | **belly_phe** | | **rump_eu** | **rump_phe** | **belly_eu** | head_eu | back_phe | | | |
| **GWAS 3** | **rump_phe** | | **back_eu** | **throat_eu** | **rump_phe** | head_phe | nape_phe | | | |
| **GWAS 4** | | | **nape_phe** | **belly_phe** | **throat_eu** | | rump_phe | | | |
| **GWAS 5** | | | **throat_eu** | **head_phe** | **nape_phe** | | rump_eu | | | |
| **GWAS 6** | | | **rump_phe** | **belly_eu** | | | | | | |
| **GWAS 7** | | | **head_eu** | **nape_phe** | | | | | | |
| **GWAS 8** | | | **head_phe** | | | | | | | |
| **GWAS 9** | | | **belly_phe** | | | | | | | |
| **GWAS 10** | | | **nape_eu** | | | | | | | |
| **GWAS 11** | | | **belly_eu** | | | | | | | |
| **GWAS 12** | | | **back_phe** | | | | | | | |
| **$F_{ST}$ comp1** | **HYPOX_RUF (61, 10)** | **HYPOX_IBE (10, 1)** | **HYPOX_CIN (23, 4)** | **IBE_CIN (13, 4)** | **RUF_MEL (19, 3)** | | | HYPOX_MEL (47, 3) | MEL_IBE (2, 0) | MEL_IBE (6, 0) |

| | | | | | | |
|---|---|---|---|---|---|---|
| $F_{ST}$ comp2 | RUF_CIN (17, 1) | IBE_CIN (9, 2) | MEL_CIN (92, 10) | RUF_CIN (35, 5) | HYPOX_MEL (11, 0) | MEL_IBE (52, 4) |
| $F_{ST}$ comp3 | HYPOX_IBE (84, 12) | RUF_IBE (11, 2) | MEL_IBE (57, 6) | RUF_HYPOCH (30, 3) | MEL_CIN (5, 0) | RUF_MEL (57, 3) |
| $F_{ST}$ comp4 | HYPOX_MEL (47, 8) | IBE_HYPOCH (3, 1) | HYPOX_MEL (46, 6) | HYPOX_MEL (13, 0) | MEL_HYPOCH (4, 0) | RUF_HYPOCH (0, 1) |
| $F_{ST}$ comp5 | HYPOX_CIN (25, 3) | MEL_IBE (4, 1) | RUF_MEL (45, 5) | HYPOX_RUF (1, 0) | RUF_CIN (1, 1) | RUF_IBE (0, 1) |
| $F_{ST}$ comp6 | IBE_HYPOCH (7, 1) | | MEL_HYPOCH (41, 6) | IBE_HYPOCH (18, 2) | | |
| $F_{ST}$ comp7 | RUF_IBE (34, 2) | | RUF_CIN (9, 0) | MEL_CIN (30, 3) | | |
| $F_{ST}$ comp8 | RUF_MEL (30, 2) | | | MEL_HYPOCH (24, 0) | | |
| $F_{ST}$ comp9 | | | | RUF_MEL (30, 3) | | |
| $F_{ST}$ comp10 | | | | HYPOX_IBE (4, 0) | | |
| $F_{ST}$ comp11 | | | | MEL_IBE (24, 2) | | |

**Table S7. GWAS singleton outliers outside peaks.** This table includes the scaffold number, position, GWAS trait with significant association (patches 1, 2, and 3 when the same variant is an outlier in different analyses), variant type, whether the variant overlaps TEs or coding regions (CDS), the overlapping genes, and SV average lengths and types. The presence of multiple SV types for certain variants is due to different alternative alleles corresponding to different SV categories, with the SV length representing the average among all alternative alleles. The variant types include: single nucleotide polymorphism (snp), multiple-nucleotide polymorphism (mnp), insertions (ins), deletions (del) and complex variants (complex). The GWAS identified 217 singletons outside the main peaks discussed in the manuscript, of which 90% were SNPs and 10% were SVs, overlapping with 58 and 7 genes, respectively. Among the outlier SNPs, 7.6% overlapped with TEs, 9.2% with coding sequences (CDS), and three SNPs overlapped both TEs and CDS, in the genes *FRMD5*, *MN1*, and *DIABLO*. For the SVs outside the peaks, 22% overlapped TEs, but none overlapped CDS. Nearly half of the SVs (~45.5%) were classified as complex, 41% as deletions, and the remaining as insertions.

| Scaffold# | Position (bp) | Patch 1 | Patch 2 | Patch 3 | Variant | Within TE | Within CDS | Gene | Length | SV type |
|---|---|---|---|---|---|---|---|---|---|---|
| 1 | 7,578,375 | head_eu | | | SNP | Yes | No | | 1 | - |
| 1 | 37,640,840 | head_eu | head_phe | | SNP | No | No | | 1 | - |
| 1 | 57,439,079 | head_phe | | | SNP | No | No | *MYB* | 1 | - |
| 1 | 59,303,255 | head_phe | | | SNP | No | No | *EPB41L2* | 1 | - |
| 1 | 59,843,989 | head_eu | | | SNP | No | No | | 1 | - |
| 1 | 66,226,001 | belly_eu | | | SNP | No | No | | 1 | - |
| 1 | 69,255,668 | rump_phe | throat_eu | | SNP | No | Yes | *OSTM1* | 1 | - |
| 1 | 70,979,897 | rump_phe | | | SNP | No | No | | 1 | - |
| 1 | 89,677,114 | head_phe | | | SNP | No | No | | 1 | - |
| 1 | 99,171,563 | head_phe | | | SNP | No | No | | 1 | - |
| 1 | 107,186,658 | belly_eu | nape_phe | | SNP | No | No | | 1 | - |
| 1 | 107,186,884 | belly_eu | | | SNP | No | No | | 1 | - |
| 1 | 118,480,298 | head_eu | | | SNP | No | No | | 1 | - |
| 1 | 121,098,490 | head_eu | | | SNP | No | No | | 1 | - |
| 10 | 4,430,034 | head_eu | | | SNP | No | No | *DHRSX* | 1 | - |
| 10 | 11,588,001 | head_eu | | | SNP | No | No | | 1 | - |
| 10 | 14,986,941 | rump_eu | | | SNP | No | No | | 1 | - |
| 10 | 17,414,771 | head_phe | | | SNP | No | No | | 1 | - |
| 10 | 17,414,776 | head_phe | | | SNP | No | No | | 1 | - |
| 10 | 20,805,932 | head_phe | | | SNP | No | No | | 1 | - |
| 10 | 22,079,259 | head_phe | | | SNP | No | No | *ZRSR2* | 1 | - |
| 10 | 27,527,039 | belly_eu | | | SNP | No | No | | 1 | - |
| 11 | 9,102,671 | throat_eu | | | SNP | No | No | *UQCRH* | 1 | - |
| 11 | 15,946,202 | throat_eu | | | SNP | Yes | No | | 1 | - |
| 11 | 18,648,931 | rump_phe | | | SNP | NO | No | *SNX7* | 1 | - |
| 11 | 18,944,959 | belly_phe | rump_phe | | SNP | NO | No | | 1 | - |

| | | | | | | | | | | |
|---|---|---|---|---|---|---|---|---|---|---|
| 11 | 21,207,822 | throat_eu | | | SNP | No | No | | 1 | - |
| 11 | 29,419,248 | rump_eu | | | SNP | No | No | | 1 | - |
| 11 | 29,999,750 | rump_eu | throat_eu | | SNP | No | No | | 1 | - |
| 12 | 2,562,099 | head_eu | | | SNP | No | No | | 1 | - |
| 12 | 7,822,348 | head_eu | | | SNP | No | No | | 1 | - |
| 12 | 7,884,106 | head_phe | | | SNP | No | No | | 1 | - |
| 12 | 10,113,698 | rump_eu | rump_phe | throat_eu | SNP | No | No | *LSM3* | 1 | - |
| 12 | 15,221,168 | rump_eu | | | SNP | No | No | | 1 | - |
| 13 | 537,329 | rump_phe | | | SNP | Yes | Yes | *FRMD5* | 1 | - |
| 13 | 4,170,303 | belly_eu | | | SNP | No | No | | 1 | - |
| 13 | 12,403,765 | rump_phe | | | SNP | No | No | | 1 | - |
| 13 | 20,011,977 | throat_phe | | | SNP | No | No | | 1 | - |
| 14 | 4,432,552 | head_phe | | | SNP | No | No | | 1 | - |
| 14 | 4,878,014 | belly_phe | | | SNP | Yes | No | | 1 | - |
| 14 | 5,044,156 | throat_eu | rump_phe | | SNP | No | No | *ATMIN* | 1 | - |
| 14 | 7,193,549 | rump_eu | | | SNP | No | Yes | *E2F4* | 1 | - |
| 14 | 11,314,890 | belly_eu | | | SNP | No | No | | 1 | - |
| 15 | 114,607 | throat_eu | rump_eu | | SNP | No | Yes | *EVA1C* | 1 | - |
| 15 | 7,252,899 | head_phe | | | SNP | No | No | | 1 | - |
| 15 | 7,252,993 | head_eu | | | SNP | No | No | | 1 | - |
| 15 | 13,047,150 | head_phe | | | SNP | Yes | No | | 1 | - |
| 16 | 291,874 | rump_eu | throat_eu | | SNP | No | No | | 1 | - |
| 16 | 2,121,219 | throat_phe | | | SNP | No | No | *STK10* | 1 | - |
| 16 | 2,472,953 | head_phe | | | SNP | No | No | | 1 | - |
| 16 | 5,763,748 | rump_eu | throat_eu | | SNP | No | No | | 1 | - |
| 16 | 9,293,614 | rump_eu | throat_eu | | SNP | No | No | | 1 | - |
| 16 | 11,387,256 | head_eu | | | SNP | No | No | | 1 | - |
| 16 | 12,017,853 | head_eu | head_phe | | SNP | No | No | *GABRG2* | 1 | - |
| 16 | 16,022,267 | head_phe | | | SNP | No | No | *ANKHD1* | 1 | - |
| 17 | 7,529,826 | head_eu | | | SNP | No | No | | 1 | - |
| 17 | 9,908,833 | rump_phe | | | SNP | No | No | | 1 | - |
| 17 | 17,375,275 | back_phe | | | SNP | No | No | | 1 | - |
| 19 | 2,440,755 | head_phe | | | SNP | No | No | | 1 | - |
| 19 | 10,484,089 | belly_phe | rump_phe | throat_eu | SNP | No | No | | 1 | - |
| 2 | 4,179,546 | head_eu | | | SNP | No | No | | 1 | - |
| 2 | 17,848,763 | head_eu | | | SNP | Yes | No | | 1 | - |
| 2 | 24,715,239 | head_eu | head_phe | | SNP | No | No | | 1 | - |
| 2 | 36,308,182 | head_phe | | | SNP | No | No | | 1 | - |
| 2 | 40,049,832 | head_eu | | | SNP | No | No | | 1 | - |
| 2 | 40,763,383 | head_eu | | | SNP | No | No | | 1 | - |

| | | | | | | | | | | |
|---|---|---|---|---|---|---|---|---|---|---|
| 2 | 62,055,562 | head_eu | | | SNP | No | No | *EYA1* | 1 | - |
| 201 | 14,063 | rump_phe | | | SNP | No | No | *FARSA* | 1 | - |
| 21 | 652,076 | head_phe | | | SNP | No | No | | 1 | - |
| 21 | 14,551,660 | head_eu | | | SNP | No | No | | 1 | - |
| 21 | 14,562,517 | rump_eu | | | SNP | No | No | | 1 | - |
| 22 | 1,995,702 | rump_eu | throat_eu | | SNP | No | Yes | *TCN2* | 1 | - |
| 22 | 3,901,398 | head_phe | | | SNP | No | Yes | *DEPDC5* | 1 | - |
| 22 | 4,244,650 | rump_phe | throat_eu | | SNP | No | No | | 1 | - |
| 22 | 6,961,401 | rump_phe | | | SNP | No | No | | 1 | - |
| 22 | 9,763,957 | throat_eu | | | SNP | Yes | Yes | *MN1* | 1 | - |
| 22 | 11,834,541 | rump_eu | | | SNP | Yes | Yes | *DIABLO* | 1 | - |
| 23 | 3,262,000 | head_eu | head_phe | | SNP | No | No | *KIAA0556* | 1 | - |
| 23 | 7,900,486 | head_phe | | | SNP | No | No | | 1 | - |
| 24 | 5,578,393 | rump_eu | rump_phe | throat_eu | SNP | No | No | | 1 | - |
| 24 | 5,578,498 | rump_eu | | | SNP | No | No | | 1 | - |
| 24 | 6,076,542 | head_eu | | | SNP | No | No | | 1 | - |
| 24 | 6,995,672 | head_eu | | | SNP | No | No | | 1 | - |
| 24 | 9,061,097 | throat_eu | | | SNP | No | Yes | *DFNB31* | 1 | - |
| 25 | 9,229,072 | rump_phe | | | SNP | No | No | | 1 | - |
| 26 | 4,903,331 | head_phe | | | SNP | No | No | | 1 | - |
| 26 | 9,215,214 | rump_phe | | | SNP | No | No | | 1 | - |
| 27 | 8,010,876 | head_phe | | | SNP | No | No | | 1 | - |
| 27 | 9,442,630 | rump_phe | | | SNP | No | No | | 1 | - |
| 28 | 516,644 | head_eu | head_phe | | SNP | No | No | *MCC* | 1 | - |
| 29 | 2,305,057 | rump_eu | throat_eu | | SNP | No | No | | 1 | - |
| 29 | 6,186,776 | rump_eu | rump_phe | | SNP | No | No | *SNTG2* | 1 | - |
| 3 | 314,107 | head_phe | | | SNP | No | No | *ELF2* | 1 | - |
| 3 | 1,291,975 | head_phe | | | SNP | No | No | | 1 | - |
| 3 | 2,132,212 | head_eu | | | SNP | No | No | *SLC7A11* | 1 | - |
| 3 | 3,255,463 | head_eu | head_phe | | SNP | No | No | | 1 | - |
| 3 | 7,540,921 | rump_eu | | | SNP | No | No | *SPATA5* | 1 | - |
| 3 | 14,324,086 | head_phe | | | SNP | No | No | | 1 | - |
| 3 | 24,060,252 | rump_phe | | | SNP | Yes | No | | 1 | - |
| 3 | 24,735,176 | rump_eu | | | SNP | No | No | *PITX2* | 1 | - |
| 3 | 25,285,978 | rump_phe | throat_eu | | SNP | No | No | | 1 | - |
| 3 | 31,269,872 | head_eu | head_phe | | SNP | No | No | | 1 | - |
| 3 | 51,178,437 | head_phe | | | SNP | No | No | *ARAP2* | 1 | - |
| 3 | 59,986,859 | head_phe | | | SNP | Yes | No | *KDM3A* | 1 | - |
| 30 | 4,053,075 | head_phe | | | SNP | No | No | | 1 | - |
| 30 | 4,461,834 | head_eu | head_phe | | SNP | No | No | *GRAMD1B* | 1 | - |

| | | | | | | | | | |
|---|---|---|---|---|---|---|---|---|---|
| 30 | 7,672,830 | rump_phe | | SNP | No | No | *BUD13* | 1 | - |
| 30 | 8,247,489 | throat_eu | rump_eu | SNP | No | No | | 1 | - |
| 31 | 1,157,356 | throat_eu | | SNP | No | No | | 1 | - |
| 31 | 4,072,881 | rump_eu | | SNP | No | No | | 1 | - |
| 31 | 4,853,417 | rump_eu | | SNP | No | Yes | *SMPDL3B* | 1 | - |
| 33 | 1,301,983 | rump_phe | | SNP | No | No | | 1 | - |
| 34 | 413,039 | rump_eu | | SNP | No | No | | 1 | - |
| 34 | 1,654,438 | rump_phe | | SNP | No | No | | 1 | - |
| 35 | 5,903,729 | head_eu | | SNP | No | No | *CKMT2* | 1 | - |
| 37 | 606,455 | rump_eu | | SNP | No | No | *TIMP2* | 1 | - |
| 37 | 897,886 | rump_phe | | SNP | No | No | | 1 | - |
| 37 | 897,910 | rump_eu | rump_phe | SNP | No | No | | 1 | - |
| 37 | 4,084,060 | rump_eu | throat_eu | SNP | No | No | | 1 | - |
| 38 | 1,095,246 | rump_eu | throat_eu | SNP | No | No | | 1 | - |
| 38 | 3,685,489 | rump_eu | | SNP | No | No | | 1 | - |
| 38 | 3,686,499 | rump_eu | | SNP | No | No | *HCN2* | 1 | - |
| 38 | 5,293,207 | throat_eu | | SNP | No | No | | 1 | - |
| 38 | 5,865,087 | throat_eu | rump_eu | SNP | No | No | *PDE4C* | 1 | - |
| 38 | 6,016,243 | head_eu | head_phe | SNP | No | No | | 1 | - |
| 38 | 6,036,064 | rump_eu | | SNP | No | No | | 1 | - |
| 38 | 6,340,581 | rump_eu | | SNP | No | No | | 1 | - |
| 39 | 851,907 | throat_eu | rump_eu | SNP | No | No | *PLEKHA6* | 1 | - |
| 4 | 2,182,679 | rump_eu | throat_eu | SNP | No | No | | 1 | - |
| 4 | 8,962,803 | rump_phe | throat_eu | SNP | No | No | | 1 | - |
| 4 | 11,871,860 | head_phe | | SNP | No | No | | 1 | - |
| 4 | 14,727,864 | throat_phe | | SNP | No | No | *PTPN5* | 1 | - |
| 4 | 19,663,082 | rump_phe | | SNP | No | No | *COMMD9* | 1 | - |
| 4 | 39,661,380 | rump_eu | throat_eu | SNP | No | No | | 1 | - |
| 4 | 40,155,146 | throat_eu | | SNP | No | No | | 1 | - |
| 4 | 45,096,065 | head_eu | | SNP | No | No | | 1 | - |
| 4 | 47,816,905 | rump_eu | throat_eu | SNP | No | No | | 1 | - |
| 4 | 49,907,794 | belly_eu | | SNP | No | No | | 1 | - |
| 4 | 53,482,965 | belly_phe | rump_phe | SNP | No | No | | 1 | - |
| 40 | 1,027,017 | head_phe | | SNP | No | No | | 1 | - |
| 40 | 2,262,636 | rump_phe | | SNP | No | No | *CNTNAP1* | 1 | - |
| 42 | 2,602,528 | head_eu | | SNP | Yes | No | | 1 | - |
| 44 | 3,056,988 | rump_phe | throat_eu | SNP | No | No | | 1 | - |
| 44 | 3,057,026 | throat_eu | rump_phe | SNP | No | No | | 1 | - |
| 47 | 399,559 | head_eu | head_phe | SNP | No | No | | 1 | - |
| 48 | 648,895 | rump_phe | | SNP | No | No | | 1 | - |

| | | | | | | | | | | |
|---|---|---|---|---|---|---|---|---|---|---|
| 5 | 2,463,675 | head_phe | | SNP | No | No | | 1 | | - |
| 5 | 3,593,173 | head_eu | | SNP | No | No | *SEMA3D* | 1 | | - |
| 5 | 3,976,790 | head_eu | | SNP | No | No | | 1 | | - |
| 5 | 3,981,472 | head_eu | | SNP | Yes | No | | 1 | | - |
| 5 | 4,957,179 | head_eu | | SNP | No | No | | 1 | | - |
| 5 | 10,880,876 | head_phe | | SNP | No | No | | 1 | | - |
| 5 | 20,568,670 | throat_eu | | SNP | No | No | | 1 | | - |
| 5 | 35,314,628 | head_phe | | SNP | No | No | | 1 | | - |
| 5 | 44,614,599 | head_eu | | SNP | No | No | | 1 | | - |
| 5 | 45,540,030 | rump_eu | | SNP | No | Yes | *PDXP* | 1 | | - |
| 5 | 47,655,611 | throat_eu | | SNP | No | No | *FBXO7* | 1 | | - |
| 5 | 57,627,929 | head_phe | | SNP | No | No | *GNPTAB* | 1 | | - |
| 5 | 58,087,466 | head_eu | | SNP | No | No | | 1 | | - |
| 55 | 1,460,315 | head_phe | | SNP | No | No | *CYP26B1* | 1 | | - |
| 56 | 1,204,891 | rump_eu | | SNP | No | Yes | *LOC107055878* | 1 | | - |
| 56 | 1,436,112 | rump_phe | throat_eu | SNP | No | Yes | *CHPF2* | 1 | | - |
| 58 | 265,479 | rump_eu | | SNP | No | No | | 1 | | - |
| 59 | 892,146 | rump_eu | | SNP | No | No | *DNMT3A* | 1 | | - |
| 6 | 3,729,745 | head_phe | | SNP | Yes | No | | 1 | | - |
| 6 | 3,730,550 | head_eu | | SNP | No | No | | 1 | | - |
| 6 | 3,731,705 | head_eu | head_phe | SNP | No | No | | 1 | | - |
| 6 | 3,735,447 | head_phe | | SNP | No | No | | 1 | | - |
| 6 | 3,735,702 | head_eu | head_phe | SNP | No | No | | 1 | | - |
| 6 | 4,666,353 | head_phe | | SNP | No | No | | 1 | | - |
| 6 | 20,448,384 | head_phe | | SNP | No | No | | 1 | | - |
| 6 | 21,250,051 | rump_eu | throat_eu | SNP | No | No | | 1 | | - |
| 6 | 35,210,332 | head_eu | | SNP | No | No | | 1 | | - |
| 6 | 48,719,134 | rump_phe | throat_eu | SNP | Yes | No | | 1 | | - |
| 6 | 49,903,583 | head_phe | | SNP | No | No | | 1 | | - |
| 6 | 55,920,639 | head_eu | | SNP | Yes | No | | 1 | | - |
| 60 | 903,836 | rump_eu | | SNP | No | No | | 1 | | - |
| 63 | 534,660 | head_phe | | SNP | No | No | *SKAP1* | 1 | | - |
| 7 | 5,257,286 | nape_eu | | SNP | No | No | *SLC45A2* | 1 | | - |
| 7 | 22,496,681 | head_phe | | SNP | No | No | | 1 | | - |
| 7 | 52,217,248 | head_phe | | SNP | No | No | | 1 | | - |
| 7 | 25,717,703 | head_phe | | SNP | No | No | | 1 | | - |
| 7 | 25,720,544 | head_phe | | SNP | No | No | | 1 | | - |
| 8 | 683,217 | rump_eu | | SNP | No | No | *GALNT5* | 1 | | - |
| 8 | 7,828,590 | rump_phe | throat_eu | SNP | No | No | *ACTR3* | 1 | | - |
| 8 | 9,494,004 | rump_eu | throat_eu | SNP | No | Yes | *ESYT3* | 1 | | - |

| | | | | | | | | | |
|---|---|---|---|---|---|---|---|---|---|
| 8 | 14,962,840 | rump_phe | | SNP | Yes | No | *IGFBP2* | 1 | - |
| 8 | 17,487,835 | throat_eu | | SNP | No | No | *IFIH1* | 1 | - |
| 8 | 20,406,173 | rump_phe | | SNP | No | No | | 1 | - |
| 8 | 20,584,199 | rump_phe | | SNP | No | No | | 1 | - |
| 8 | 23,379,412 | head_phe | | SNP | No | No | | 1 | - |
| 8 | 28,281,792 | rump_eu | throat_eu | SNP | No | No | | 1 | - |
| 8 | 35,747,120 | head_phe | | SNP | Yes | No | | 1 | - |
| 87 | 440,167 | throat_eu | | SNP | No | No | | 1 | - |
| 9 | 22,427,681 | head_phe | | SNP | No | No | | 1 | - |
| 9 | 23,666,504 | head_phe | | SNP | Yes | No | | 1 | - |
| 9 | 36,454,825 | throat_eu | | SNP | No | Yes | *DPYSL4* | 1 | - |
| 106 | 159,086 | rump_eu | | SV | No | No | | 25.80 | del,complex |
| 24 | 1,673,081 | rump_phe | | SV | No | No | | 1 | del |
| 20 | 3,589,633 | rump_eu | | SV | No | No | | 24.75 | del,snp,mnp |
| 24 | 6,064,183 | rump_eu | | SV | No | No | *SH2D3C* | 1 | del |
| 22 | 6,286,259 | rump_phe | | SV | No | No | *LOC101234199* | 1 | del |
| 25 | 7,141,747 | rump_eu | | SV | No | No | *SEBOX* | 1 | del |
| 2 | 16,900,423 | head_eu | | SV | No | No | | 12.81 | ins |
| 14 | 19,123,028 | rump_eu | | SV | No | No | | 11.00 | ins |
| 1 | 66,905,159 | rump_eu | | SV | Yes | No | | 9.74 | snp,del,complex,mnp,ins |
| 2 | 69,354,581 | rump_eu | | SV | Yes | No | | 1 | del |
| 2 | 80,341,265 | rump_eu | | SV | No | No | | 82.87 | ins |
| 14 | 19,729,351 | head_eu | | SV | No | No | *LOC100228259* | 25.00 | complex |
| 23 | 6,399,290 | throat_eu | | SV | No | No | | 1 | del |
| 27 | 366,063 | rump_eu | | SV | Yes | No | | 6.50 | complex,ins,snp,del |
| 3 | 29,915,880 | throat_eu | | SV | No | No | *DCHS2* | 7.50 | del,snp |
| 4 | 38,634,089 | rump_eu | | SV | Yes | No | | 16.36 | complex,mnp,del,snp |
| 40 | 1,216,889 | rump_eu | | SV | No | No | *PPP1R1B* | 10.27 | complex,snp,mnp,del |
| 41 | 112,001 | head_phe | | SV | Yes | No | | 1 | del |
| 45 | 158,883 | rump_eu | | SV | Yes | No | | 314.38 | ins,complex |
| 50 | 1,529,481 | rump_eu | | SV | No | No | *MEX3A* | 2.71 | snp,complex,mnp,ins |
| 55 | 925,590 | throat_phe | | SV | No | No | | 5.00 | ins,complex |
| 6 | 40,762,710 | rump_eu | | SV | No | No | | 6.67 | ins,del |

**Table S8. $F_{ST}$ outliers for each dataset.** The table includes the number of SNPs, SVs (grouped as < or > 50 bp), and SNPs/SVs within and outside repetitive elements and TEs for each $F_{ST}$ comparison. Details include the number of genome-wide variants with $F_{ST} > 0.75$, number of outlier windows, peaks, and windows outside of peaks. Most of the variants with $F_{ST} > 0.75$ were found in the SNP dataset, and the majority of the outlier windows and peaks were also recovered by the SNP dataset, followed by the dataset of variants outside repetitive elements and TEs. The dataset with the fewest outliers was the long SVs. *In the *S. cinnamomea* vs *S. hypochroma* comparison, there are no variants with $F_{ST} > 0.75$. However, there are variants with $F_{ST} > 0.5$ that can be used to distinguish the species, specifically: 191 for SNPs, 28 for SVs < 50bp, 0 for SVs > 50bp, 24 for variants within TEs, and 195 for variants outside TEs. Besides the 55 bp deletion, an additional 8 SVs > 50 bp are found with $F_{ST} > 0.75$; however, they are outside the peaks. Some of these variants appear in more than one comparison: one in five comparisons, and another in two comparisons. For details, see Table S10.

| | SNPs & MNPs with minor allele count > 4 | SVs < 50bp | SVs > 50bp | SNPs & SVs within TEs | SNPs & SVs outside TEs | Comparison |
|---|---|---|---|---|---|---|
| #variants | 27,875,982 | 3,434,358 | 96,682 | 4,388,258 | 27,018,764 | All |
| # variants $F_{ST}$>0.75 | 0* | 0* | 0* | 0* | 0* | cin_hypoch |
| # outlier windows | 0 | 0 | 0 | 0 | 0 | cin_hypoch |
| # peaks | 0 | 0 | 0 | 0 | 0 | cin_hypoch |
| # out windows outside the peaks | 0 | 0 | 0 | 0 | 0 | cin_hypoch |
| # variants $F_{ST}$ >0.75 | 70 | 12 | 1 | 21 | 62 | hypox_cin |
| # outlier windows | 9 | 3 | 1 | 9 | 6 | hypox_cin |
| # peaks | 2 | 2 | 0 | 1 | 2 | hypox_cin |
| # out windows outside the peaks | 3 | 0 | 1 | 7 | 0 | hypox_cin |
| # variants $F_{ST}$ >0.75 | 10 | 3 | 2 | 6 | 9 | hypox_hypoch |
| # outlier windows | 0 | 1 | 1 | 4 | 0 | hypox_hypoch |
| # peaks | 0 | 0 | 0 | 0 | 0 | hypox_hypoch |
| # out windows outside the peaks | 0 | 1 | 1 | 4 | 0 | hypox_hypoch |
| # variants $F_{ST}$ >0.75 | 98 | 13 | 1 | 5 | 107 | hypox_ibe |
| # outlier windows | 8 | 4 | 1 | 3 | 8 | hypox_ibe |
| # peaks | 3 | 2 | 1 | 1 | 3 | hypox_ibe |
| # out windows outside the peaks | 0 | 0 | 0 | 0 | 0 | hypox_ibe |
| # variants $F_{ST}$ >0.75 | 234 | 24 | 0 | 31 | 227 | hypox_mel |
| # outlier windows | 16 | 7 | 0 | 4 | 18 | hypox_mel |
| # peaks | 5 | 3 | 0 | 2 | 5 | hypox_mel |
| # out windows outside the peaks | 1 | 0 | 0 | 0 | 1 | hypox_mel |
| # variants $F_{ST}$ >0.75 | 154 | 19 | 0 | 21 | 152 | hypox_ruf |
| # outlier windows | 6 | 4 | 0 | 5 | 7 | hypox_ruf |
| # peaks | 2 | 1 | 0 | 1 | 2 | hypox_ruf |
| # out windows outside the peaks | 0 | 0 | 0 | 3 | 1 | hypox_ruf |
| # variants $F_{ST}$ >0.75 | 51 | 23 | 2 | 39 | 37 | ibe_cin |
| # outlier windows | 9 | 3 | 1 | 8 | 6 | ibe_cin |

| | | | | | | |
|---|---|---|---|---|---|---|
| # peaks | 2 | 2 | 0 | 1 | 2 | ibe_cin |
| # out windows outside the peaks | 3 | 0 | 1 | 7 | 0 | ibe_cin |
| # variants $F_{ST}$ >0.75 | 42 | 14 | 0 | 16 | 40 | ibe_hypoch |
| # outlier windows | 7 | 4 | 0 | 7 | 5 | ibe_hypoch |
| # peaks | 3 | 3 | 0 | 1 | 3 | ibe_hypoch |
| # out windows outside the peaks | 2 | 1 | 0 | 5 | 0 | ibe_hypoch |
| # variants $F_{ST}$ >0.75 | 1,640 | 210 | 4 | 372 | 1,482 | mel_cin |
| # outlier windows | 17 | 19 | 2 | 26 | 17 | mel_cin |
| # peaks | 3 | 2 | 0 | 1 | 3 | mel_cin |
| # out windows outside the peaks | 3 | 13 | 2 | 22 | 6 | mel_cin |
| # variants $F_{ST}$ >0.75 | 1,183 | 132 | 2 | 226 | 1,091 | mel_hypoch |
| # outlier windows | 8 | 9 | 1 | 19 | 10 | mel_hypoch |
| # peaks | 3 | 1 | 0 | 1 | 3 | mel_hypoch |
| # out windows outside the peaks | 1 | 7 | 1 | 18 | 2 | mel_hypoch |
| # variants $F_{ST}$ >0.75 | 566 | 59 | 0 | 80 | 545 | mel_ibe |
| # outlier windows | 21 | 9 | 0 | 17 | 23 | mel_ibe |
| # peaks | 5 | 4 | 0 | 3 | 5 | mel_ibe |
| # out windows outside the peaks | 3 | 0 | 0 | 9 | 4 | mel_ibe |
| # variants $F_{ST}$ >0.75 | 1,923 | 260 | 1 | 361 | 1,823 | ruf_cin |
| # outlier windows | 18 | 25 | 1 | 32 | 16 | ruf_cin |
| # peaks | 4 | 3 | 0 | 0 | 4 | ruf_cin |
| # out windows outside the peaks | 9 | 20 | 1 | 32 | 7 | ruf_cin |
| # variants $F_{ST}$ >0.75 | 1,298 | 152 | 0 | 234 | 1,216 | ruf_hypoch |
| # outlier windows | 19 | 12 | 0 | 20 | 16 | ruf_hypoch |
| # peaks | 1 | 2 | 0 | 1 | 1 | ruf_hypoch |
| # out windows outside the peaks | 15 | 10 | 0 | 18 | 12 | ruf_hypoch |
| # variants $F_{ST}$ >0.75 | 874 | 122 | 1 | 133 | 864 | ruf_ibe |
| # outlier windows | 13 | 18 | 1 | 22 | 14 | ruf_ibe |
| # peaks | 2 | 2 | 0 | 1 | 2 | ruf_ibe |
| # out windows outside the peaks | 9 | 15 | 1 | 21 | 10 | ruf_ibe |
| # variants $F_{ST}$ >0.75 | 217 | 21 | 0 | 39 | 199 | ruf_mel |
| # outlier windows | 20 | 9 | 0 | 13 | 19 | ruf_mel |
| # peaks | 5 | 5 | 0 | 5 | 5 | ruf_mel |
| # out windows outside the peaks | 3 | 0 | 0 | 10 | 3 | ruf_mel |

**Table S9. $F_{ST}$ outlier windows outside peaks.** The table provides details on window coordinates, comparisons with significant $F_{ST}$, variant types and lengths, overlap with genes and TEs, the total number of SNPs and SVs, as well as the number and length within TEs, highly differentiated variants ($F_{ST} > 0.75$), including SNPs within coding regions, SVs within and outside coding regions, and SVs > 50 bp overlapping coding regions. Windows detected with both SNPs and SVs are highlighted in purple, the windows and comparisons detected only with SNPs and SVs are in green and blue, respectively. The proportion of SVs in outlier windows was higher, accounting for 61% of the cases, than that of SNPs which made up only 39% of such windows. One window was identified as an outlier for both SNPs and SVs, but in different comparisons. Among the outlier windows detected with SVs, 32 (61%) overlapped with genes and 23 (70%) of the SNP outlier windows overlapped with genes. The single window detected by both datasets but in different comparisons also overlapped with a gene. These results show that while the main regions of differentiation are concentrated in peaks, we are still able to detect smaller regions (in this case 10 kb windows) which also show differences across Capuchinos.

See Excel file.

**Table S10. Structural variants longer than 50 bp with $F_{ST}$ > 0.75 outside the peaks.** The table includes the scaffold, position, population comparisons with their corresponding $F_{ST}$ values, overlapping genes (bold) or the closest gene with the distance in brackets, SV length (the mean length among all alleles), and SV type. The presence of multiple SV types for certain variants is due to different alternative alleles corresponding to different SV categories, with the SV length representing the average among all alternative alleles. The variant types include: insertions (ins), deletions (del) and complex variants (complex). The mean SV length could be less than 50 bp due to the presence of single nucleotide polymorphisms (SNPs), multiple-nucleotide polymorphisms (MNPs), or shorter SVs in the alternative alleles. The SVs in scaffolds 34 and 38 overlap with the genes *SDBH* and *TPM4*, respectively.

| Scaffold | Position | Comp 1 | $F_{ST}$ 1 | Comp 2 | $F_{ST}$ 2 | Comp 3 | $F_{ST}$ 3 | Comp 4 | $F_{ST}$ 4 | Comp 5 | $F_{ST}$ 5 | Genes | SV length | MeanSV length | SV type |
|---|---|---|---|---|---|---|---|---|---|---|---|---|---|---|---|
| 86 | 57,208 | HYPOX_CIN | 0.94 | HYPOX_HYPOCH | 0.78 | IBE_CIN | 0.80 | MEL_CIN | 0.91 | RUF_CIN | 0.84 | *TCF12* (279 bp) | 68 | 45.5 | ins, complex |
| 68 | 343,721 | HYPOX_HYPOCH | 0.78 | | | | | | | | | *TCF15* (90,370 bp) | 73, 74 | 73.9 | del |
| 34 | 6,433,081 | IBE_CIN | 0.75 | | | | | | | | | ***SDHB*** | 198-200, 202, 203, 206-210, 212, 215, 216 | 207.6 | ins |
| 225 | 9,842 | MEL_CIN | 0.81 | | | | | | | | | *LOC107049469* (3,388 bp) | 76 | 76.0 | complex |
| 24 | 644,469 | MEL_CIN | 0.78 | | | | | | | | | *PBX3* (10,496 bp) | 65 | 62.0 | ins, complex |
| 54 | 1,580,013 | MEL_CIN | 0.91 | MEL_HYPOCH | 0.83 | | | | | | | *C6orf132* (2,551 bp) | 163, 164, 178 | 155.8 | ins, complex |
| 20 | 456,801 | MEL_HYPOCH | 0.81 | | | | | | | | | *ZIC1* (69,988 bp) | 81 | 50.1 | complex |
| 38 | 4,673,394 | RUF_IBE | 0.80 | | | | | | | | | ***TPM4*** | 73 | 37.0 | ins |

**Table S11. SNP outliers overlapping coding regions with $F_{ST}$ > 0.75.** The table includes information on genomic coordinates, overlapping genes, comparisons, and $F_{ST}$ values for each comparison. Of the 118 SNPs overlapping coding regions only eight had $F_{ST}$ values above 0.75, four of which were found in multiple comparisons, generally involving *S. melanogaster*.

| Scaffold | Position | Gene | Comp 1 | $F_{ST}$ 1 | Comp 2 | $F_{ST}$ 2 | Comp 3 | $F_{ST}$ 3 |
|---|---|---|---|---|---|---|---|---|
| 10 | 6,204,110 | *HERC2* | HYPOX_RUF | 0.77 | | | | |
| 14 | 13,964,245 | *GPT2* | HYPOX_IBE | 0.80 | IBE_CIN | 0.76 | RUF_IBE | 0.75 |
| 3 | 9,964,755 | *ALB* | RUF_MEL | 0.85 | | | | |
| 3 | 9,965,640 | *ALB* | MEL_IBE | 0.83 | RUF_MEL | 0.79 | | |
| 3 | 9,968,037 | *ALB* | HYPOX_MEL | 0.76 | RUF_MEL | 0.75 | | |
| 3 | 9,971,007 | *ALB* | MEL_IBE | 0.84 | | | | |
| 7 | 25,297,310 | *TYRP1* | RUF_MEL | 0.85 | | | | |
| 7 | 25,300,749 | *TYRP1* | MEL_CIN | 0.86 | MEL_HYPOCH | 0.92 | | |

**Table S12. Information on long-read sequencing per individual.** The table shows the sequencing provider, yield in GB, HiFi mean read length in bp per sample, and genome coverage. Genome coverage is calculated as the yield in bp divided by the sum of the primary and alternate assemblies' lengths in bp (the lengths can be found in Table S16).

| Species | Sample ID | Museum ID / CECOAL ID | Locality | Province | Country | Sex | Sequencing | Yield (Gb) | HiFi average read length (bp) | Coverage (x) | NCBI Bioproject Accession (primary/alternate) |
|---|---|---|---|---|---|---|---|---|---|---|---|
| *S. cinnamomea* | CIN3121 | MACN 3121 | Iberá | Corrientes | Argentina | Male | Weill Cornell Medicine | 85.4 | 18,342 | 38.0 | PRJNA1223508/ PRJNA1223499 |
| *S. cinnamomea* | CIN3122 | MACN 3122 | Gualeguaychú | Entre Ríos | Argentina | Male | Weill Cornell Medicine | 93.4 | 19,637 | 41.3 | PRJNA1223507/ PRJNA1223498 |
| *S. hypochroma* | CRO3131 | MACN 3131 | Gualeguaychú | Entre Ríos | Argentina | Male | Weill Cornell Medicine | 77.3 | 15,981 | 34.2 | PRJNA1223506/ PRJNA1223497 |
| *S. hypoxantha* | HYPOXB009684 | Cecoal-Or-00114 | San Miguel | Corrientes | Argentina | Male | Novogene | 76.8 | 17,657 | 34.2 | PRJNA1223521/ PRJNA1223514 |
| *S. hypoxantha* | HYPOXB009666 | Cecoal-Or-00098 | San Miguel | Corrientes | Argentina | Female | Novogene | 62.4 | 17,684 | 28.9 | PRJNA1223522/ PRJNA1223515 |
| *S. hypoxantha* | HYPOXB009213 | Cecoal-Or-00119 | San Miguel | Corrientes | Argentina | Male | Novogene | 52.3 | 18,063 | 23.5 | PRJNA1223523/ PRJNA1223516 |
| *S. iberaensis* | IBEB009240 | Cecoal-Or-00077 | San Miguel | Corrientes | Argentina | Male | Weill Cornell Medicine | 88.0 | 13,663 | 39.1 | PRJNA1223520/ PRJNA1223513 |
| *S. iberaensis* | IBEB009699 | Cecoal-Or-00058 | San Miguel | Corrientes | Argentina | Male | Novogene | 65.4 | 17,366 | 29.2 | PRJNA1223517/ PRJNA1223510 |
| *S. iberaensis* | IBEB009696 | Cecoal-Or-00055 | San Miguel | Corrientes | Argentina | Male | Novogene | 61.6 | 17,522 | 27.3 | PRJNA1223518/ PRJNA1223511 |
| *S. iberaensis* | IBEB009632 | Cecoal-Or-00042 | San Miguel | Corrientes | Argentina | Female | Novogene | 36.7 | 16,468 | 16.7 | PRJNA1223519/ PRJNA1223512 |
| *S. palustris* | PAL3118 | MACN 3118 | Iberá | Corrientes | Argentina | Male | Weill Cornell Medicine | 75.9 | 11,376 | 34.2 | PRJNA1223504/ PRJNA1223495 |
| *S. palustris* | PAL3117 | MACN 3117 | Gualeguaychú | Entre Ríos | Argentina | Male | Weill Cornell Medicine | 81.7 | 12,563 | 36.6 | PRJNA1223505/ PRJNA1223496 |
| *S. palustris* | PAL3372 | MACN 3372 | Gualeguaychú | Entre Ríos | Argentina | Male | Weill Cornell Medicine | 82.1 | 12,463 | 36.7 | PRJNA1223503/ PRJNA1223494 |
| *S. pileata* | PIL3664 | KUNHM 3664 | San Rafael National Park | Itapuá | Paraguay | Male | Weill Cornell Medicine | 47.4 | 16,255 | 21.5 | PRJNA1223502/ PRJNA1223493 |
| *S. ruficollis* | RUF3129 | MACN 3129 | Gualeguaychú | Entre Ríos | Argentina | Male | Weill Cornell Medicine | 89.0 | 12,816 | 39.4 | PRJNA1223500/ PRJNA1223491 |
| *S. ruficollis* | RUF3128 | MACN 3128 | Gualeguaychú | Entre Ríos | Argentina | Male | Weill Cornell Medicine | 78.1 | 11,695 | 35.0 | PRJNA1223501/ PRJNA1223492 |
| *MEAN* | | | | | | | | 72.1 | 15,597 | 32.2 | |

**Table S13. Information for the 121 whole genome sequences obtained from previous studies.** The table provides information on species identity and the number of individuals available. The Accession numbers are all part of the NCBI BioProject PRJNA382416. The *S. hypochroma* sample marked with an asterisk (*) was also sequenced for this study (detailed in Table S14), and the data from both sequencing runs were combined.

| Species | #Ind. | Biosample Accession numbers | SRA Accession numbers |
|---|---|---|---|
| *S. bouvreuil* | 4 | SAMN19189818 - SAMN19189821 | SRX10889609, SRX10889610, SRX10889618, SRX10889619 |
| *S. cinnamomea* | 3 | SAMN19189822 - SAMN19189824 | SRX10889620 - SRX10889622 |
| *S. hypochroma* | 3* | SAMN19189825* - SAMN19189827 | SRX10889623* - SRX10889625 |
| *S. hypoxantha* | 28 | SAMN16367250 - SAMN16367265, SAMN06756684 -SAMN06756695 | SRS7478216 - SRS7478223, SRS7478247 - SRS7478252, SRS7478239, SRS7478228, SRS2123732 - SRS2123741 |
| *S. iberaensis* | 21 | SAMN16367266 - SAMN16367286 | SRX9245538 - SRX9245541, SRX9245543- SRX9245552, SRX9245554- SRX9245560. |
| *S. melanogaster* | 12 | SAMN06756696 - SAMN06756707 | SRX2736334 - SRX2736345 |
| *S. nigrorufa* | 12 | SAMN06756720 - SAMN06756731 | SRX2736310 - SRX2736321 |
| *S. palustris* | 12 | SAMN06756708 - SAMN06756719 | SRX2736322 - SRX2736333 |
| *S. pileata* | 12 | SAMN06756672 - SAMN06756683 | SRX2736358 - SRX2736369 |
| *S. ruficollis* | 14 | SAMN19220317 - SAMN19220327, SAMN19189828 - SAMN19189830 | SRX10897553 - SRX10897563, SRX10889611 - SRX10889613 |

**Table S14. Information for the 41 whole genome sequences obtained via short-read sequencing for this study.** The table shows details for species and individuals, accession numbers, location, sex, sequencing provider, total amount of raw reads, raw data in Gb (Raw reads*sequence length), Q30: (Base count of Phred value > 30) / (Total base count) and GC content. The sample ID marked with an asterisk (*) was previously sequenced. All sequencing was performed using the Illumina NovaSeq X platform with 2×150 Paired End reads.

| Species | Sample ID | Accession number | Locality | Province | Country | Sex | Sequencing | Raw Reads | Raw data (Gb) | Q30(%) | GC(%) |
|---|---|---|---|---|---|---|---|---|---|---|---|
| *S. bouvreuil* | BOU4243 | SRR32671667 | Lagoa Da Confusao | Tocantins | Brazil | male | Novogene | 97,316,352 | 14.6 | 93.1 | 42.5 |
| *S. bouvreuil* | BOU4246 | SRR32671666 | Lagoa Da Confusao | Tocantins | Brazil | male | Novogene | 106,674,336 | 16 | 93.0 | 42.6 |
| *S. bouvreuil* | BOU4252 | SRR32671659 | Roda Velha | Bahia | Brazil | male | Novogene | 101,179,798 | 15.2 | 93.4 | 42.5 |
| *S. bouvreuil* | BOU4368 | SRR32671658 | Campos do Goytacazes | Rio de Janeiro | Brazil | male | Novogene | 79,819,314 | 12 | 93.2 | 42.7 |
| *S. bouvreuil* | BOU4906 | SRR32671657 | Britania | Goias | Brazil | male | Novogene | 85,526,002 | 12.8 | 93.1 | 43.5 |
| *S. bouvreuil* | BOU4907 | SRR32671656 | Lagoa Da Confusao | Tocantins | Brazil | male | Novogene | 93,715,250 | 14.1 | 93.5 | 43.4 |
| *S. bouvreuil* | BOU4912 | SRR32671655 | Coromandel | Minas Gerais | Brazil | male | Novogene | 90,932,554 | 13.6 | 92.9 | 43.2 |
| *S. bouvreuil* | BOU4915 | SRR32671654 | Armação dos Búzios | Rio de Janeiro | Brazil | male | Novogene | 82,911,476 | 12.4 | 93.6 | 43.8 |
| *S. bouvreuil* | BOU4916 | SRR32671653 | Mogi das Cruzes | Sao Paulo | Brazil | male | Novogene | 103,482,012 | 15.5 | 93.2 | 43.2 |
| *S. bouvreuil* | BOU4917 | SRR32671652 | Mogi das Cruzes | Sao Paulo | Brazil | male | Novogene | 70,787,128 | 10.6 | 93.1 | 42.5 |
| *S. bouvreuil* | BOU4918 | SRR32671665 | Mogi das Cruzes | Sao Paulo | Brazil | male | Novogene | 78,689,752 | 11.8 | 93.1 | 43.4 |
| *S. cinnamomea* | cin_A09741 | SRR32725266 | Santo Tome | Corrientes | Argentina | male | BRC-Genomics | 223,006,646 | 38.3 | 95.3 | 42.1 |
| *S. cinnamomea* | cin_A09742 | SRR32725265 | Santo Tome | Corrientes | Argentina | male | BRC-Genomics | 230,073,396 | 39.5 | 95.4 | 42.5 |
| *S. cinnamomea* | cin_A09743 | SRR32725264 | Santo Tome | Corrientes | Argentina | male | BRC-Genomics | 227,689,736 | 39.1 | 94.1 | 41.8 |
| *S. cinnamomea* | cin_A09752 | SRR32725263 | Galarza | Corrientes | Argentina | male | BRC-Genomics | 204,052,986 | 35 | 94.5 | 42.2 |
| *S. cinnamomea* | cin_A09763 | SRR32725611 | Santo Tome | Corrientes | Argentina | male | BRC-Genomics | 233,336,232 | 40.0 | 94.9 | 42.4 |
| *S. cinnamomea* | cin_A09764 | SRR32725610 | Santo Tome | Corrientes | Argentina | male | BRC-Genomics | 198,073,216 | 34.0 | 94.7 | 42 |
| *S. cinnamomea* | cin_A09766 | SRR32725609 | Galarza | Corrientes | Argentina | male | BRC-Genomics | 206,712,266 | 35.5 | 95.2 | 42.1 |
| *S. cinnamomea* | cin_A09773 | SRR32725608 | Tres Cerros | Corrientes | Argentina | male | BRC-Genomics | 210,869,394 | 36.2 | 94.6 | 41.9 |
| *S. cinnamomea* | cin_B009069 | SRR32671662 | Pelegrini | Corrientes | Argentina | male | Novogene | 89,851,486 | 13.5 | 92.3 | 42 |
| *S. cinnamomea* | cin_B009147 | SRR32671661 | Pelegrini | Corrientes | Argentina | male | Novogene | 92,600,758 | 13.9 | 92.8 | 42.4 |

| | | | | | | | | | | | |
|---|---|---|---|---|---|---|---|---|---|---|---|
| *S. cinnamomea* | cin_B009149 | SRR32671660 | Pelegrini | Corrientes | Argentina | male | Novogene | 84,776,310 | 12.7 | 93.6 | 43.1 |
| *S. cinnamomea* | cin_C110481 | SRR32727699 | Quarai | Rio Grande do Sul | Brazil | male | BRC-Genomics | 224,457,988 | 38.5 | 95.1 | 43 |
| *S. cinnamomea* | cin_C110486 | SRR32727702 | Quarai | Rio Grande do Sul | Brazil | male | BRC-Genomics | 186,084,128 | 31.9 | 94.9 | 42.1 |
| *S. cinnamomea* | cin_C111768 | SRR32727701 | Manoel Viana | Rio Grande do Sul | Brazil | male | BRC-Genomics | 194,384,482 | 33.4 | 93.3 | 42.6 |
| *S. cinnamomea* | cin_C79910 | SRR32727700 | Manoel Viana | Rio Grande do Sul | Brazil | male | BRC-Genomics | 203,840,606 | 35.0 | 94.7 | 44.8 |
| *S. hypochroma* | hypoch_A09710 | SRR32698892 | Loreto | Corrientes | Argentina | male | BRC-Genomics | 240,342,440 | 41.2 | 94.7 | 42 |
| *S. hypochroma* | hypoch_A09711 | SRR32698891 | Loreto | Corrientes | Argentina | male | BRC-Genomics | 197,437,572 | 33.9 | 95.1 | 42 |
| *S. hypochroma* | hypoch_A09712 | SRR32698890 | Loreto | Corrientes | Argentina | male | BRC-Genomics | 200,215,156 | 34.4 | 95.5 | 42.2 |
| *S. hypochroma* | hypoch_A09713 | SRR32698889 | Loreto | Corrientes | Argentina | male | BRC-Genomics | 226,502,064 | 38.9 | 94.9 | 41.9 |
| *S. hypochroma* | hypoch_A09714 | SRR32698888 | Loreto | Corrientes | Argentina | male | BRC-Genomics | 221,947,662 | 38.1 | 94.3 | 41.8 |
| *S. hypochroma* | hypoch_A09715 | SRR32698887 | Loreto | Corrientes | Argentina | male | BRC-Genomics | 244,048,860 | 41.9 | 95 | 41.9 |
| *S. hypochroma* | hypoch_A09716 | SRR32698886 | Loreto | Corrientes | Argentina | male | BRC-Genomics | 239,839,230 | 41.2 | 95 | 42.2 |
| *S. hypochroma* | hypoch_A09718 | SRR32698885 | Loreto | Corrientes | Argentina | male | BRC-Genomics | 190,348,992 | 32.7 | 94.3 | 42 |
| *S. hypochroma* | hypoch_A09720 | SRR32738498 | Loreto | Corrientes | Argentina | male | BRC-Genomics | 213,342,884 | 36.6 | 94.7 | 41.8 |
| *S. hypochroma* | hypoch_A09721 | SRR32738497 | Loreto | Corrientes | Argentina | male | BRC-Genomics | 215,920,110 | 37.0 | 94.6 | 42.2 |
| *S. hypochroma* | hypoch_A09754 | SRR32738496 | Loreto | Corrientes | Argentina | male | BRC-Genomics | 200,903,430 | 34.5 | 94.5 | 42 |
| *S. hypochroma* | hypoch_A09801 | SRR32738495 | Loreto | Corrientes | Argentina | male | BRC-Genomics | 217,820,672 | 37.4 | 95.4 | 42.3 |
| *S. hypochroma* | hypoch_B009624* | SRR32331697 | San Miguel | Corrientes | Argentina | male | Novogene | 96,346,226 | 14.5 | 93.7 | 44.2 |
| *S. hypochroma* | hypoch_B009631 | SRR32671664 | San Miguel | Corrientes | Argentina | male | Novogene | 91,888,198 | 13.8 | 93.3 | 42.9 |
| *S. hypochroma* | hypoch_B009636 | SRR32671663 | Loreto | Corrientes | Argentina | male | Novogene | 90,924,884 | 13.6 | 92.4 | 43.1 |

**Table S15. Statistics for individual genome assemblies obtained from GenomeScope.** GenomeScope is a tool used to estimate the genome size and assess genome assembly quality based on k-mer frequency analysis. The metrics provided help evaluate both the completeness and quality of the genome assembly. The table includes details on k-mer size used, as well as the maximum and minimum values for the following estimated metrics: percentage of heterozygosity, haploid genome size, repeat and unique sequence lengths (in base pairs), percentage of model fit, and percentage of read error rate. The percentage of model fit represents the percentage of the genomic data that fits the expected model during genome assembly. A higher percentage indicates better alignment between the observed data and the model, reflecting a more accurate genome assembly.

| Sample ID | Species | Sex | Kmer | Heterozygosity (%) | | Genome Haploid Length (bp) | | Genome Repeat Length (bp) | | Genome Unique Length (bp) | | Model Fit (%) | | Read Error Rate (%) | |
|---|---|---|---|---|---|---|---|---|---|---|---|---|---|---|---|
| | | | | min | max | min | max | min | max | min | max | min | max | min | max |
| CIN3121 | *S. cinnamomea* | Male | 21 | 1.20 | 1.21 | 959,414,918 | 961,112,849 | 26,390,237 | 26,436,941 | 933,024,681 | 934,675,908 | 93.8 | 94.3 | 0.24 | 0.24 |
| CIN3122 | *S. cinnamomea* | Male | 21 | 1.22 | 1.23 | 963,858,794 | 965,347,625 | 34,015,080 | 34,067,622 | 929,843,714 | 931,280,003 | 94.9 | 95.5 | 0.23 | 0.23 |
| CRO3131 | *S. hypochroma* | Male | 19 | 1.01 | 1.10 | 1,053,974,948 | 1,054,281,639 | 161,339,322 | 161,386,269 | 892,635,627 | 892,895,370 | 95.9 | 96.9 | 0.16 | 0.16 |
| HYPOXB009213 | *S. hypoxantha* | Male | 19 | 1.22 | 1.22 | 1,011,216,292 | 1,011,652,286 | 135,740,805 | 135,799,331 | 875,475,487 | 875,852,955 | 95.8 | 96.7 | 0.19 | 0.19 |
| HYPOXB009666 | *S. hypoxantha* | Female | 21 | 1.27 | 1.28 | 987,604,331 | 988,602,748 | 63,519,458 | 63,583,672 | 924,084,873 | 925,019,076 | 98.0 | 98.7 | 0.25 | 0.25 |
| HYPOXB009684 | *S. hypoxantha* | Male | 21 | 1.22 | 1.22 | 997,807,174 | 998,752,059 | 58,632,524 | 58,688,047 | 939,174,650 | 940,064,012 | 97.6 | 98.2 | 0.19 | 0.19 |
| IBEB009240 | *S. iberaensis* | Male | 21 | 1.12 | 1.13 | 984,604,023 | 986,065,667 | 34,623,946 | 34,675,345 | 949,980,077 | 951,390,321 | 94.5 | 95.0 | 0.17 | 0.17 |
| IBEB009632 | *S. iberaensis* | Female | 21 | 1.41 | 1.42 | 943,896,397 | 944,744,794 | 50,649,757 | 50,695,282 | 893,246,641 | 894,049,512 | 97.0 | 97.7 | 0.31 | 0.31 |
| IBEB009696 | *S. iberaensis* | Male | 21 | 1.15 | 1.15 | 1,028,504,691 | 1,029,239,642 | 78,268,883 | 78,324,813 | 950,235,807 | 950,914,829 | 98.6 | 99.3 | 0.16 | 0.16 |
| IBEB009699 | *S. iberaensis* | Male | 19 | 1.13 | 1.14 | 1,018,500,633 | 1,018,935,170 | 139,731,729 | 139,791,345 | 878,768,903 | 879,143,825 | 94.7 | 95.6 | 0.17 | 0.17 |
| PAL3117 | *S. palustris* | Male | 21 | 1.18 | 1.19 | 1,008,376,723 | 1,008,890,917 | 67,839,371 | 67,873,964 | 940,537,352 | 941,016,952 | 98.4 | 99.1 | 0.17 | 0.17 |
| PAL3118 | *S. palustris* | Male | 21 | 1.18 | 1.20 | 967,702,866 | 969,441,095 | 26,487,817 | 26,535,396 | 941,215,049 | 942,905,699 | 94.0 | 94.5 | 0.16 | 0.16 |
| PAL3372 | *S. palustris* | Male | 21 | 1.17 | 1.18 | 988,033,924 | 978,324,938 | 40,248,601 | 40,301,748 | 936,785,322 | 938,023,154 | 95.5 | 96.0 | 0.22 | 0.22 |
| PIL3664 | *S. pileata* | Male | 19 | 1.39 | 1.30 | 968,769,637 | 969,258,522 | 119,748,507 | 119,808,937 | 849,021,130 | 849,449,584 | 95.7 | 96.5 | 0.18 | 0.18 |
| RUF3128 | *S. ruficollis* | Male | 21 | 1.17 | 1.19 | 968,785,374 | 970,213,715 | 32,335,377 | 32,383,051 | 936,449,997 | 937,830,665 | 94.7 | 95.2 | 0.22 | 0.22 |
| RUF3129 | *S. ruficollis* | Male | 21 | 1.12 | 1.13 | 984,604,023 | 986,065,667 | 34,623,946 | 34,675,345 | 949,980,077 | 951,390,321 | 94.5 | 95.0 | 0.17 | 0.17 |

**Table S16. Reference genome assembly statistics.** Statistics obtained with assembly-stats and Merqury for the primary and alternate assemblies after purging with purge_dups. The table includes total assembly length, the number of scaffolds (#scaff), the average and longest scaffold lengths, and various summary statistics of scaffold length distribution (N50, L50, N90, L90), as well as the QV score from Merqury (QV Merqury), which reflects the overall quality of the assembly. A QV score of 60 corresponds to 99.9999% accuracy with an error rate of fewer than 1 in 1 million bases.

| Assembly | Total length | #Scaff | Average length | Longest Scaffold | N50 | L50 | N90 | L90 | QV Merqury |
|---|---|---|---|---|---|---|---|---|---|
| RUF3129_PRI | 1,146,290,513 | 267 | 4,293,223 | 94,742,372 | 40,913,997 | 8 | 8,040,665 | 31 | 62.08 |
| RUF3129_ALT | 1,110,450,307 | 920 | 1,207,011 | 17,176,647 | 3,392,195 | 90 | 928,611 | 324 | 63.55 |
| IBEB009240_PRI | 1,149,040,470 | 253 | 4,541,662 | 118,976,643 | 47,202,461 | 8 | 8,796,480 | 30 | 62.85 |
| IBEB009240_ALT | 1,099,709,422 | 1,015 | 1,083,458 | 17,624,126 | 3,183,133 | 99 | 770,869 | 369 | 64.48 |
| PAL3118_PRI | 1,128,643,828 | 438 | 2,576,812 | 69,909,865 | 18,321,296 | 19 | 2,166,640 | 89 | 61.19 |
| PAL3118_ALT | 1,088,367,715 | 1,226 | 887,739 | 15,200,516 | 2,280,547 | 127 | 524,445 | 496 | 63.34 |
| PAL3117_PRI | 1,136,960,334 | 369 | 3,081,193 | 55,109,529 | 19,073,713 | 20 | 3,292,957 | 71 | 61.78 |
| PAL3117_ALT | 1,097,376,711 | 1,035 | 1,060,267 | 12,242,520 | 2,750,673 | 113 | 649,217 | 430 | 64.16 |
| PAL3372_PRI | 1,128,716,638 | 358 | 3,152,840 | 90,165,078 | 19,783,407 | 17 | 3,831,394 | 61 | 61.53 |
| PAL3372_ALT | 1,105,521,224 | 1,070 | 1,033,197 | 10,443,939 | 2,550,160 | 122 | 677,950 | 442 | 64.06 |
| RUF3128_PRI | 1,131,988,263 | 409 | 2,767,697 | 67,689,051 | 16,874,569 | 20 | 2,448,741 | 76 | 61.77 |
| RUF3128_ALT | 1,101,792,386 | 1,143 | 963,948 | 13,169,795 | 2,583,021 | 122 | 593,556 | 447 | 63.78 |
| HYPOXB009213_PRI | 1,135,591,397 | 252 | 4,506,315 | 95,746,362 | 34,667,066 | 10 | 3,681,269 | 46 | 60.56 |
| HYPOXB009213_ALT | 1,091,086,072 | 956 | 1,141,303 | 19,255,472 | 3,120,974 | 97 | 716,173 | 374 | 62.5 |
| IBEB009632_PRI | 1,125,838,309 | 423 | 2,661,556 | 77,260,784 | 20,709,386 | 13 | 1,998,517 | 74 | 58.61 |
| IBEB009632_ALT | 1,070,377,057 | 1,413 | 757,521 | 14,550,535 | 2,166,875 | 136 | 428,325 | 545 | 60.75 |
| HYPOXB009666_PRI | 1,147,486,443 | 530 | 2,165,069 | 35,250,551 | 10,988,915 | 33 | 1,440,130 | 151 | 60.1 |
| HYPOXB009666_ALT | 1,013,291,307 | 1,267 | 799,756 | 15,803,046 | 1,927,616 | 149 | 450,819 | 569 | 62.41 |
| HYPOXB009684_PRI | 1,144,606,514 | 236 | 4,850,028 | 134,417,494 | 41,767,700 | 8 | 6,615,089 | 38 | 60.6 |
| HYPOXB009684_ALT | 1,102,220,544 | 1,183 | 931,716 | 12,196,746 | 2,456,997 | 130 | 614,682 | 469 | 61.88 |
| IBEB009696_PRI | 1,150,342,677 | 178 | 6,462,599 | 153,566,929 | 37,263,916 | 8 | 6,701,677 | 35 | 62.04 |
| IBEB009696_ALT | 1,104,331,787 | 904 | 1,221,606 | 19,447,870 | 3,679,808 | 92 | 807,608 | 338 | 63.21 |
| IBEB009699_PRI | 1,148,684,330 | 145 | 7,921,961 | 92,772,356 | 42,051,479 | 9 | 9,227,367 | 32 | 62.51 |
| IBEB009699_ALT | 1,094,216,911 | 922 | 1,186,786 | 13,399,975 | 3,540,419 | 95 | 797,998 | 339 | 63.63 |
| CIN3121_PRI | 1,144,075,825 | 250 | 4,576,303 | 113,592,198 | 57,159,568 | 8 | 5,206,629 | 35 | 62.46 |
| CIN3121_ALT | 1,102,792,574 | 1,008 | 1,094,040 | 15,707,152 | 2,783,624 | 117 | 783,564 | 393 | 63.85 |
| CIN3122_PRI | 1,145,943,948 | 199 | 5,758,512 | 192,003,346 | 37,151,129 | 8 | 7,086,252 | 31 | 62.7 |
| CIN3122_ALT | 1,111,285,234 | 979 | 1,135,123 | 17,206,311 | 3,425,069 | 95 | 799,368 | 345 | 64.28 |
| CRO3131_PRI | 1,157,215,112 | 239 | 4,841,904 | 170,088,522 | 32,602,669 | 10 | 6,275,244 | 37 | 61.96 |
| CRO3131_ALT | 1,104,442,460 | 868 | 1,272,399 | 17,744,939 | 3,240,927 | 93 | 866,114 | 338 | 63.83 |
| PIL3664_PRI | 1,122,967,458 | 507 | 2,214,926 | 83,002,095 | 23,471,106 | 13 | 1,884,327 | 73 | 59.87 |
| PIL3664_ALT | 1,079,294,003 | 1,300 | 830,226 | 13,487,222 | 2,844,671 | 115 | 456,801 | 479 | 62.29 |

**Table S17. Structural variants detected using long-read sequencing data for each individual.** Number of variants found by the three SV callers (PBSV, Sniffles, and Svim_asm) and the total number of variants recovered by SURVIVOR, supported by all three SV callers, categorized by SV type. SV type abbreviations are: deletions (DEL), duplications (DUP), insertions (INS), inversions (INV), and translocations (TRA). PBSV was the caller that recovered the most variants, followed by Sniffles, and finally Svim_asm. Across all assemblies, the SV callers and SURVIVOR recovered a similar number of variants, which were also distributed similarly in terms of SV types.

| Individual | PIL3664 | CIN3121 | CIN3122 | CRO3131 | HYPOX B009666 | HYPOX B009213 | IBE B009696 | IBE B009632 | RUF3129 | RUF3128 | PAL3117 | PAL3372 | IBE B009699 | IBE B009240 | PAL3118 | MEAN | SD |
|---|---|---|---|---|---|---|---|---|---|---|---|---|---|---|---|---|---|
| PBSV | 229,783 | 247,801 | 250,676 | 253,351 | 247,404 | 240,703 | 245,478 | 233,058 | 248,698 | 246,657 | 247,949 | 246,598 | 249,120 | 248,086 | 229,430 | 244,319.5 | 7,550.3 |
| Sniffles | 108,592 | 110,854 | 111,372 | 111,015 | 109,768 | 110,757 | 110,803 | 109,979 | 110,194 | 108,307 | 109,019 | 108,389 | 111,489 | 109,619 | 108,216 | 109,891.5 | 1,156.6 |
| SVIM | 92,356 | 99,300 | 100,572 | 100,169 | 93,301 | 96,795 | 99,628 | 92,952 | 101,412 | 97,901 | 97,307 | 97,693 | 98,311 | 100,759 | 96,487 | 97,662.9 | 2,891.5 |
| SURVIVOR | 53,763 | 56,175 | 56,560 | 56,469 | 54,286 | 55,810 | 56,038 | 54,409 | 56,097 | 55,481 | 55,447 | 55,457 | 55,811 | 56,163 | 54,072 | 55,469.2 | 906.6 |
| DEL | 27,113 | 28,221 | 28,288 | 28,302 | 27,115 | 27,817 | 28,307 | 27,260 | 28,072 | 27,803 | 28,212 | 27983 | 28,254 | 28,452 | 29,006 | 28,013.7 | 523.4 |
| DUP | 5 | 2 | 3 | 2 | 0 | 4 | 2 | 4 | 2 | 6 | 5 | 5 | 4 | 4 | 2 | 3.3 | 1.6 |
| INS | 24,047 | 25,730 | 25,846 | 25,973 | 24,832 | 25,288 | 25,224 | 24,335 | 25,748 | 25,417 | 25,177 | 25,181 | 25,184 | 25,398 | 24,286 | 25,177.7 | 581.4 |
| INV | 16 | 23 | 23 | 21 | 15 | 20 | 17 | 18 | 21 | 24 | 21 | 16 | 18 | 16 | 25 | 19.6 | 3.3 |
| TRA | 2 | 1 | 0 | 2 | 2 | 4 | 1 | 1 | 3 | 6 | 3 | 2 | 0 | 1 | 0 | 1.9 | 1.6 |

**Table S18. Information on short-read data for each individual.** The table includes individual and species information, number of variants and coverage after mapping against the pangenome. The individuals marked with an asterisk did not pass the 4× coverage filter, the ones in bold were sequenced for this study and the individuals highlighted in green were used to calculate the mean mapping quality in 50 kb windows along the pangenome. For the sample marked with an (#) we combined sequences from two runs. Out of a total of 161 individuals, 127 passed the 4X coverage filter and were included in the final dataset.

| # | Individual | Species | #Variants | VCFs coverage |
|---|---|---|---|---|
| 1 | **bou_4243** | *S. bouvreuil* | 34,611,785 | 9.2 |
| 2 | **bou_4246** | *S. bouvreuil* | 34,610,526 | 9.7 |
| 3 | **bou_4252** | *S. bouvreuil* | 34,611,414 | 9.4 |
| 4 | **bou_4368** | *S. bouvreuil* | 34,614,852 | 7.8 |
| 5 | **bou_4906** | *S. bouvreuil* | 34,609,422 | 6.5 |
| 6 | **bou_4907** | *S. bouvreuil* | 34,608,940 | 7.7 |
| 7 | **bou_4912** | *S. bouvreuil* | 34,608,775 | 8.2 |
| 8 | **bou_4915** | *S. bouvreuil* | 34,616,947 | 7.3 |
| 9 | **bou_4916** | *S. bouvreuil* | 34,610,435 | 9.4 |
| 10 | **bou_4917** | *S. bouvreuil* | 34,615,525 | 5.5 |
| 11 | **bou_4918** | *S. bouvreuil* | 34,612,143 | 7.0 |
| 12 | bou_L3_13 | *S. bouvreuil* | 34,634,594 | 5.2 |
| 13 | bou_L3_14 | *S. bouvreuil* | 34,638,334 | 4.7 |
| 14 | bou_L3_15 | *S. bouvreuil* | 34,621,801 | 8.7 |
| 15 | bou_O6 | *S. bouvreuil* | 34,633,631 | 4.7 |
| 16 | **cin_A09741** | *S. cinnamomea* | 34,606,910 | 24.4 |
| 17 | **cin_A09742** | *S. cinnamomea* | 34,606,954 | 25.3 |
| 18 | **cin_A09743** | *S. cinnamomea* | 34,609,163 | 25.4 |
| 19 | **cin_A09752** | *S. cinnamomea* | 34,609,252 | 23.0 |
| 20 | **cin_A09763** | *S. cinnamomea* | 34,609,748 | 25.5 |
| 21 | **cin_A09764** | *S. cinnamomea* | 34,608,607 | 19.1 |
| 22 | **cin_A09766** | *S. cinnamomea* | 34,607,412 | 22.8 |
| 23 | **cin_A09773** | *S. cinnamomea* | 34,609,436 | 23.4 |
| 24 | **cin_B009069** | *S. cinnamomea* | 34,614,732 | 8.0 |
| 25 | **cin_B009147** | *S. cinnamomea* | 34,610,818 | 9.5 |
| 26 | **cin_B009149** | *S. cinnamomea* | 34,623,565 | 6.9 |
| 27 | **cin_C110481** | *S. cinnamomea* | 34,608,195 | 23.0 |
| 28 | **cin_C110486** | *S. cinnamomea* | 34,608,094 | 20.6 |
| 29 | **cin_C111768** | *S. cinnamomea* | 34,607,659 | 21.8 |
| 30 | **cin_C79910** | *S. cinnamomea* | 34,605,718 | 22.0 |
| 31 | cin_L3_16* | *S. cinnamomea* | 34,652,888 | 3.3 |
| 32 | cin_L3_18* | *S. cinnamomea* | 34,651,243 | 3.3 |

| | | | | |
|---|---|---|---|---|
| 33 | cin_L3_19* | *S. cinnamomea* | 34,652,122 | 3.9 |
| 34 | **hypoch_A09710** | *S. hypochroma* | 34,607,123 | 26.6 |
| 35 | **hypoch_A09711** | *S. hypochroma* | 34,607,745 | 22.0 |
| 36 | **hypoch_A09712** | *S. hypochroma* | 34,606,606 | 21.3 |
| 37 | **hypoch_A09713** | *S. hypochroma* | 34,608,556 | 25.4 |
| 38 | **hypoch_A09714** | *S. hypochroma* | 34,610,622 | 25.2 |
| 39 | **hypoch_A09715** | *S. hypochroma* | 34,608,200 | 27.5 |
| 40 | **hypoch_A09716** | *S. hypochroma* | 34,605,729 | 26.3 |
| 41 | **hypoch_A09718** | *S. hypochroma* | 34,610,388 | 21.3 |
| 42 | **hypoch_A09720** | *S. hypochroma* | 34,609,649 | 24.0 |
| 43 | **hypoch_A09721** | *S. hypochroma* | 34,609,404 | 24.2 |
| 44 | **hypoch_A09754** | *S. hypochroma* | 34,608,549 | 22.4 |
| 45 | **hypoch_A09801** | *S. hypochroma* | 34,606,304 | 23.6 |
| 46 | **hypoch_B009624_O3**[#] | *S. hypochroma* | 34,607,860 | 11.3 |
| 47 | **hypoch_B009631** | *S. hypochroma* | 34,613,995 | 9.0 |
| 48 | **hypoch_B009636** | *S. hypochroma* | 34,611,773 | 9.5 |
| 49 | hypoch_L3_20 | *S. hypochroma* | 34,616,803 | 8.3 |
| 50 | hypoch_L3_21 | *S. hypochroma* | 34,643,473 | 4.4 |
| 51 | hypox_L1_20* | *S. hypoxantha* | 34,659,434 | 2.2 |
| 52 | hypox_L1_21* | *S. hypoxantha* | 34,664,233 | 2.6 |
| 53 | hypox_L1_22* | *S. hypoxantha* | 34,665,895 | 3.0 |
| 54 | hypox_L1_23 | *S. hypoxantha* | 34,661,458 | 4.3 |
| 55 | hypox_L1_25 | *S. hypoxantha* | 34,652,538 | 4.0 |
| 56 | hypox_L1_27* | *S. hypoxantha* | 34,650,667 | 3.8 |
| 57 | hypox_L2_13 | *S. hypoxantha* | 34,646,900 | 4.2 |
| 58 | hypox_L2_14 | *S. hypoxantha* | 34,643,707 | 4.4 |
| 59 | hypox_L2_15 | *S. hypoxantha* | 34,645,807 | 4.3 |
| 60 | hypox_L2_16* | *S. hypoxantha* | 34,662,234 | 3.5 |
| 61 | hypox_L2_18 | *S. hypoxantha* | 34,638,956 | 4.1 |
| 62 | hypox_L2_19* | *S. hypoxantha* | 34,677,738 | 3.6 |
| 63 | hypox_SH10 | *S. hypoxantha* | 34,636,945 | 5.0 |
| 64 | hypox_SH11 | *S. hypoxantha* | 34,632,128 | 6.2 |
| 65 | hypox_SH12 | *S. hypoxantha* | 34,632,911 | 6.2 |
| 66 | hypox_SH15 | *S. hypoxantha* | 34,632,963 | 5.8 |
| 67 | hypox_SH16 | *S. hypoxantha* | 34,642,876 | 4.6 |
| 68 | hypox_SH21 | *S. hypoxantha* | 34,646,485 | 4.1 |
| 69 | hypox_SH22 | *S. hypoxantha* | 34,641,478 | 4.6 |
| 70 | hypox_SH23 | *S. hypoxantha* | 34,645,316 | 4.8 |
| 71 | hypox_SH24 | *S. hypoxantha* | 34,645,105 | 4.5 |
| 72 | hypox_SH28 | *S. hypoxantha* | 34,639,556 | 5.1 |

| | | | | |
|---|---|---|---|---|
| 73 | hypox_SH29 | *S. hypoxantha* | 34,639,336 | 5.3 |
| 74 | hypox_SH3 | *S. hypoxantha* | 34,651,754 | 4.1 |
| 75 | hypox_SH30 | *S. hypoxantha* | 34,636,887 | 6.4 |
| 76 | hypox_SH31 | *S. hypoxantha* | 34,633,725 | 6.0 |
| 77 | hypox_SH8 | *S. hypoxantha* | 34,643,426 | 5.0 |
| 78 | hypox_SH9 | *S. hypoxantha* | 34,636,772 | 4.9 |
| 79 | ibe_SH1 | *S. iberaensis* | 34,651,542 | 4.8 |
| 80 | ibe_SH13 | *S. iberaensis* | 34,638,008 | 5.1 |
| 81 | ibe_SH14 | *S. iberaensis* | 34,648,907 | 4.1 |
| 82 | ibe_SH17* | *S. iberaensis* | 34,658,523 | 3.5 |
| 83 | ibe_SH18 | *S. iberaensis* | 34,641,078 | 4.8 |
| 84 | ibe_SH19 | *S. iberaensis* | 34,633,979 | 5.2 |
| 85 | ibe_SH2 | *S. iberaensis* | 34,640,554 | 4.8 |
| 86 | ibe_SH20 | *S. iberaensis* | 34,648,765 | 4.2 |
| 87 | ibe_SH25 | *S. iberaensis* | 34,646,828 | 4.3 |
| 88 | ibe_SH26 | *S. iberaensis* | 34,642,383 | 5.4 |
| 89 | ibe_SH27* | *S. iberaensis* | 34,666,770 | 3.3 |
| 90 | ibe_SH32 | *S. iberaensis* | 34,645,878 | 4.5 |
| 91 | ibe_SH33 | *S. iberaensis* | 34,629,053 | 6.2 |
| 92 | ibe_SH34 | *S. iberaensis* | 34,637,523 | 5.4 |
| 93 | ibe_SH35 | *S. iberaensis* | 34,648,652 | 4.7 |
| 94 | ibe_SH36 | *S. iberaensis* | 34,641,655 | 4.9 |
| 95 | ibe_SH37 | *S. iberaensis* | 34,620,878 | 9.2 |
| 96 | ibe_SH4 | *S. iberaensis* | 34,637,316 | 5.1 |
| 97 | ibe_SH5 | *S. iberaensis* | 34,635,577 | 5.3 |
| 98 | ibe_SH6 | *S. iberaensis* | 34,647,554 | 4.4 |
| 99 | ibe_SH7 | *S. iberaensis* | 34,642,280 | 4.6 |
| 100 | mel_L1_13* | *S. melanogaster* | 34,661,006 | 3.7 |
| 101 | mel_L1_14* | *S. melanogaster* | 34,671,090 | 3.1 |
| 102 | mel_L1_15 | *S. melanogaster* | 34,638,855 | 5.3 |
| 103 | mel_L1_16 | *S. melanogaster* | 34,632,354 | 4.7 |
| 104 | mel_L1_18 | *S. melanogaster* | 34,651,079 | 4.4 |
| 105 | mel_L1_19* | *S. melanogaster* | 34,665,563 | 3.4 |
| 106 | mel_L2_10 | *S. melanogaster* | 34,645,622 | 4.4 |
| 107 | mel_L2_11 | *S. melanogaster* | 34,641,782 | 5.0 |
| 108 | mel_L2_12 | *S. melanogaster* | 34,646,801 | 5.0 |
| 109 | mel_L2_7 | *S. melanogaster* | 34,646,780 | 4.5 |
| 110 | mel_L2_8 | *S. melanogaster* | 34,653,427 | 4.6 |
| 111 | mel_L2_9 | *S. melanogaster* | 34,648,435 | 4.2 |
| 112 | nig_L1_1* | *S. nigrorufa* | 34,821,788 | 1.2 |

| | | | | |
|---|---|---|---|---|
| 113 | nig_L1_2* | *S. nigrorufa* | 34,694,581 | 2.0 |
| 114 | nig_L1_3* | *S. nigrorufa* | 34,656,957 | 3.0 |
| 115 | nig_L1_4* | *S. nigrorufa* | 34,846,843 | 1.1 |
| 116 | nig_L1_5* | *S. nigrorufa* | 34,666,766 | 2.9 |
| 117 | nig_L1_6* | *S. nigrorufa* | 34,653,193 | 2.8 |
| 118 | nig_L2_1* | *S. nigrorufa* | 34,662,472 | 3.6 |
| 119 | nig_L2_2 | *S. nigrorufa* | 34,646,540 | 4.4 |
| 120 | nig_L2_3* | *S. nigrorufa* | 34,658,126 | 3.8 |
| 121 | nig_L2_4 | *S. nigrorufa* | 34,641,625 | 4.5 |
| 122 | nig_L2_5 | *S. nigrorufa* | 34,639,278 | 4.1 |
| 123 | nig_L2_6 | *S. nigrorufa* | 34,641,351 | 4.7 |
| 124 | pal_L3_1 | *S. palustris* | 34,622,867 | 7.4 |
| 125 | pal_L3_10* | *S. palustris* | 34,670,447 | 2.7 |
| 126 | pal_L3_11* | *S. palustris* | 34,671,637 | 2.4 |
| 127 | pal_L3_12 | *S. palustris* | 34,640,874 | 4.0 |
| 128 | pal_L3_2 | *S. palustris* | 34,637,560 | 4.3 |
| 129 | pal_L3_3 | *S. palustris* | 34,631,747 | 5.5 |
| 130 | pal_L3_4 | *S. palustris* | 34,641,384 | 4.0 |
| 131 | pal_L3_5* | *S. palustris* | 34,644,047 | 3.9 |
| 132 | pal_L3_6* | *S. palustris* | 34,692,007 | 2.0 |
| 133 | pal_L3_7 | *S. palustris* | 34,621,385 | 7.8 |
| 134 | pal_L3_8 | *S. palustris* | 34,639,541 | 4.2 |
| 135 | pal_L3_9* | *S. palustris* | 34,697,695 | 2.0 |
| 136 | pil_L1_10* | *S. pileata* | 34,644,365 | 3.7 |
| 137 | pil_L1_11* | *S. pileata* | 34,669,733 | 3.2 |
| 138 | pil_L1_12* | *S. pileata* | 34,744,534 | 1.6 |
| 139 | pil_L1_7* | *S. pileata* | 34,662,748 | 3.6 |
| 140 | pil_L1_8* | *S. pileata* | 34,743,392 | 1.9 |
| 141 | pil_L1_9* | *S. pileata* | 34,701,945 | 2.5 |
| 142 | pil_L2_20 | *S. pileata* | 34,645,400 | 4.2 |
| 143 | pil_L2_21 | *S. pileata* | 34,635,788 | 4.3 |
| 144 | pil_L2_22 | *S. pileata* | 34,634,031 | 4.0 |
| 145 | pil_L2_23 | *S. pileata* | 34,634,314 | 4.5 |
| 146 | pil_L2_25 | *S. pileata* | 34,635,912 | 4.5 |
| 147 | pil_L2_27 | *S. pileata* | 34,638,370 | 4.5 |
| 148 | ruf_C1 | *S. ruficollis* | 34,639,288 | 5.8 |
| 149 | ruf_C10 | *S. ruficollis* | 34,666,635 | 4.8 |
| 150 | ruf_C11 | *S. ruficollis* | 34,654,551 | 4.0 |
| 151 | ruf_C12 | *S. ruficollis* | 34,643,748 | 4.9 |
| 152 | ruf_C13 | *S. ruficollis* | 34,654,337 | 4.0 |

| | | | | |
|---|---|---|---|---|
| 153 | ruf_C2 | *S. ruficollis* | 34,631,428 | 7.5 |
| 154 | ruf_C3 | *S. ruficollis* | 34,632,091 | 7.2 |
| 155 | ruf_C4 | *S. ruficollis* | 34,634,041 | 7.8 |
| 156 | ruf_C6 | *S. ruficollis* | 34,624,862 | 8.3 |
| 157 | ruf_C7 | *S. ruficollis* | 34,653,050 | 5.0 |
| 158 | ruf_C8 | *S. ruficollis* | 34,639,452 | 7.1 |
| 159 | ruf_L3_23 | *S. ruficollis* | 34,626,443 | 6.0 |
| 160 | ruf_L3_25* | *S. ruficollis* | 34,683,550 | 2.3 |
| 161 | ruf_L3_27 | *S. ruficollis* | 34,634,123 | 5.2 |

**Table S19. Summary of variant numbers and differences across VCF files used in GWAS and $F_{ST}$ analyses.** Includes SNPs, variants within and outside annotated repetitive elements and TEs, and SVs < and > 50 bp. The difference in the number of variants between the datasets is due to the exclusion of SVs with more than 254 alternative alleles in the GWAS dataset. SNPs+MNPs refers to single nucleotide polymorphisms and multi-nucleotide polymorphisms, while TEs_SNPs_SVs and noTEs_SNPs_SVs refer to variants (both SNPs and SVs) within and outside annotated repetitive elements and transposable elements, respectively.

| VCF File | GWAS | $F_{ST}$ | Difference |
|---|---|---|---|
| SNPs+MNPs | 27,875,982 | 27,875,982 | 0 |
| TEs_SNPs_SVs | 4,387,155 | 4,388,258 | 1,103 |
| noTEs_SNPs_SVs | 27,016,423 | 27,018,764 | 2,341 |
| SVs Long | 93,247 | 96,682 | 3,435 |
| SVs Short (indels) | 3,434,349 | 3,434,358 | 9 |
| SVs Total | 3,527,596 | 3,531,040 | 3,444 |

### Supplementary Figures

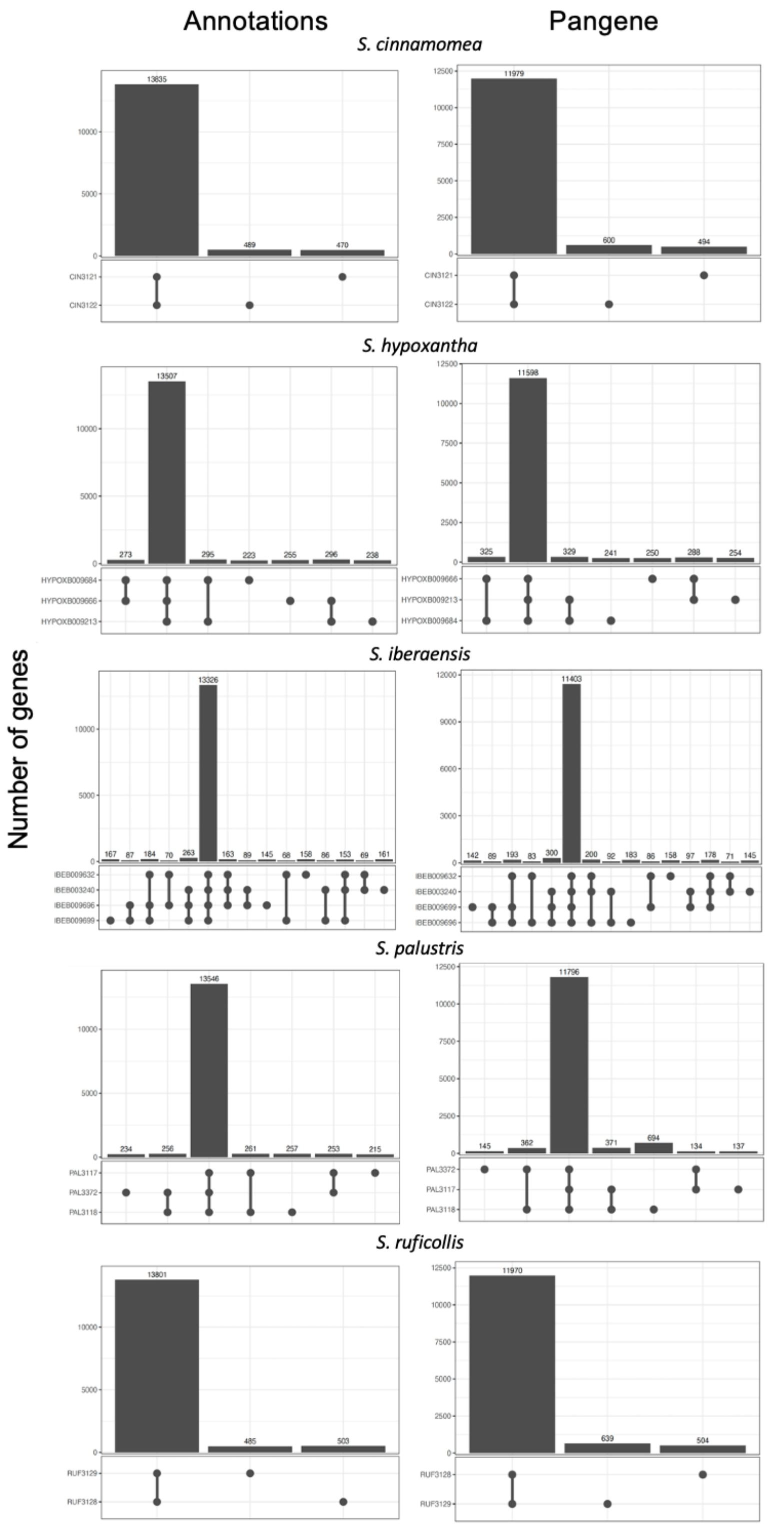

**Figure S1. Presence-Absence Variation Analysis (PAV) using annotated genes and Pangene per species.** We used two complementary approaches to assess gene-level variation across assemblies within species. First, we analyzed gene presence and absence using GFF-derived gene lists from the annotations. Second, we used Pangene with Miniprot to align protein-coding exons across assemblies. (A) Upset plots for the gene annotation (left) and Pangene (right) approaches, showing unique and shared genes among individuals within each species. Across all species, most genes are shared among all individuals or a subset of them, with only a small proportion unique to a single individual.

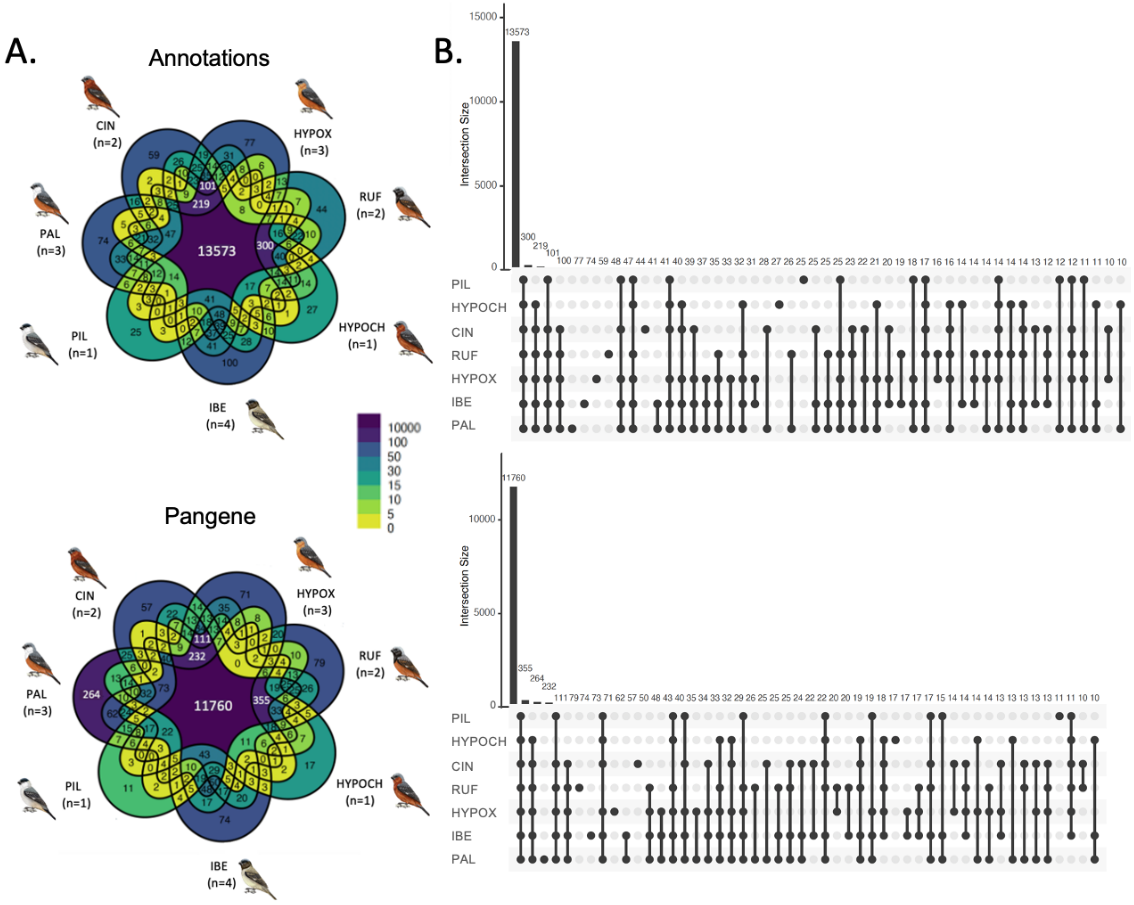

**Figure S2. Presence-Absence Variation (PAV) analysis using annotated genes and Pangene across all species.** We combined the individual lists of genes per species for the plots. (A) Venn diagrams for the gene annotation (top) and Pangene (bottom) approaches, illustrating unique genes per species and the overlap of genes among different species. The color gradient transitions from yellow to purple, indicates low to high gene counts. (B) Upset plots showing the number of genes shared across various combinations of species for the annotation (top) and Pangene (bottom) methods. Most genes are shared among all assemblies, while each method detects a small number of unique genes per species that do not overlap between the two approaches. Species names are shortened to the first three to six letters.

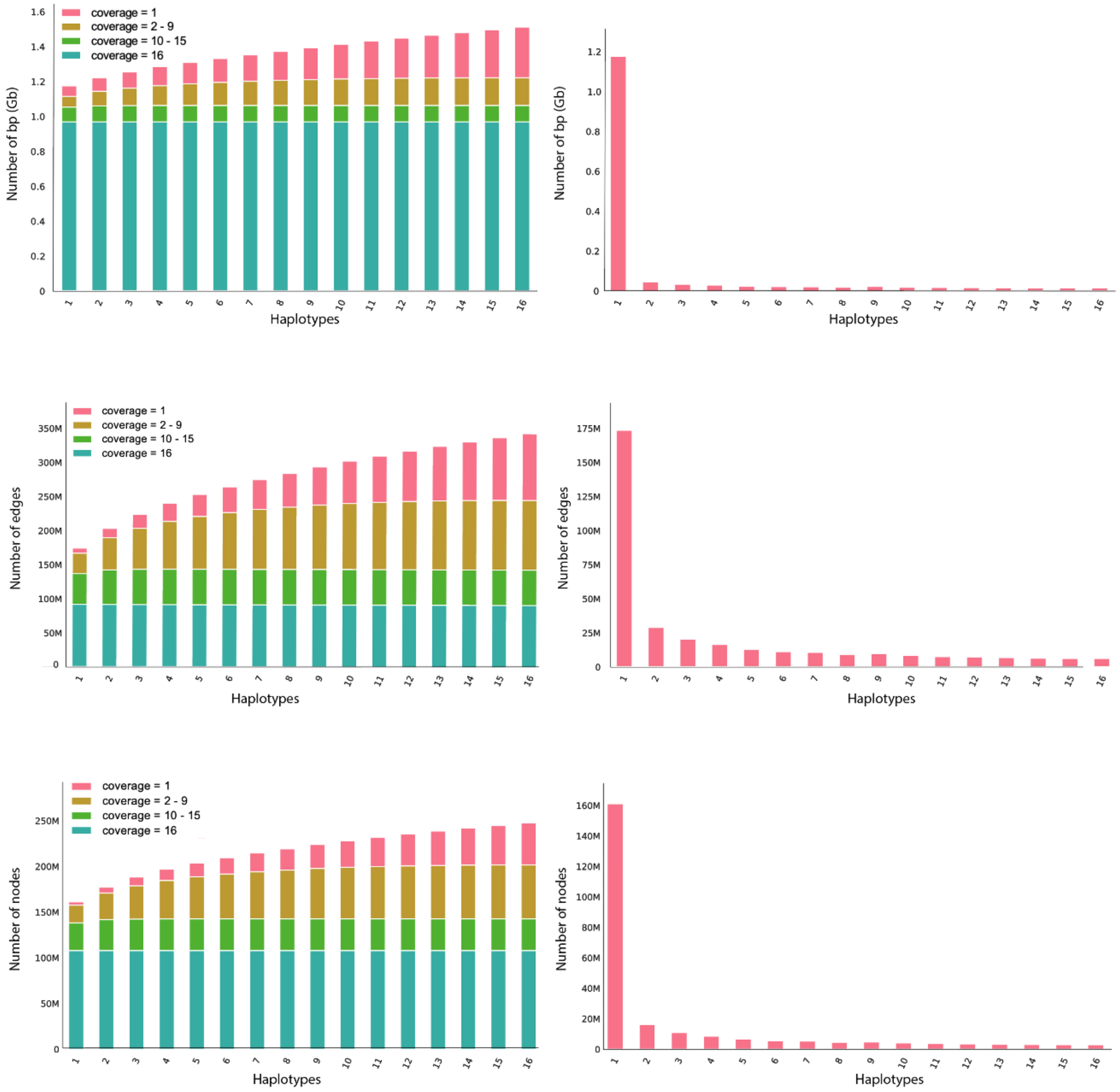

**Figure S3. Pangenome statistics.** Most of the genome is shared among all assemblies. These statistics were computed using Panacus. From top to bottom, shared sequence length (Gb), number of edges, and nodes across varying numbers of haplotypes (1, 2, 10, and 16) as haplotypes are incrementally added. Nodes represent genomic segments, while edges indicate the connections between these segments, defining the structure of the pangenome graph. The right-hand plot in each panel illustrates the incremental contribution of each haplotype to the total sequence length, edges, and nodes as haplotypes are sequentially added.

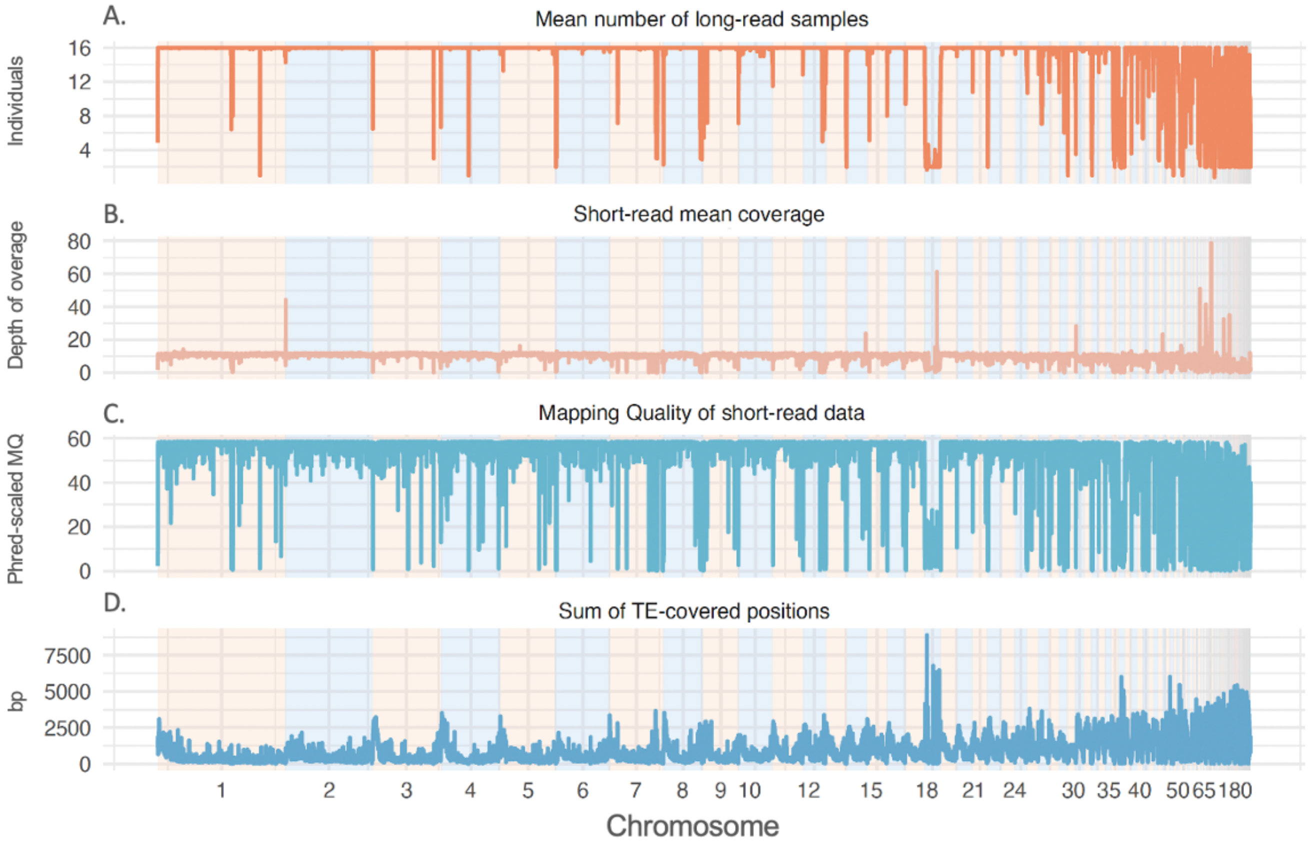

**Figure S4. Genomic overview of long and short-read coverage, short-read mapping quality and TE distribution across chromosomes.** The genomic profile across scaffolds shows: **A)** the mean number of samples supporting the different variants identified in the pangenome through long-read sequencing, **B)** the coverage at each position in the short-read dataset (prior to filtering), **C)** the average Phred-scaled mapping quality of the short-read data aligned to the pangenome across five individuals, where higher Phred scores indicate better mapping quality, and **D)** the total number of positions within a window covered by transposable elements (TEs). Scaffolds are ordered by descending size and represented with alternating background colors, and data are aggregated into 50,000 base pair windows. The x-axis represents the genomic position across all scaffolds, while the y-axis shows the respective values for each track. Regions with lower coverage in long-read sequencing coincide with areas lacking short-read coverage, exhibiting low mapping quality and high TE content, highlighting the challenges of mapping short-read sequencing data in repetitive regions. The short-read data fails to recover 29,598,424 variants from the pangenome.

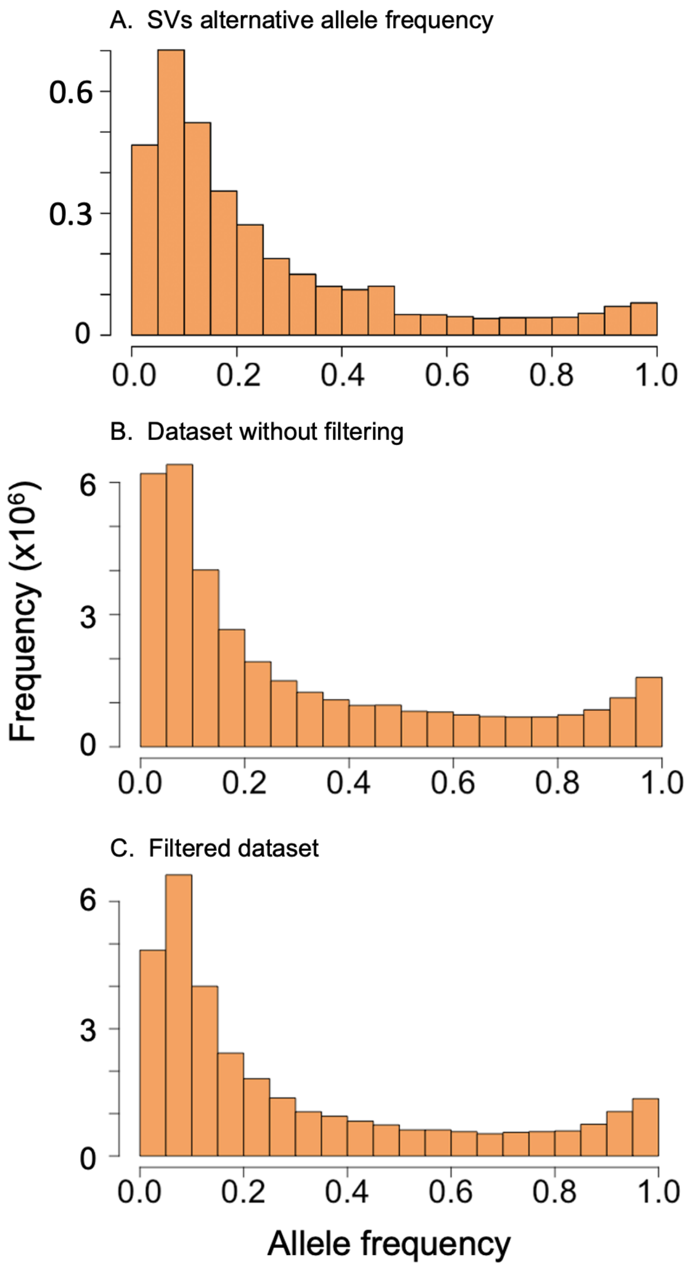

**Figure S5. Allele frequency distribution of variants** for **A)** Alternative allele frequency distribution of SVs identified from short-read data mapped to the pangenome **B)** the dataset including both SNPs and SVs without filtering and **C)** the dataset including both SNPs and SVs filtered with VCFtools, which was refined based on missing data (0.8), minimum and maximum mean depth (4 and 50, respectively), and a non-reference allele count of at least 4 (--non-ref-ac-any 4). Alternative allele frequency distribution of SVs shows the same pattern as the SNPs. In the unfiltered dataset, the first two bins have nearly the same frequency. After filtering, the bin representing the smallest allele frequencies (0–0.05) is reduced because low-frequency variants are removed during filtering. For instance, the non-reference allele count filter eliminates variants observed in very few individuals, reducing the number of rare alleles. The filtered distribution closely follows the pattern observed in the SNP and SV datasets separately.

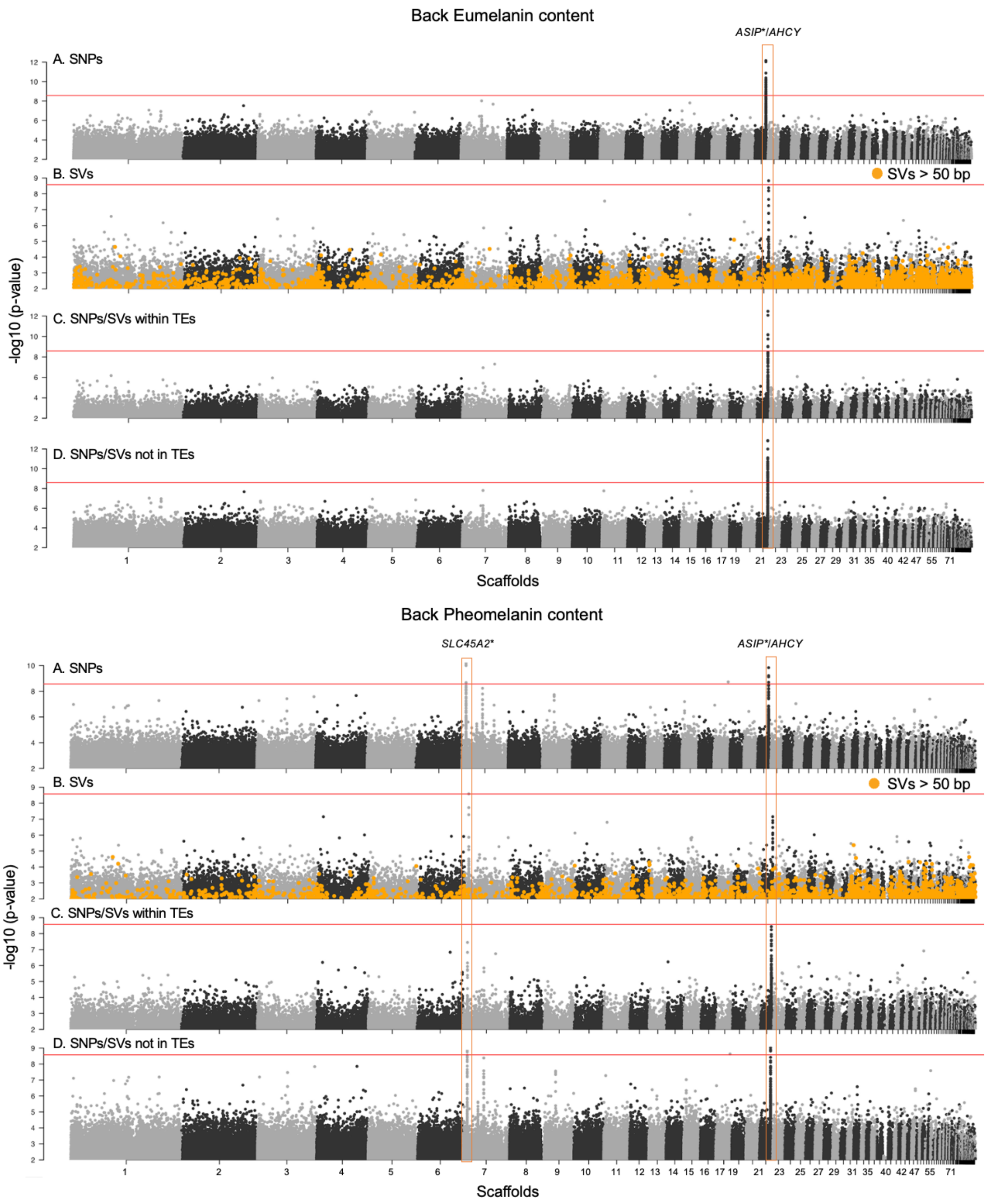

**Figure S6. Genome wide association study for the eumelanin and pheomelanin content in the back plumage patch.** The analysis includes four datasets, displayed from top to bottom: A) SNPs, B) Structural variants including short (indel) and long variants (SVs > 50 bp, highlighted in orange), C) Variants (SNPs and SVs) within annotated transposable elements (TEs), and D)

Variants (SNPs and SVs) outside TEs. The y axis represents the -log10(p-value) obtained in the GWAS analysis. The red line represents the significance threshold after Bonferroni correction, corresponding to a p-value of $2.65 * 10^{-9}$, corrected for all comparisons, including patches/pigment combinations and all variants. Scaffolds are ordered by decreasing size and represented in alternating black/gray. Peaks are highlighted with orange rectangles, and known genes are labeled above the peaks. The genes marked with an asterisk (*) belong to the melanogenesis pathway. The *ASIP* and *AHCY* genes are consistently associated to changes in both eumelanin and pheomelanin pigmentation. For eumelanin, these genes are recovered across all datasets, although in the SV dataset variants larger than 50 bp are not outliers. For pheomelanin, the *ASIP*/*AHCY* peak is detected only by the SNP dataset and the dataset that includes variants outside transposable elements (TEs). Additionally, the peak including the *SLC45A2* gene, which is associated with pheomelanin content, is recovered in all datasets except the one restricted to variants within TEs. Again, we did not find large SVs involved in this peak.

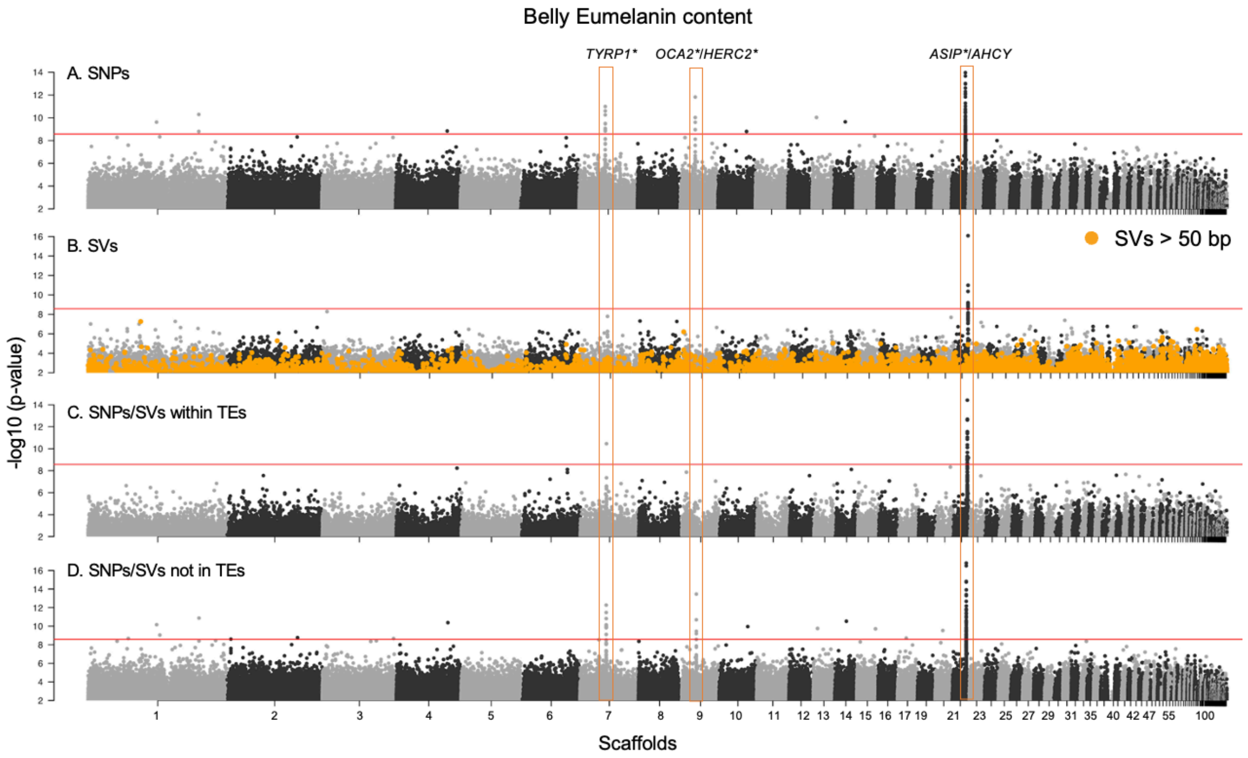

**Figure S7. Genome wide association study for the eumelanin content in the belly plumage patch.** Details as in Figure S6. The peaks associated with eumelanin content include *TRYP1* on scaffold 7, which is recovered by all datasets except the one focusing on structural variants (larger and smaller than 50 bp), and *OCA2*/*HERC2* on scaffold 10, which is exclusively recovered by the SNP dataset and variants outside transposable elements (TEs). Additionally, the peak containing the *ASIP* and *AHCY* genes on scaffold 21 is consistently recovered across all datasets, although large variants were not outliers in the SV dataset. The corresponding plots for the pheomelanin content of the belly plumage patch can be found in the main manuscript (Figure 3).

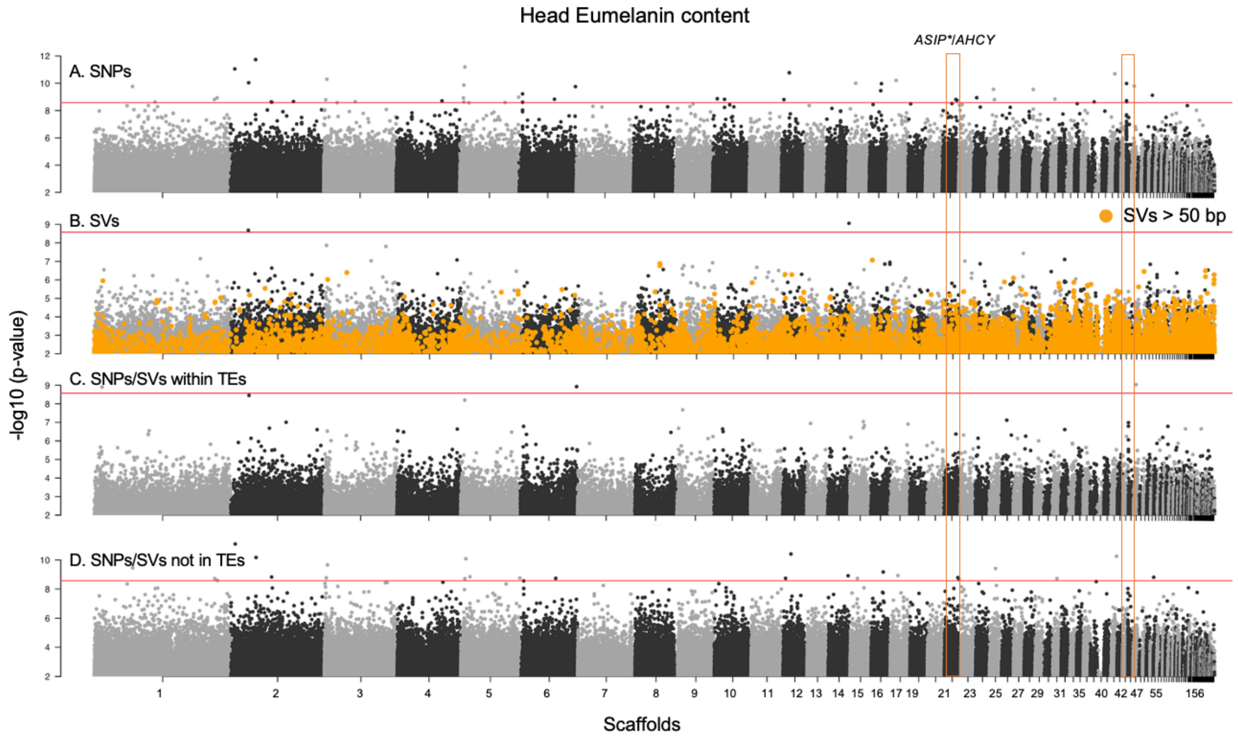

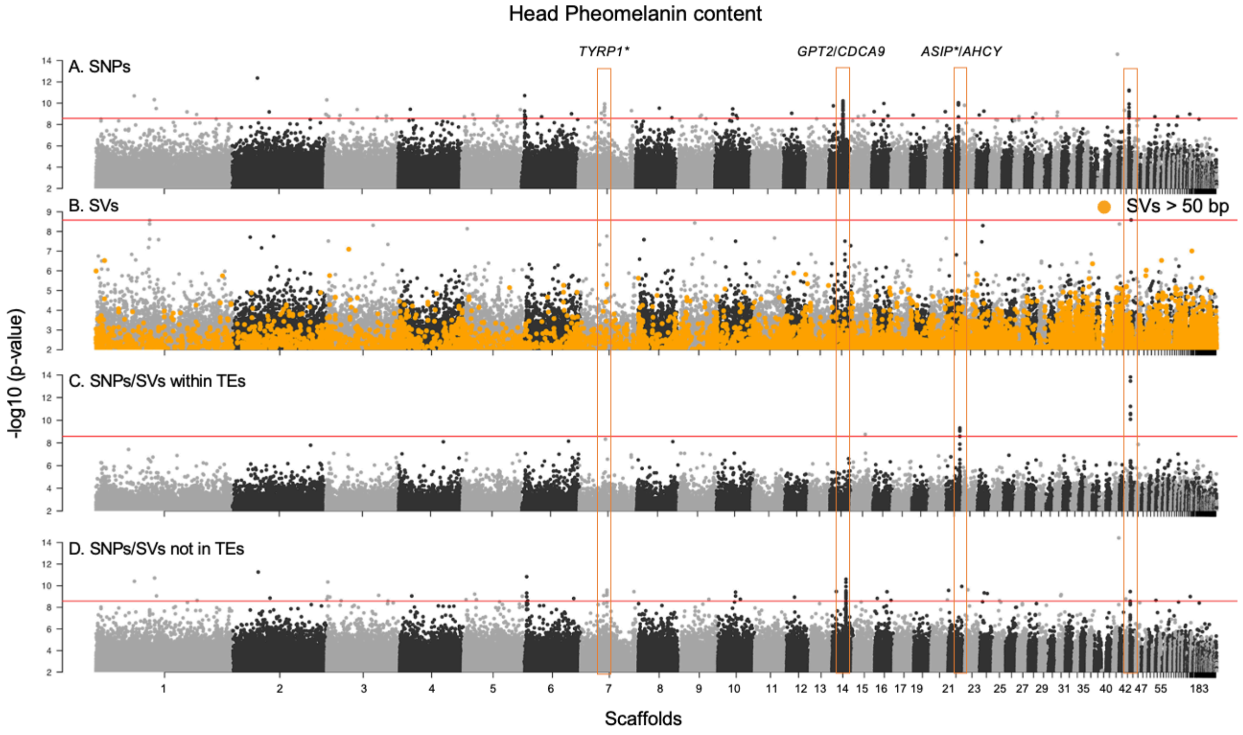

**Figure S8. Genome wide association study for the eumelanin and pheomelanin content in the head plumage patch.** Details as in Figure S6. Overall, the comparisons for the head plumage patch are noisier than those for other regions of the body. The peaks associated with eumelanin content include the *ASIP* and *AHCY* genes on scaffold 21 and an additional peak on scaffold 41 with no annotated genes. Both peaks are recovered exclusively by the SNP dataset. For pheomelanin content, we identified four peaks. These include again the peak containing the *ASIP* and *AHCY* genes on scaffold 21 and the peak on scaffold 41 that are recovered across all datasets except the one focusing on structural variants. Additionally, the *TRYP1* gene on scaffold 7 and the *GPT2* and *CDCA9* genes on scaffold 14, are both recovered only by the SNP dataset and variants outside TEs.

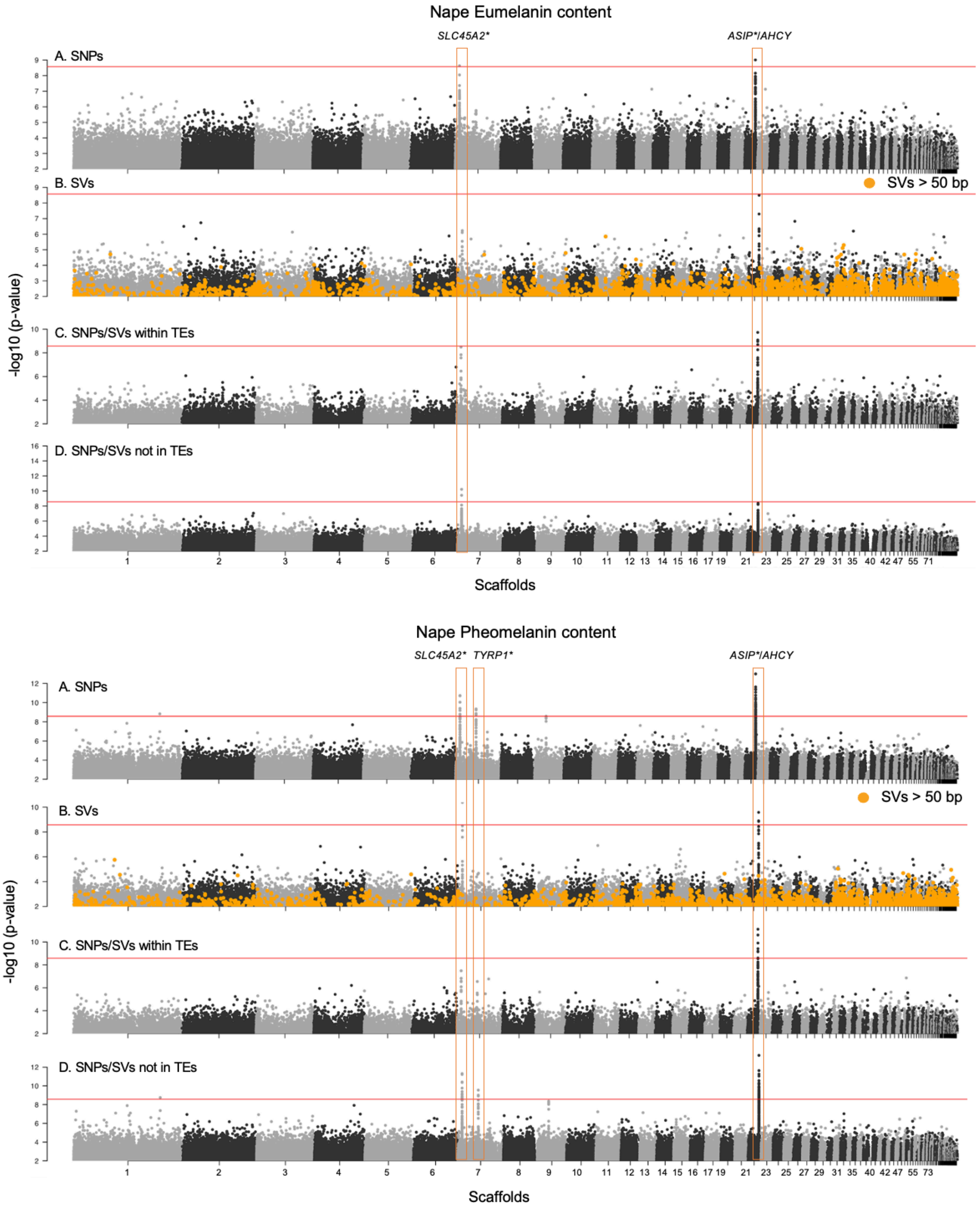

**Figure S9. Genome wide association study for the eumelanin and pheomelanin content in the nape plumage patch.** Details as in Figure S6. The peaks associated with eumelanin content include the *SLC45A2* gene on scaffold 7 and the *ASIP* and *AHCY* genes on scaffold 21. Both peaks are recovered by the SNP dataset; the one on scaffold 7 is also detected by the dataset of variants outside transposable elements (TEs), while the one on scaffold 21 is also identified by the dataset of variants within TEs. For pheomelanin content, in addition to these two peaks—which are recovered by all datasets except for the variants within TEs for the *SLC45A2* peak—another peak is identified on scaffold 7 containing the *TYRP1* gene. This peak is recovered by the SNP dataset and the dataset of variants outside TEs. In all cases in which an outlier peak was found with the SV dataset, the variants involved were smaller than 50 bp.

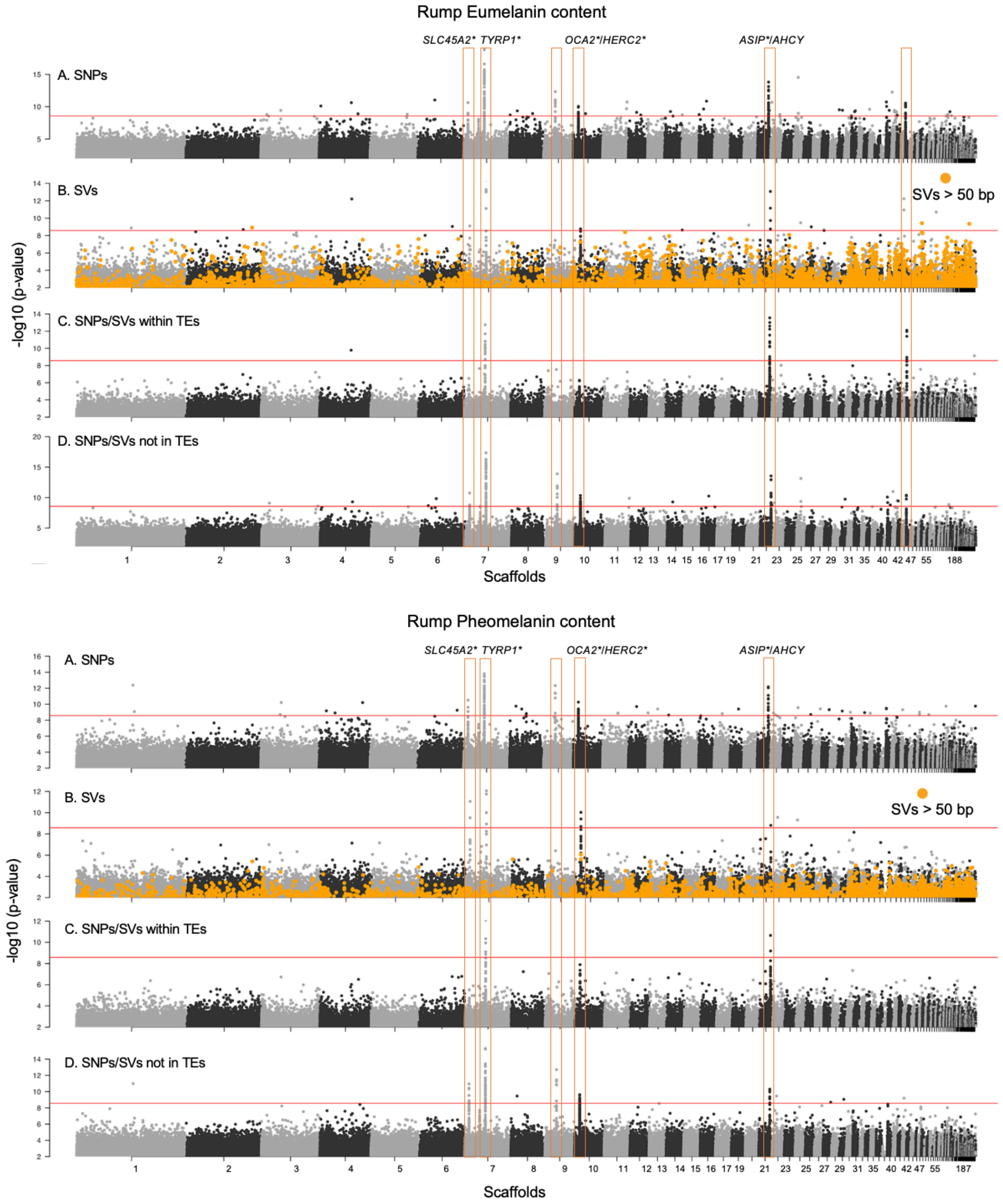

**Figure S10. Genome wide association study for the eumelanin and pheomelanin content in the rump plumage patch.** Details as in Figure S6. We detected six peaks associated with eumelanin content, including two on scaffold 7: one containing the *SLC45A2* gene and the other containing *TYRP1*. Additionally, there is one peak on scaffold 9 and another on scaffold 41, both lacking annotated genes. A peak on scaffold 10 contains the *HERC2* gene, and a peak on scaffold

21 contains the *ASIP* and *AHCY* genes. The peaks containing *TYRP1*, *ASIP*/*AHCY*, and the one on scaffold 41 are detected by all datasets. The *SLC45A2* and *HERC2* peaks are detected by all datasets except the one containing variants within TEs, while the peak on scaffold 9 is detected only by the SNP and variants outside TEs datasets. Notably, three long SVs were associated with eumelanin content; however, these variants are not located within the peaks, do not overlap any gene, and show a very low $F_{ST}$ for all comparisons. The only variant with slightly higher differentiation is an 83 bp insertion on scaffold 2 (position 80,341,265), where $F_{ST}$ values range from 0.280 to 0.474 in several comparisons involving *S. hypoxantha*, *S. melanogaster*, *S. S. ruficollis*, *S. cinnamomea*, *S. hypochroma*, *and S. iberaensis*. For pheomelanin content, we detected the same peaks except for the one on scaffold 41. The detected peaks were identified by the same datasets as in the eumelanin GWAS for this plumage patch. In all cases in which an outlier peak was found with the SV dataset, the variants involved were smaller than 50 bp

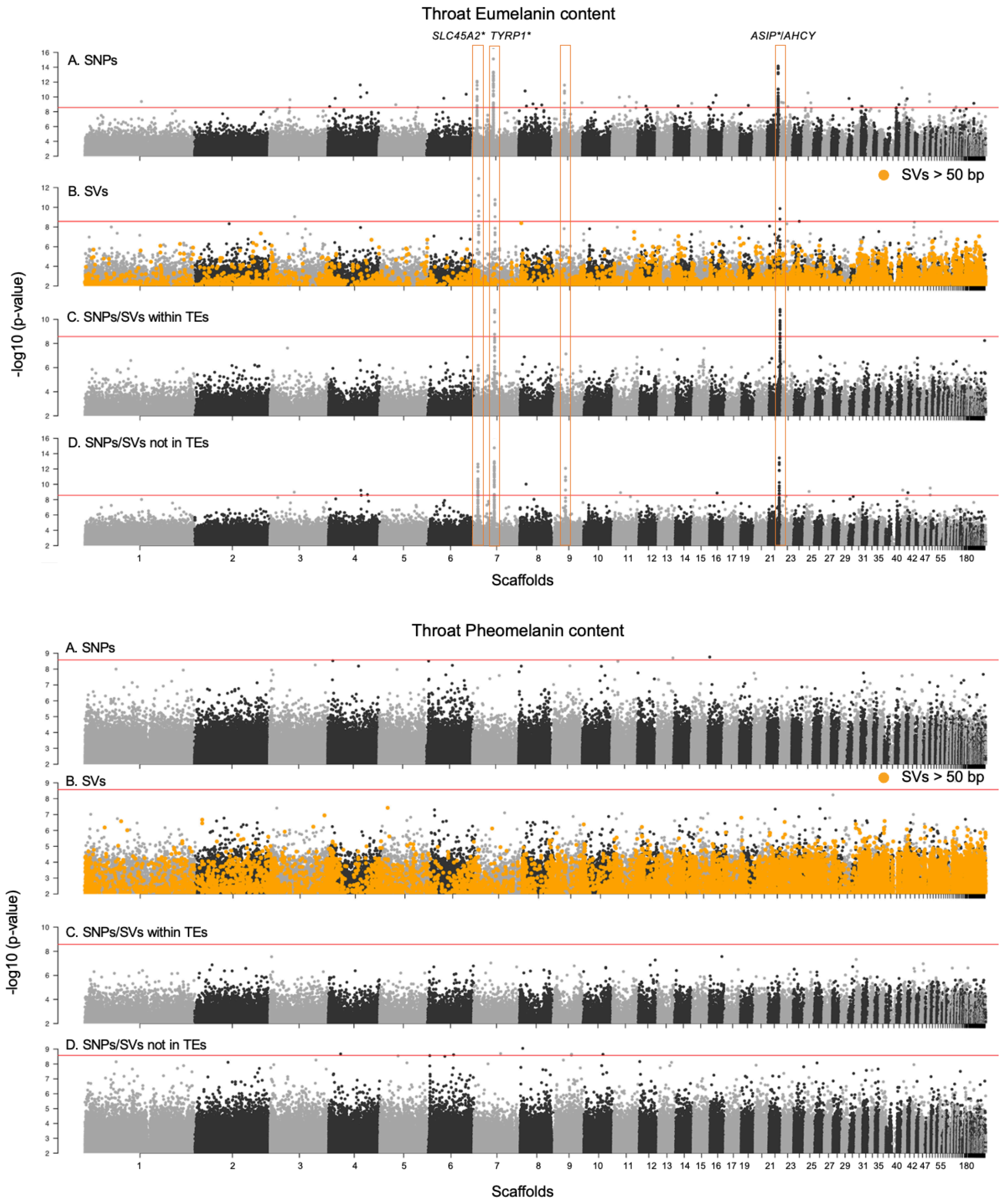

**Figure S11. Genome wide association study for the eumelanin and pheomelanin content in the throat plumage patch.** Details as in Figure S6. We detected four peaks associated with eumelanin content, including two on scaffold 7: one containing the *SLC45A2* gene and the other containing *TYRP1*. Additionally, there is one peak on scaffold 9 with no annotated genes and a peak on scaffold 21 that contains the *ASIP* and *AHCY* genes. The peaks containing *TYRP1* and

*ASIP*/*AHCY* are detected by all datasets. The *SLC45A2* peak is detected by all datasets except the one containing variants within TEs, while the peak on scaffold 9 is detected only by the SNP and variants outside TEs datasets. For pheomelanin content, no peaks were detected. In all cases in which an outlier peak was found with the SV dataset, the variants involved were smaller than 50 bp.

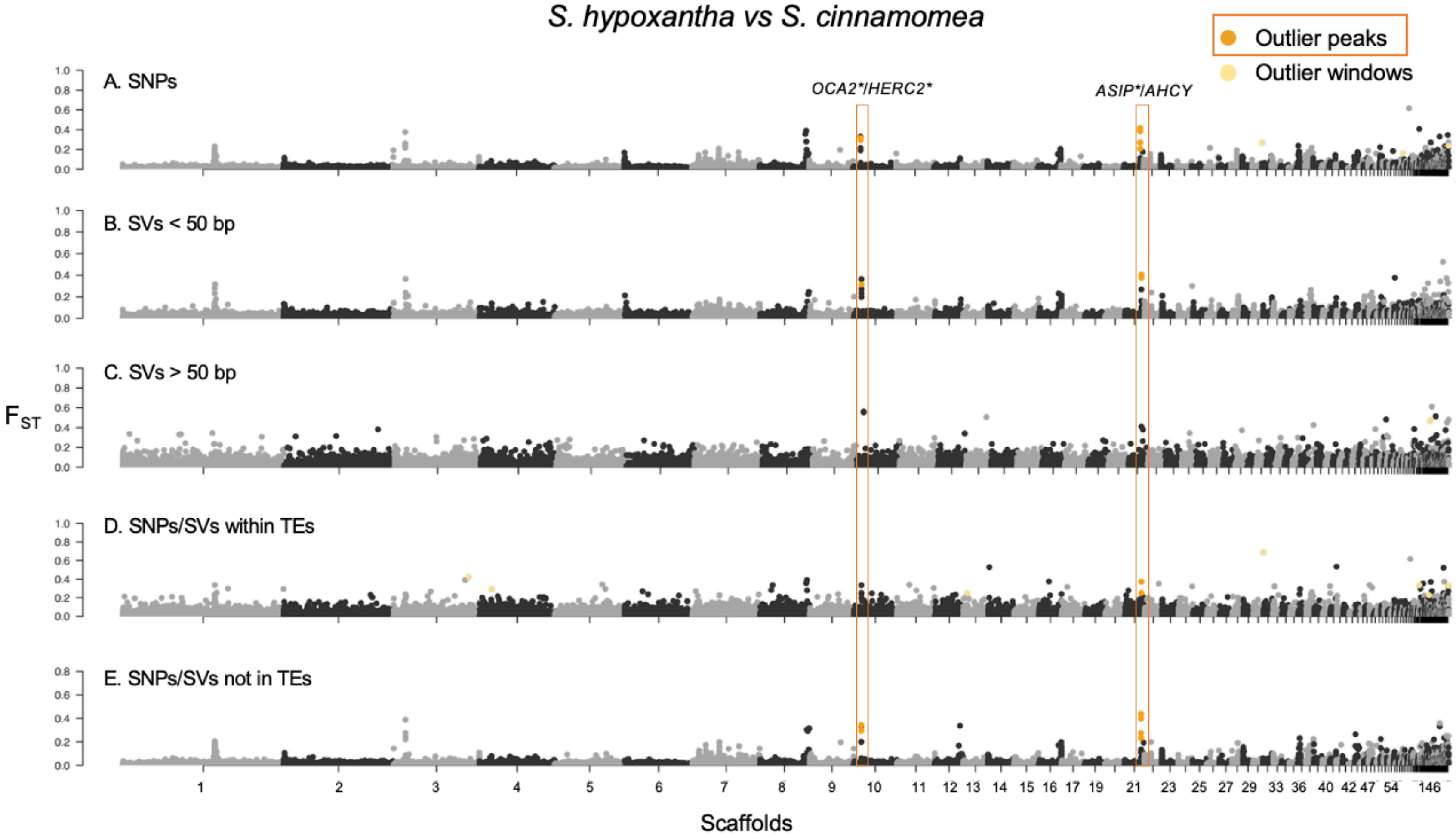

**Figure S12. $F_{ST}$ scan in 10 kb windows for the comparison between *S. hypoxantha* and *S. cinnamomea*.** The analysis includes five datasets, displayed from top to bottom: A) SNPs, B) Indels: Structural variants <50bp, C) Structural variants >50bp, D) Variants (SNPs and SVs) within transposable elements (TEs), and E) Variants (SNPs and SVs) outside TEs. Scaffolds are ordered by decreasing size and represented in alternating colors. Peaks are marked with an orange rectangle, and the known genes are labeled on top of the peaks. The gene marked with an asterisk (*) belong to the melanogenesis pathway. Outlier windows outside the peaks are marked in light orange. In this comparison, we detected two peaks: one including *OCA2*/*HERC2* on scaffold 10 and the other containing *ASIP* and *AHCY* on scaffold 21. Both peaks are detected using the SNP, indel, and variants outside TEs datasets, while the *ASIP*/*AHCY* peak is also detected by the variants within TEs dataset. Outlier windows outside the peaks are marked in light orange (see methods for details).

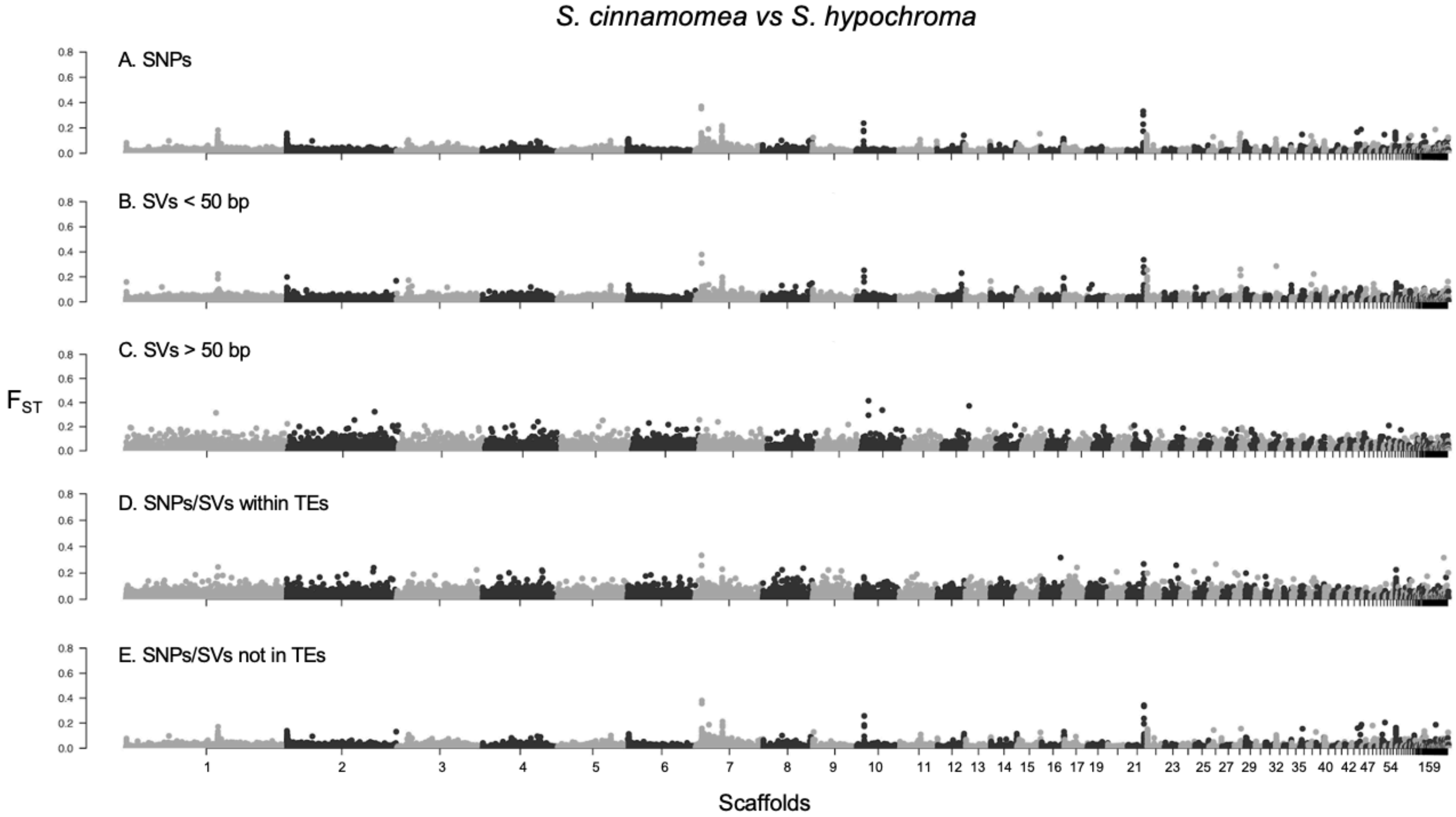

**Figure S13. $F_{ST}$ scan in 10 kb windows for the comparison between *S. cinnamomea* and *S. hypochroma*.** Details as in Figure S12. There are no outliers in this pairwise comparison under the set of criteria described in the methods section.

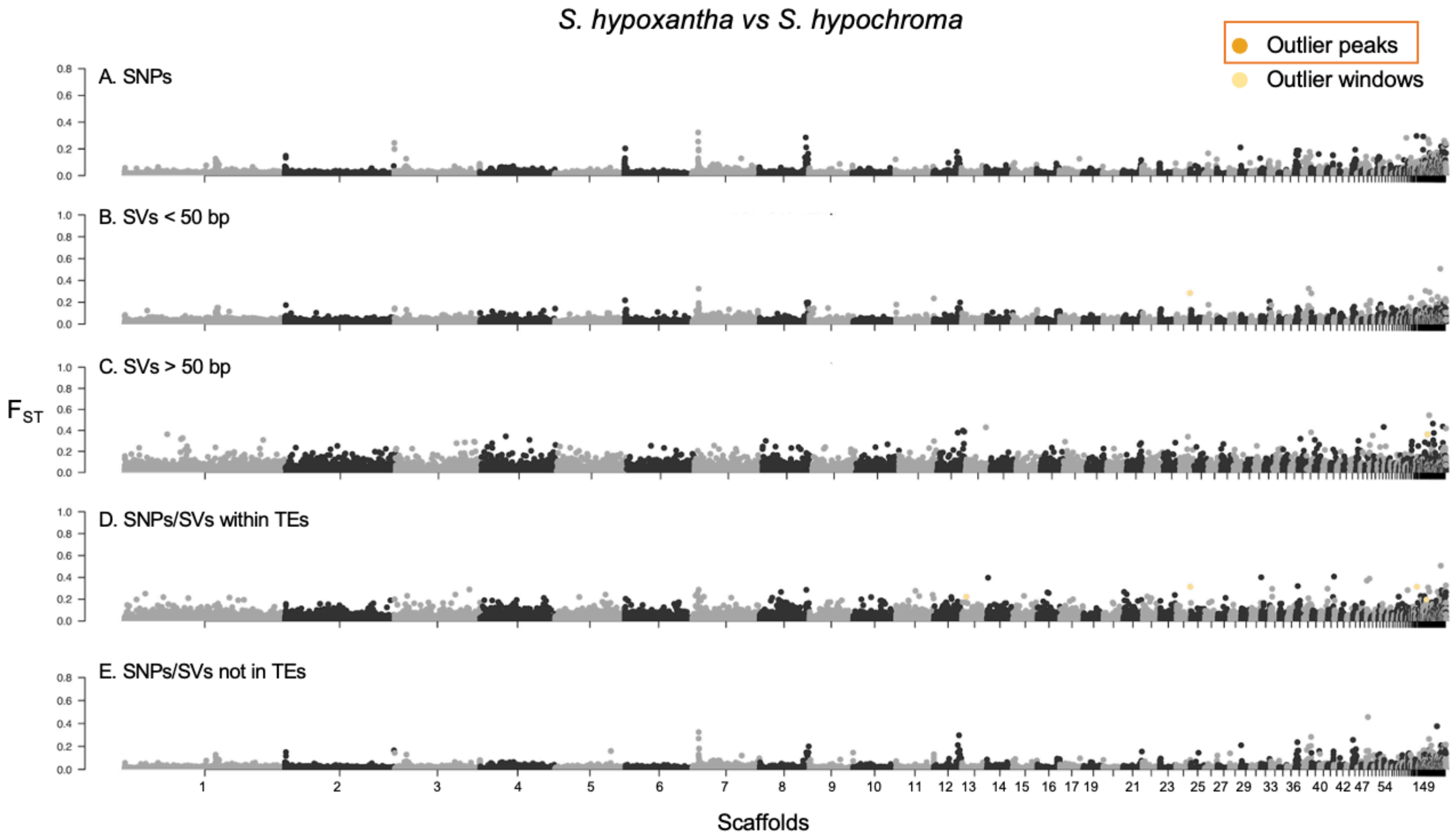

**Figure S14. $F_{ST}$ scan in 10 kb windows for the comparison between *S. hypoxantha* and *S. hypochroma*.** Details as in Figure S12. There are no outlier peaks in this pairwise comparison.

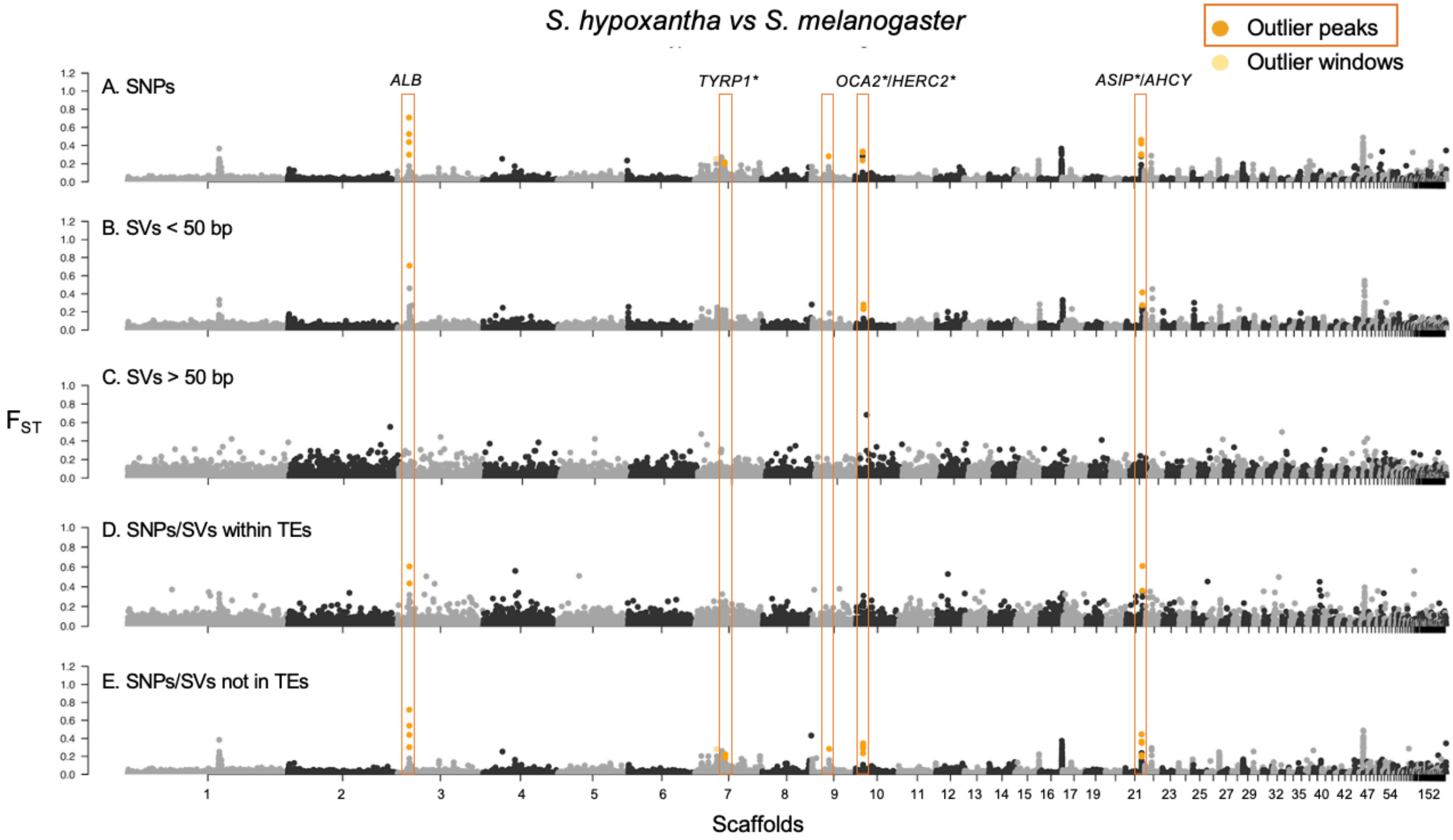

**Figure S15. $F_{ST}$ scan in 10 kb windows for the comparison between *S. hypoxantha* and *S. melanogaster*.** Details as in Figure S12. There are five peaks in this comparison, including one containing the *ALB* gene on scaffold 3 and one containing *ASIP* and *AHCY* on scaffold 21, which are detected by all datasets except the long SVs. Additionally, there is one peak on scaffold 7 containing *TYRP1* and another on scaffold 9 with no annotated genes, both detected only by the SNPs and variants outside TEs datasets. A fifth peak on scaffold 10, containing *OCA2*/*HERC2*, is detected by the same datasets as the previous peaks, as well as the short SVs.

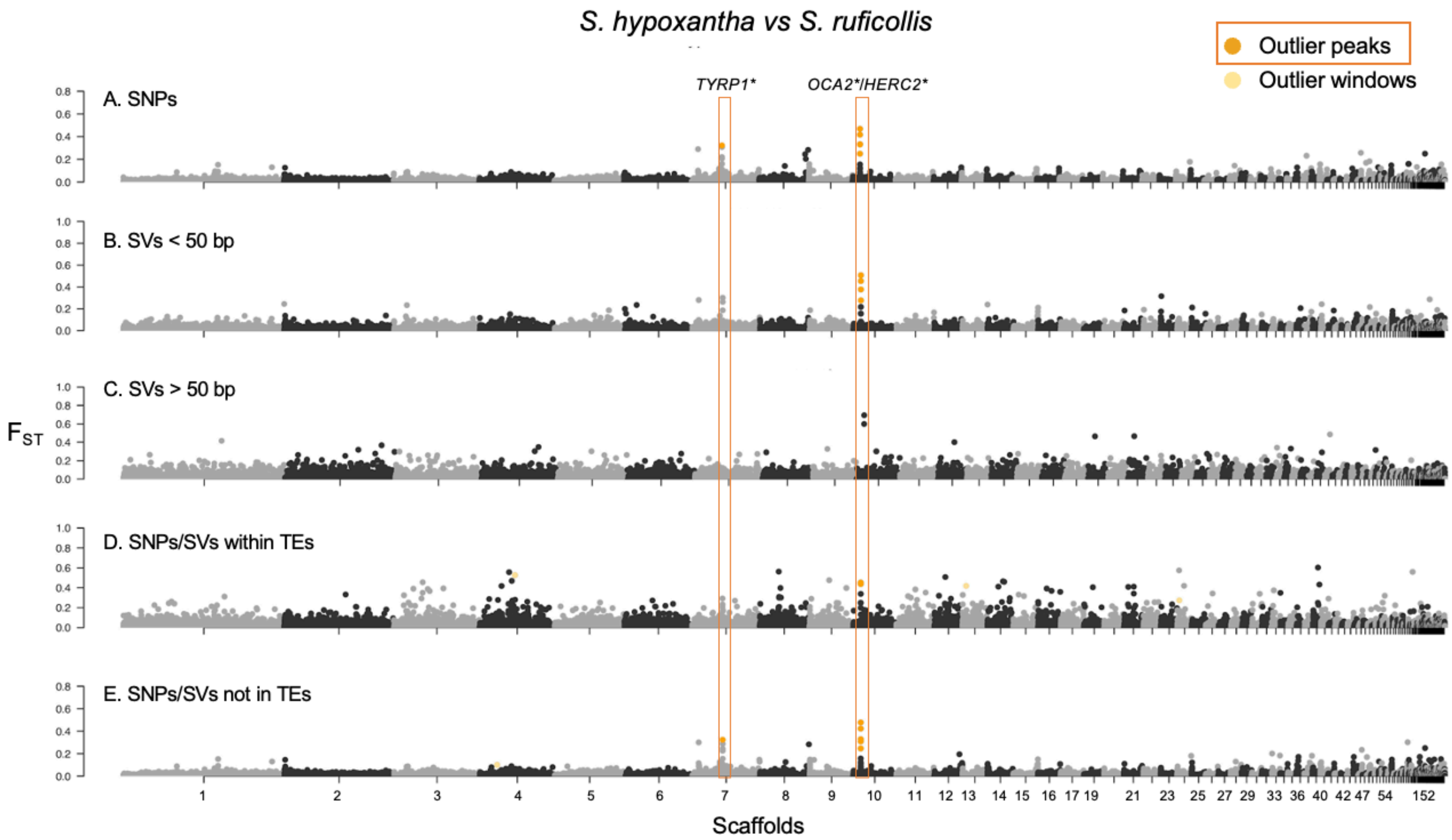

**Figure S16. $F_{ST}$ scan in 10 kb windows for the comparison between *S. hypoxantha* and *S. ruficollis*.** Details as in Figure S12. There are two peaks in this comparison: one containing *TYRP1* on scaffold 7, detected only by the SNP and variants outside TEs datasets, and another containing *OCA2*/*HERC2* on scaffold 10, detected by all datasets except the long SVs.

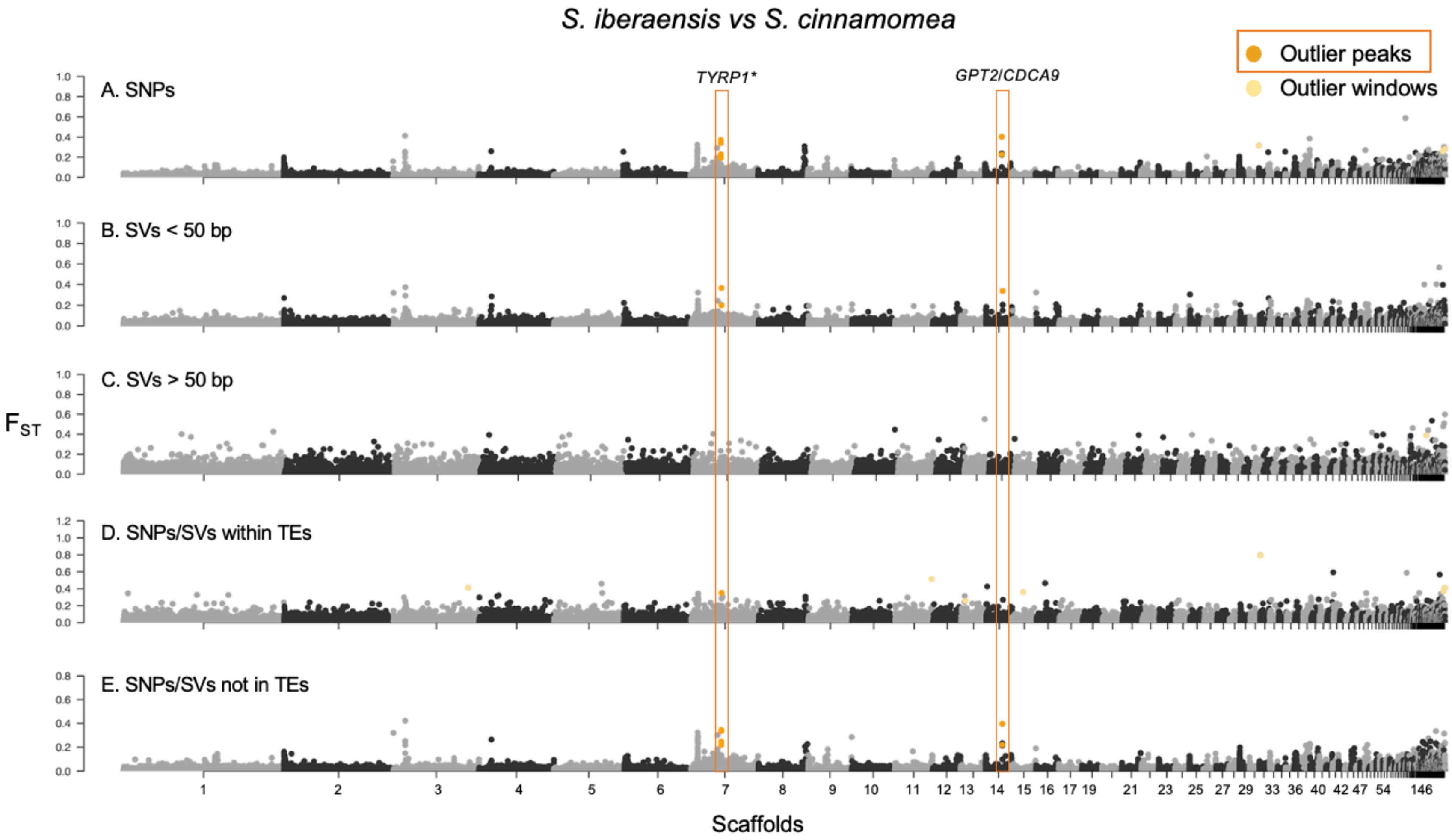

**Figure S17. $F_{ST}$ scan in 10 kb windows for the comparison between *S. iberaensis* and *S. cinnamomea*.** Details as in Figure S12. There are two peaks in this comparison: one containing *TYRP1* on scaffold 7, detected by all datasets except the long SVs, and another containing *GPT2* and *CDCA9* on scaffold 14, detected by all datasets except the long SVs and the variants within TEs.

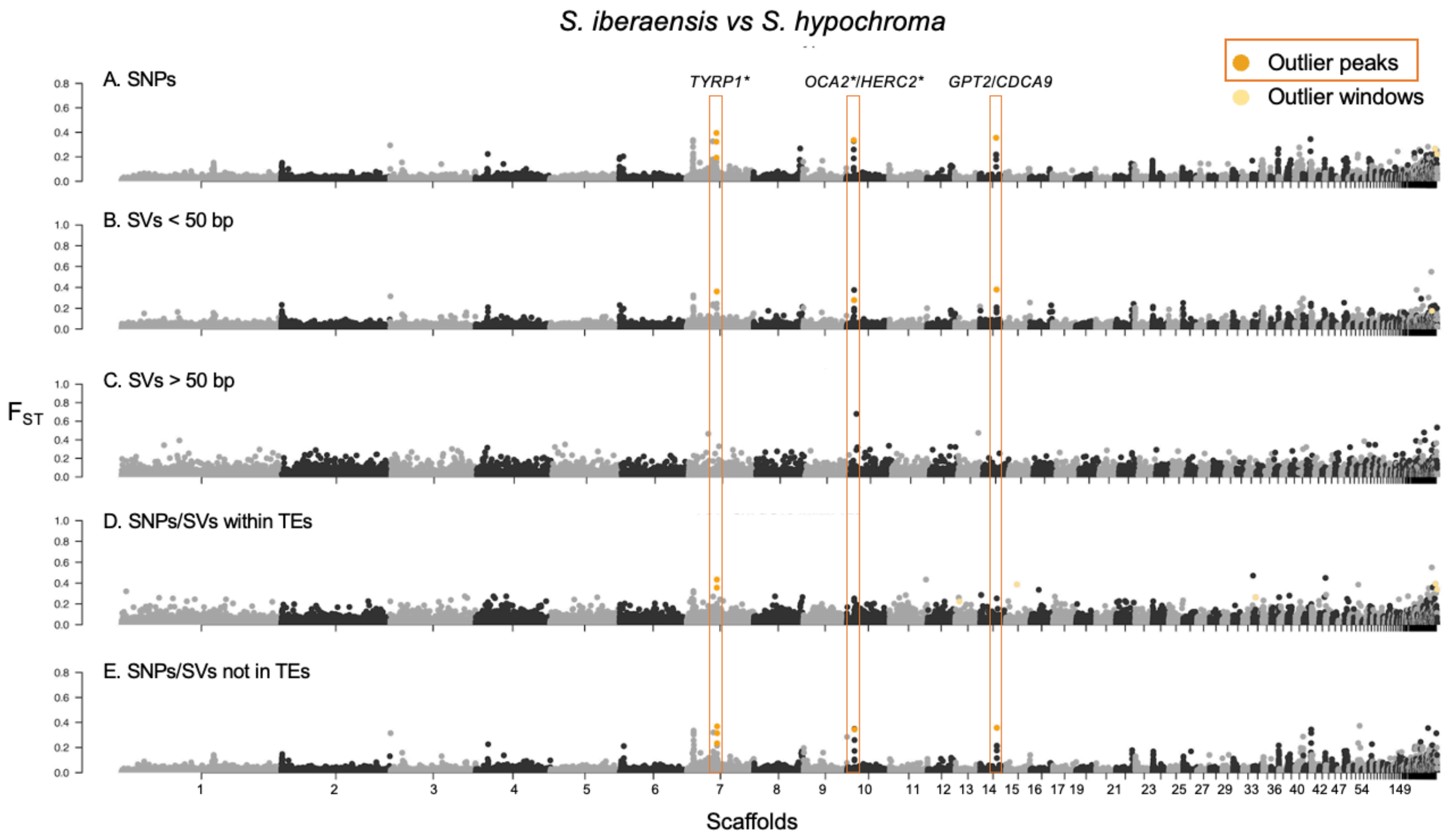

**Figure S18. $F_{ST}$ scan in 10 kb windows for the comparison between *S. iberaensis* and *S. hypochroma*.** Details as in Figure S12. There are three peaks in this comparison: one containing *TYRP1* on scaffold 7, detected by all datasets except the long SVs; and two others, one containing *OCA2*/*HERC2* on scaffold 10 and one containing *GPT2* and *CDCA9* on scaffold 14, both detected by all datasets except the long SVs and the variants within TEs.

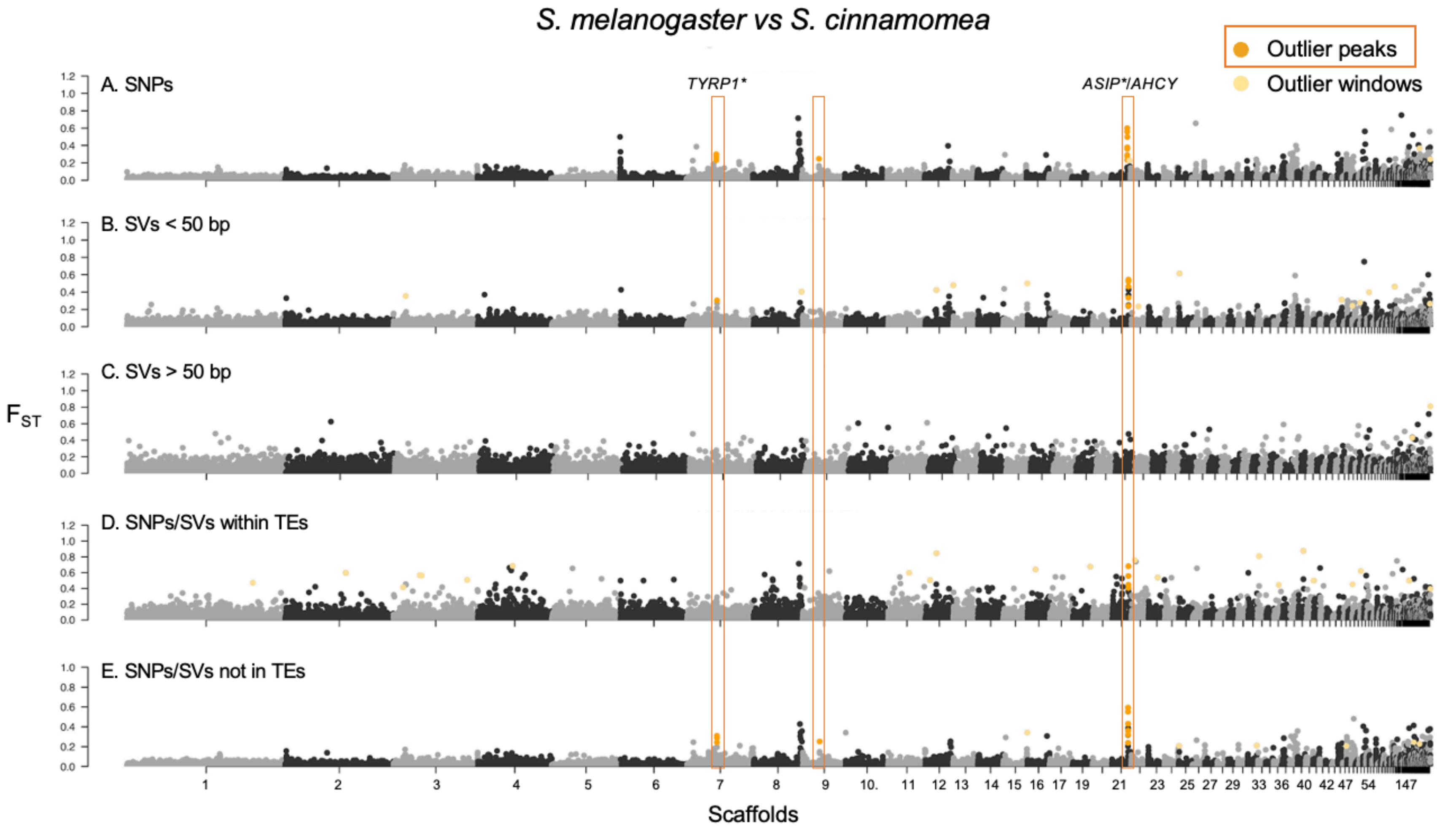

**Figure S19. $F_{ST}$ scan in 10 kb windows for the comparison between *S. melanogaster* and *S. cinnamomea*.** Details as in Figure S12. There are three peaks in this comparison. One containing *ASIP* and *AHCY* on scaffold 21, detected by all datasets except the long SVs. A second peak containing *TYRP1* on scaffold 7, detected by all datasets except the long SVs and the variants within TEs. The third peak, with no annotated genes, on scaffold 9, was detected only by the SNPs and variants outside TEs datasets.

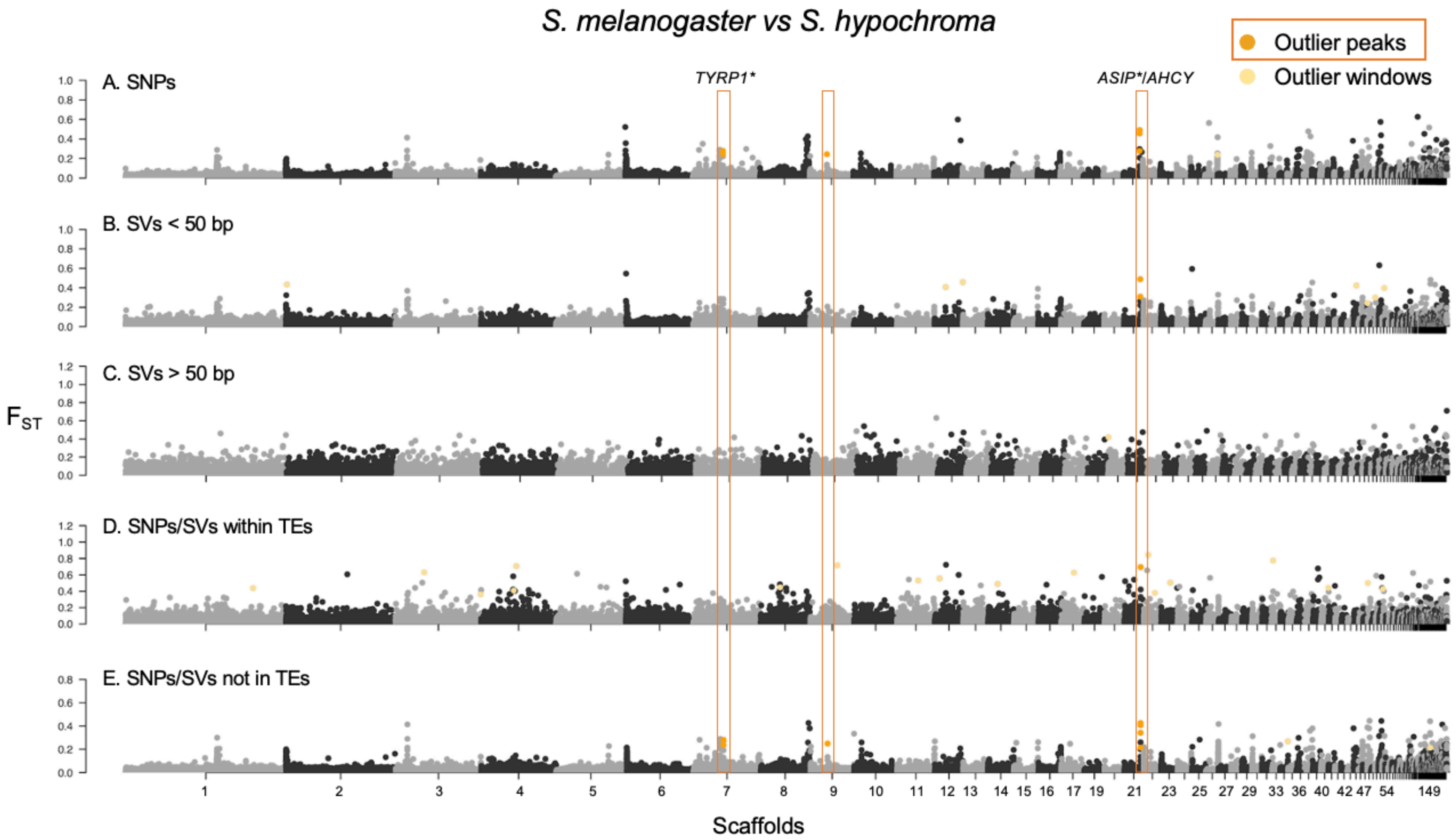

**Figure S20. $F_{ST}$ scan in 10 kb windows for the comparison between *S. melanogaster* and *S. hypochroma*.** Details as in Figure S12. There are three peaks in this comparison. One peak containing *ASIP* and *AHCY* on scaffold 21, detected by all datasets except the long SVs. The two remaining peaks, one containing *TYRP1* on scaffold 7 and one with no annotated genes on scaffold 9, were both detected only by the SNPs and variants outside TEs datasets.

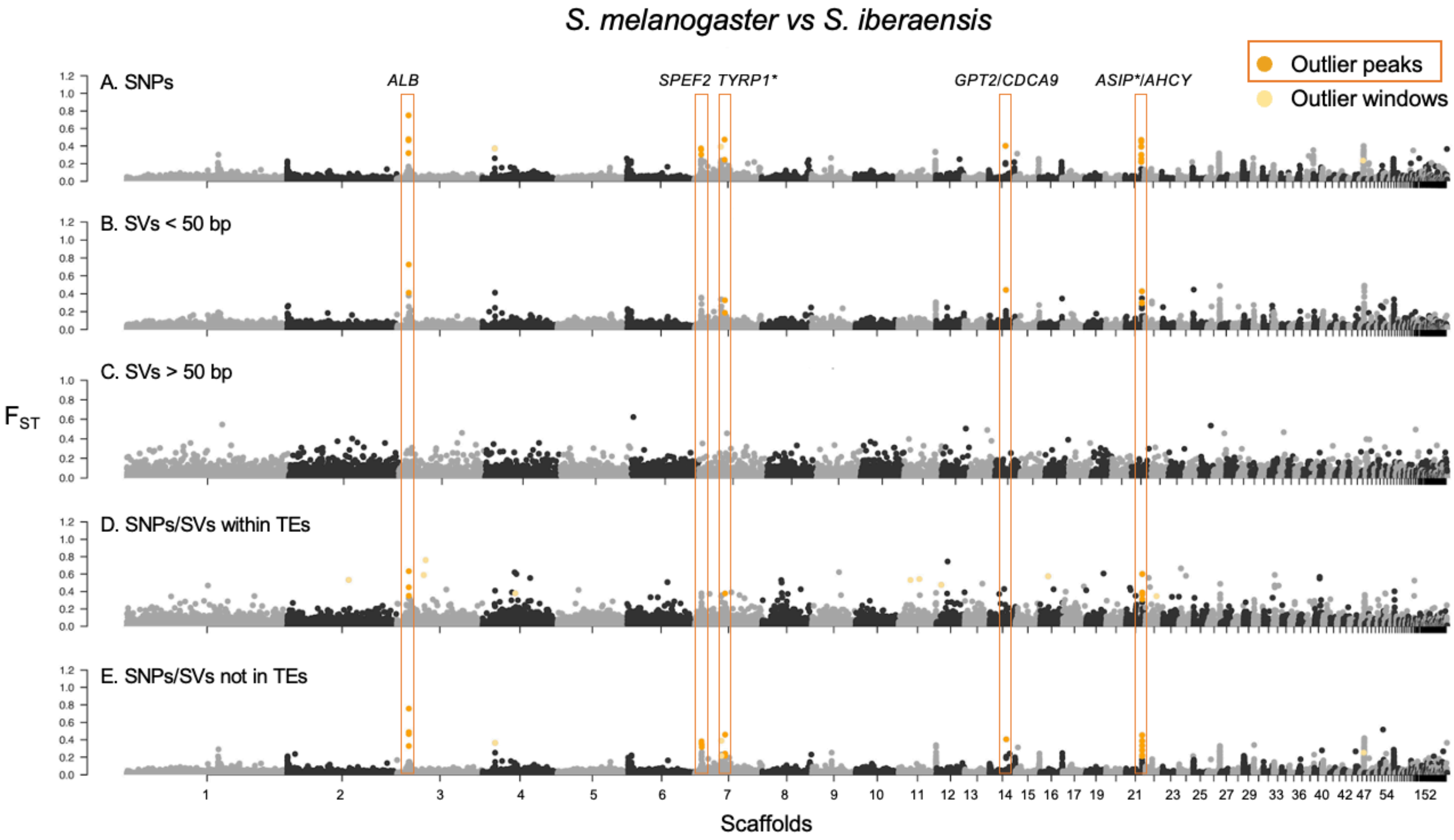

**Figure S21. $F_{ST}$ scan in 10 kb windows for the comparison between *S. melanogaster* and *S. iberaensis*.** Details as in Figure S12. There are five peaks in this comparison, including the genes *ALB* on scaffold 3, *TYRP1* on scaffold 7, and *ASIP* and *AHCY* on scaffold 21, which are detected by all datasets except the long SVs. Additionally, there is one peak on scaffold 7 containing *SPEF2*, detected only by the SNPs and variants outside TEs datasets, and another on scaffold 14 containing the genes *GPT2* and *CDCA9*, detected by the same datasets as the previous peak plus the one including the short SVs.

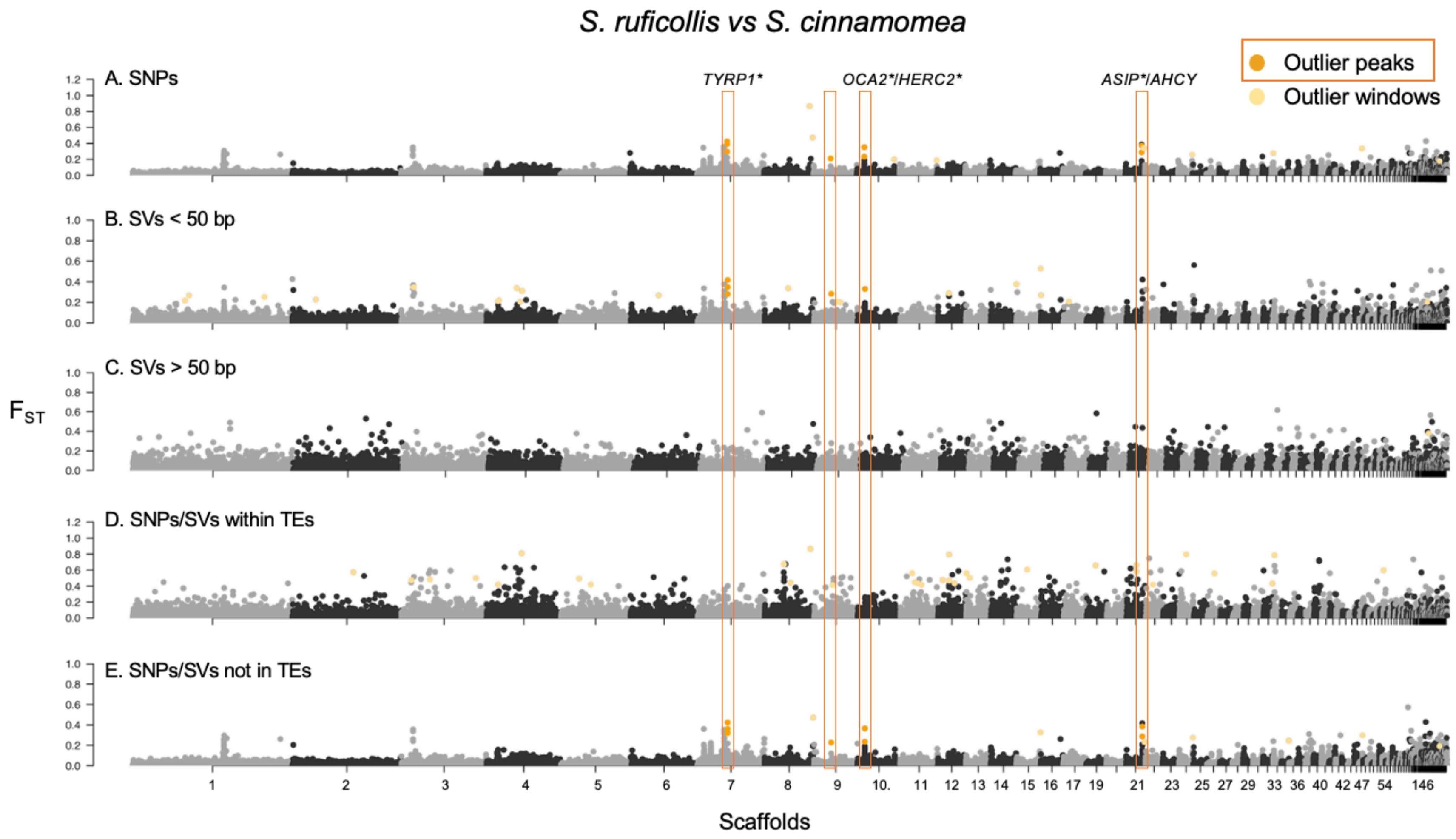

**Figure S22. $F_{ST}$ scan in 10 kb windows for the comparison between *S. ruficollis* and S. *cinnamomea*.** Details as in Figure S12. There are four peaks in this comparison: one containing the gene *TYRP1* on scaffold 7, one on scaffold 9 with no annotated genes, and one containing *OCA2*/*HERC2* on scaffold 10, which are detected by all datasets except the long SVs and the variants within TEs.

Additionally, there is one peak on scaffold 21 containing *ASIP* and *AHCY*, which is detected only by the SNPs and variants outside TEs datasets.

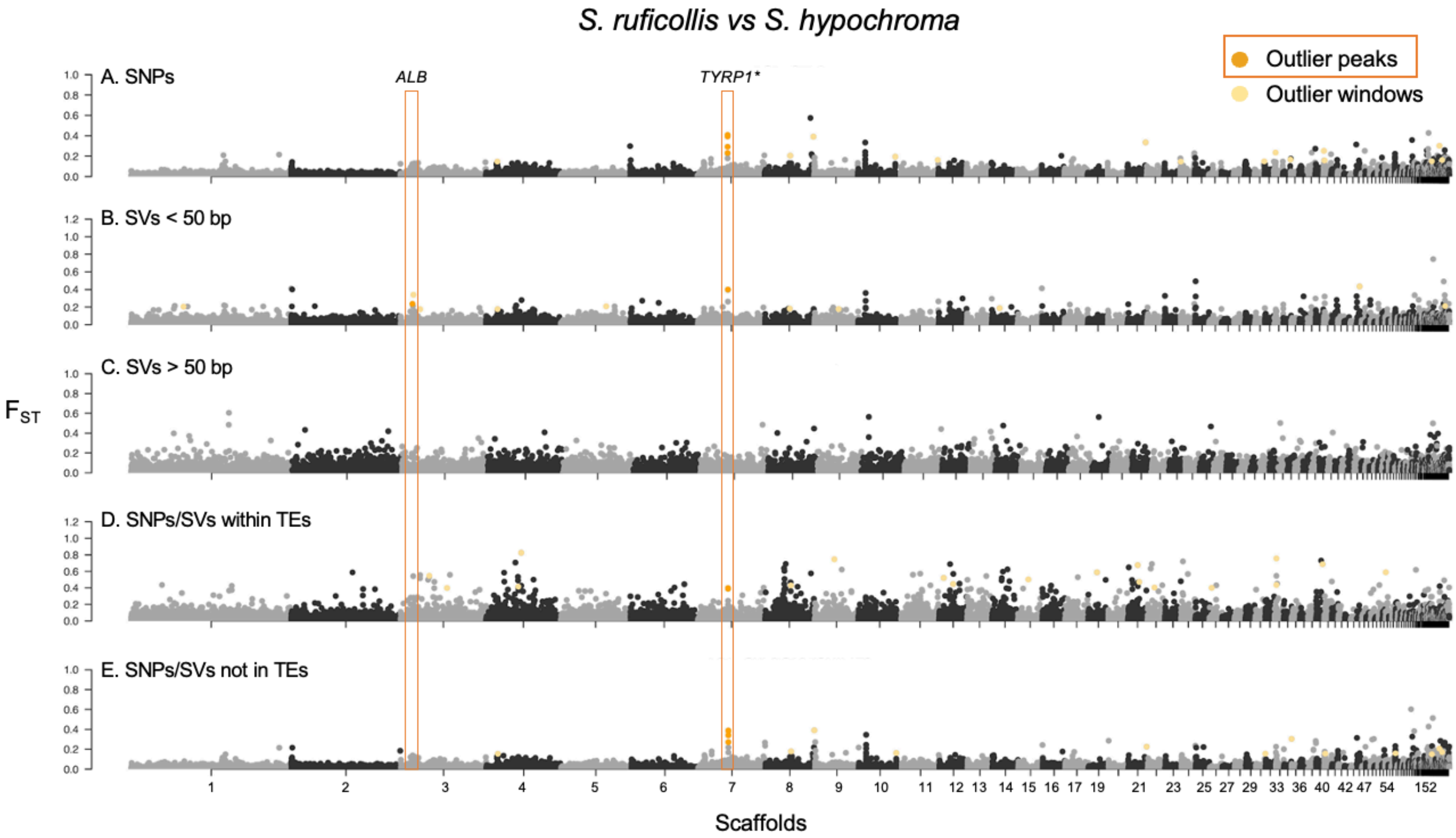

**Figure S23. $F_{ST}$ scan in 10 kb windows for the comparison between *S. ruficollis* and *S. hypochroma*.** Details as in Figure S12. There are two peaks in this comparison: one containing the gene *TYRP1* on scaffold 7, detected by all datasets except the long SVs, and another containing *ALB* on scaffold 3, detected only by the short SVs dataset.

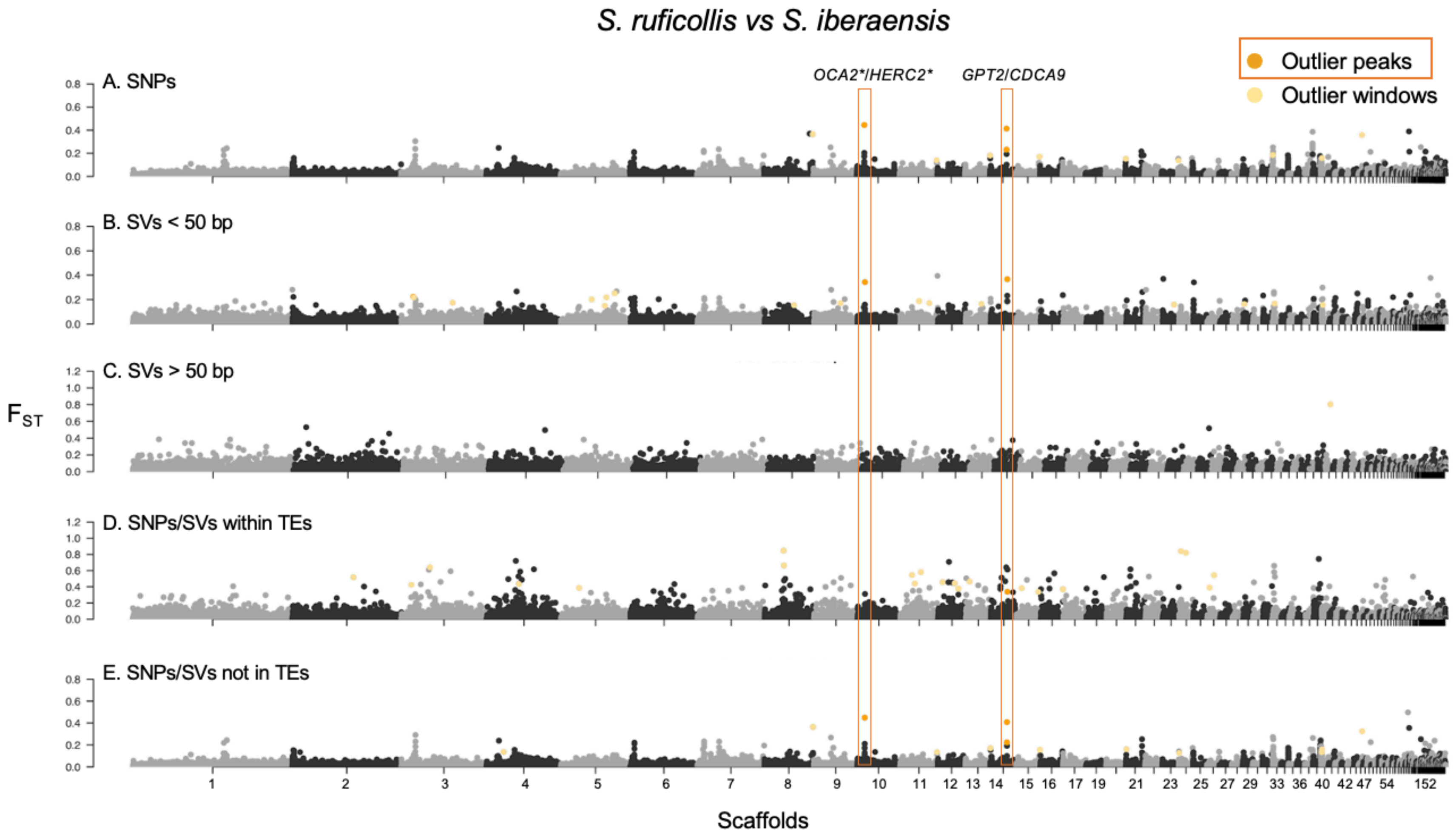

**Figure S24. $F_{ST}$ scan in 10 kb windows for the comparison between *S. ruficollis* and *S. iberaensis*.** Details as in Figure S12. There are two peaks in this comparison: one containing *GPT2* and *CDCA9* on scaffold 14, detected by all datasets except the long SVs, and another containing *OCA2*/*HERC2* on scaffold 10, detected by the same datasets as the previous peak, except the variants within TEs.

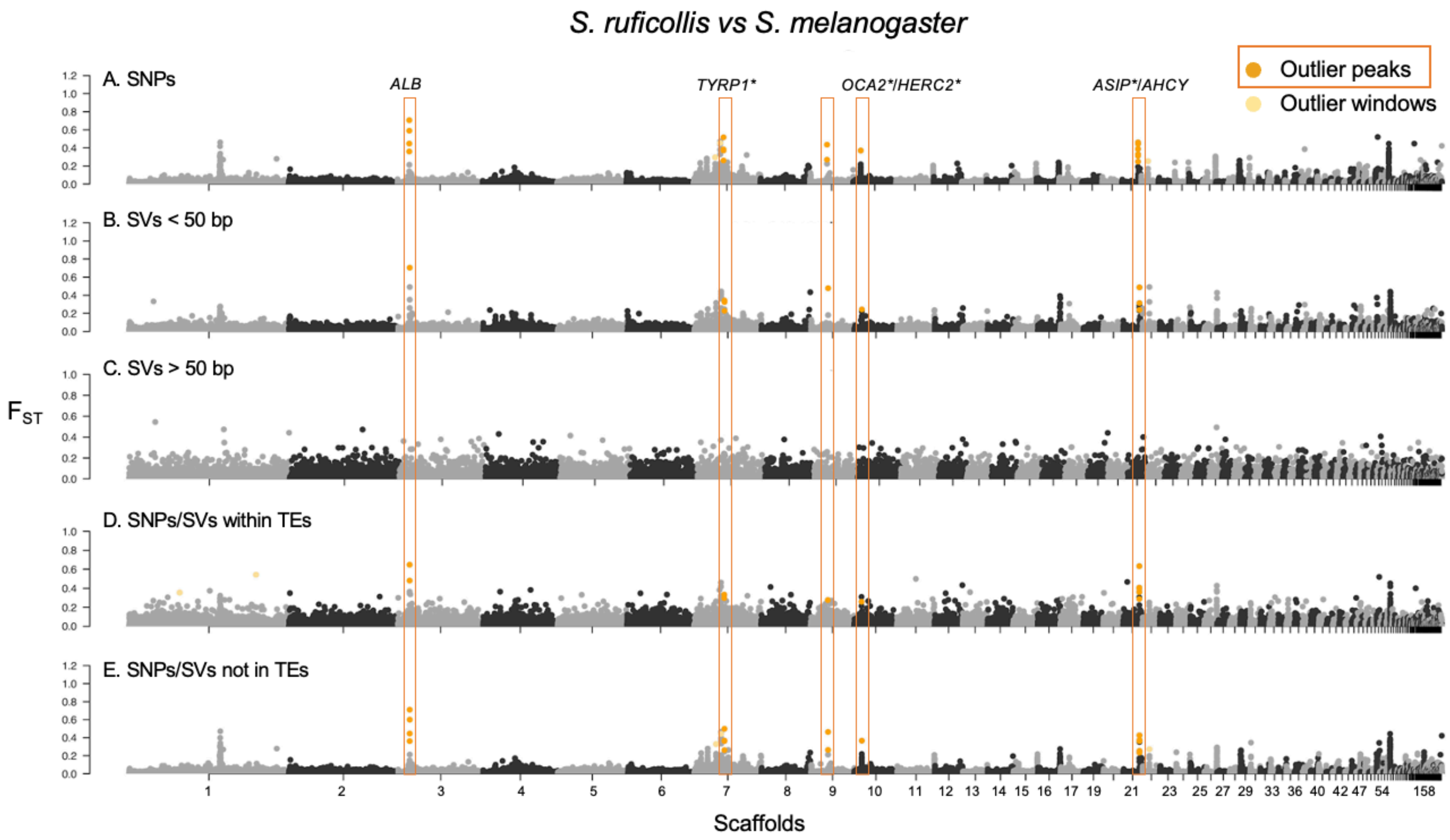

**Figure S25. $F_{ST}$ scan in 10 kb windows for the comparison between *S. ruficollis* and *S. melanogaster*.** Details as in Figure S12. There are five peaks in this comparison: one on scaffold 3 containing the gene *ALB*, one on scaffold 7 containing *TYRP1*, one on scaffold 9 with no annotated genes, one on scaffold 10 containing *OCA2*/*HERC2*, and another on scaffold 21 containing *ASIP* and *AHCY*. All peaks are detected by every dataset except the one including long SVs.

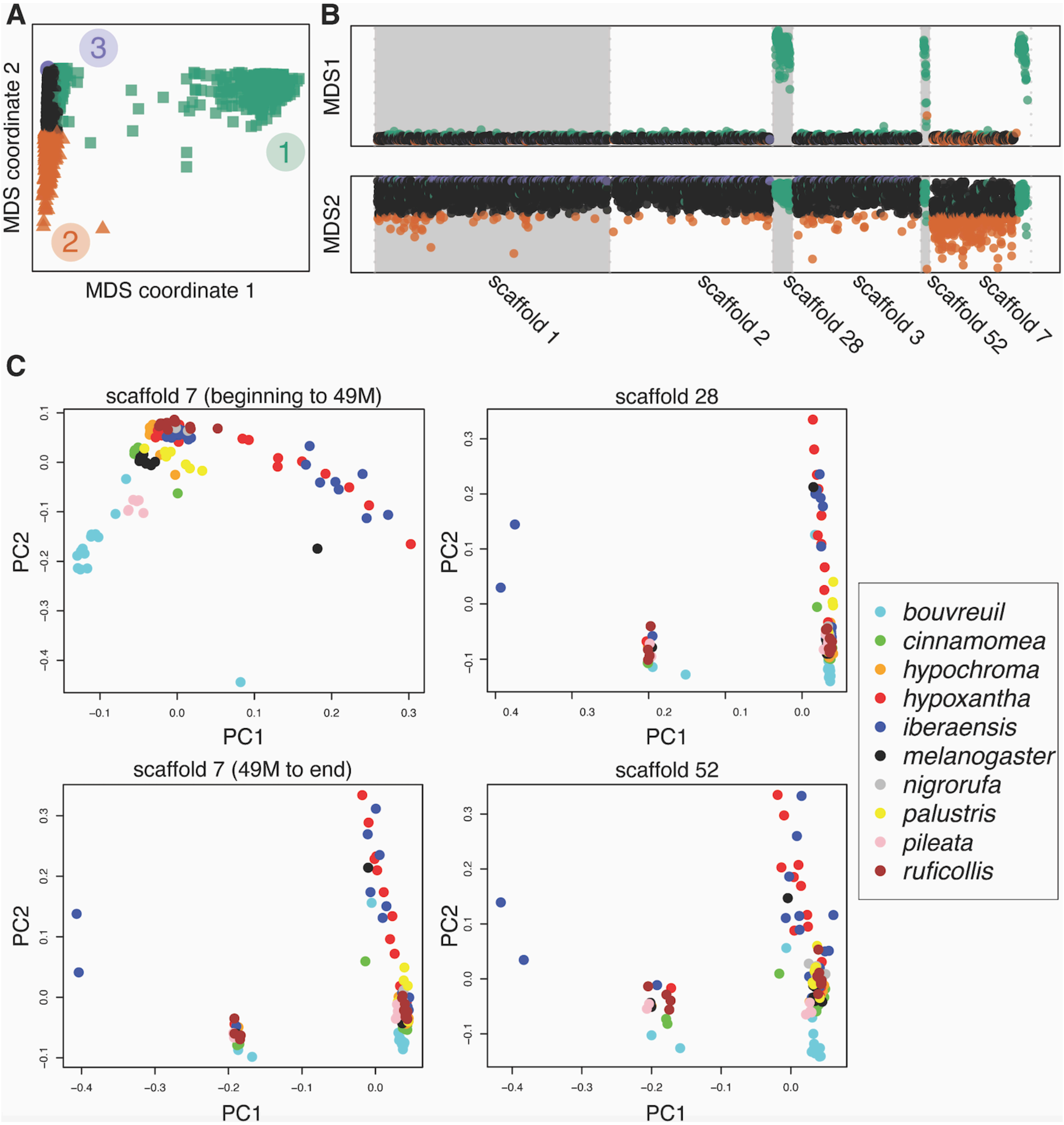

**Figure S26. Evidence of a large inversion on the Z sex chromosome. A)** Multidimensional scaling (MDS) analysis displaying the distance between local PCAs performed on windows (each 1,000 SNPs in length) from scaffolds 1, 2, 3, 7, 28, and 52 using lostruct in R. The outlier PCAs are displayed in each corner and color-coded (1, 2, and 3), and the genomic regions from which they originated are shown in **B)** with the corresponding colors (independently for MDS1 and 2). We performed the analysis on the 57 scaffold larger than 1 mb, that also contained at least 1,000 biallelic SNPs with no missing data. We identified scaffolds 28, 52, and the final portion of scaffold 7 (from positions ~49 Mb until the end) as outliers in MDS1 (shown in green). Here we display the three largest scaffolds (1-3) as non-outlier scaffolds to compare with those scaffolds identified as outliers. **C)** PCAs from the outlier scaffolds showing the different patterns observed on scaffold 7 in the first portion of the scaffold (beginning to ~49 Mb) and the final section (~49 Mb until ~56.3 Mb). The final portion of scaffold 7, 28 and 52 display a pattern suggestive of a chromosomal inversion, separating individuals with each version of the inversion on either extreme of PC1 and showing heterozygotes in the middle. The three scaffolds belong to the Z sex chromosome, and the two samples on the left side of the plot, likely homozygotes for one version of the inversion, are the same *S. iberaensis* individuals in each plot. Moreover, the presumably heterozygote individuals for the inversion in the middle of each plot (end of scaffold 7, 28, and 52) are also the same and belong to every species except *S. nigrorufa* and *S. palustris*. Because the outlier scaffolds belong to the same chromosome and individuals seem to show the same genotype for the inversion in the three different outlier scaffolds, we interpret this as evidence of a large inversion likely spanning the three scaffolds (~7.3 Mb of scaffold 7, 8.8 Mb of scaffold 28, and 2.4 Mb of scaffold 52). One version of the inversion seems to be much more common than the other (110 homozygote individuals vs. 2, with 15 heterozygotes). Therefore, even though the 2 homozygotes for the inversion are *S. iberaensis*, we cannot conclude that there is a pattern due to the low number of individuals in that group.

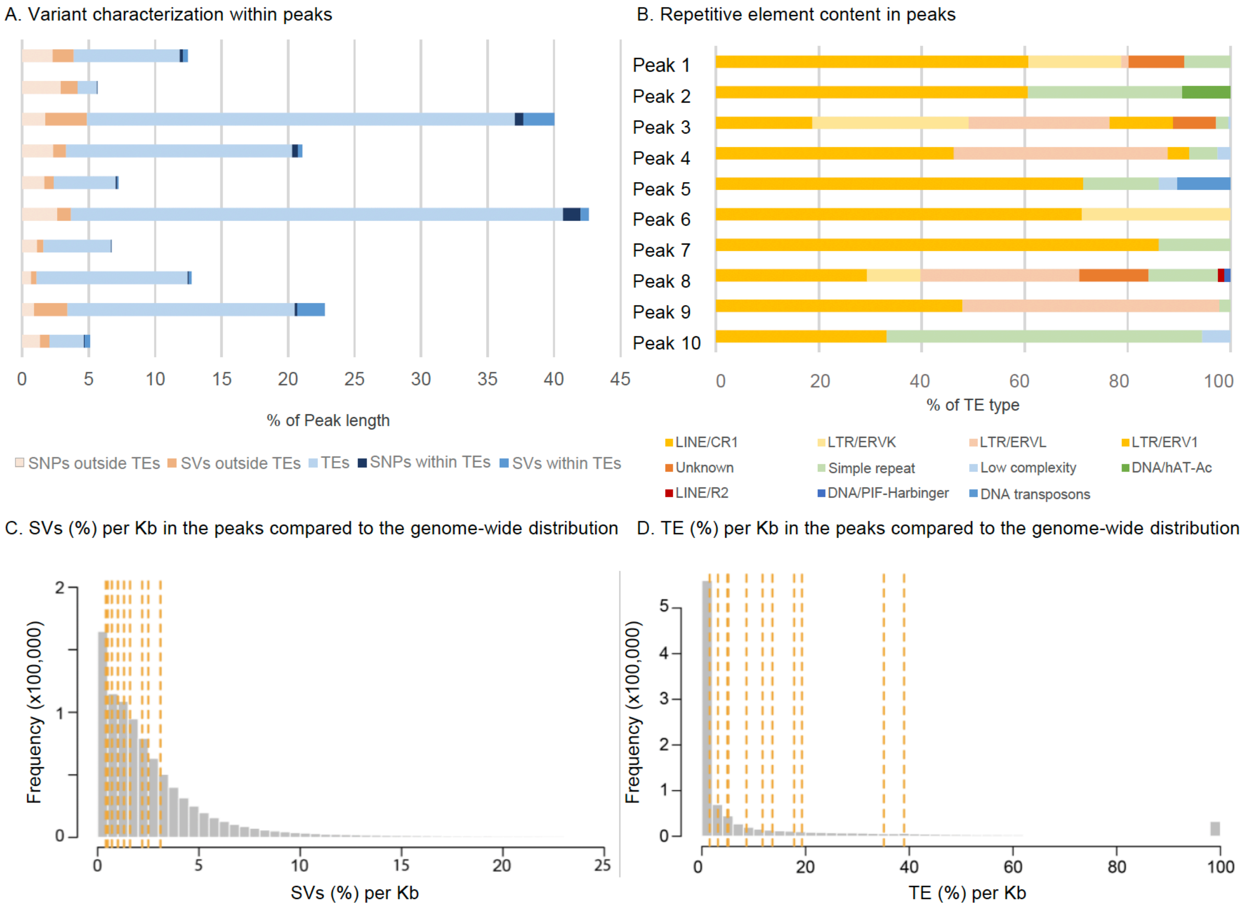

**Figure S27. Characterization of the different types of genetic variants within peaks. A)** Proportions of different types of genetic variants (SNPs and SVs), repetitive elements (including TEs) and their overlap within the peaks. **B)** Percentage composition of TE types within the peaks. Unclassified elements are labeled as "unknown". The peak IDs correspond to those in Table 1. **C)** Distribution of SV and TE (**D**) lengths as a percentage per kilobase, with that of peaks highlighted by dashed orange lines. None of the peaks fall within the top or bottom 2.5% of the distribution.

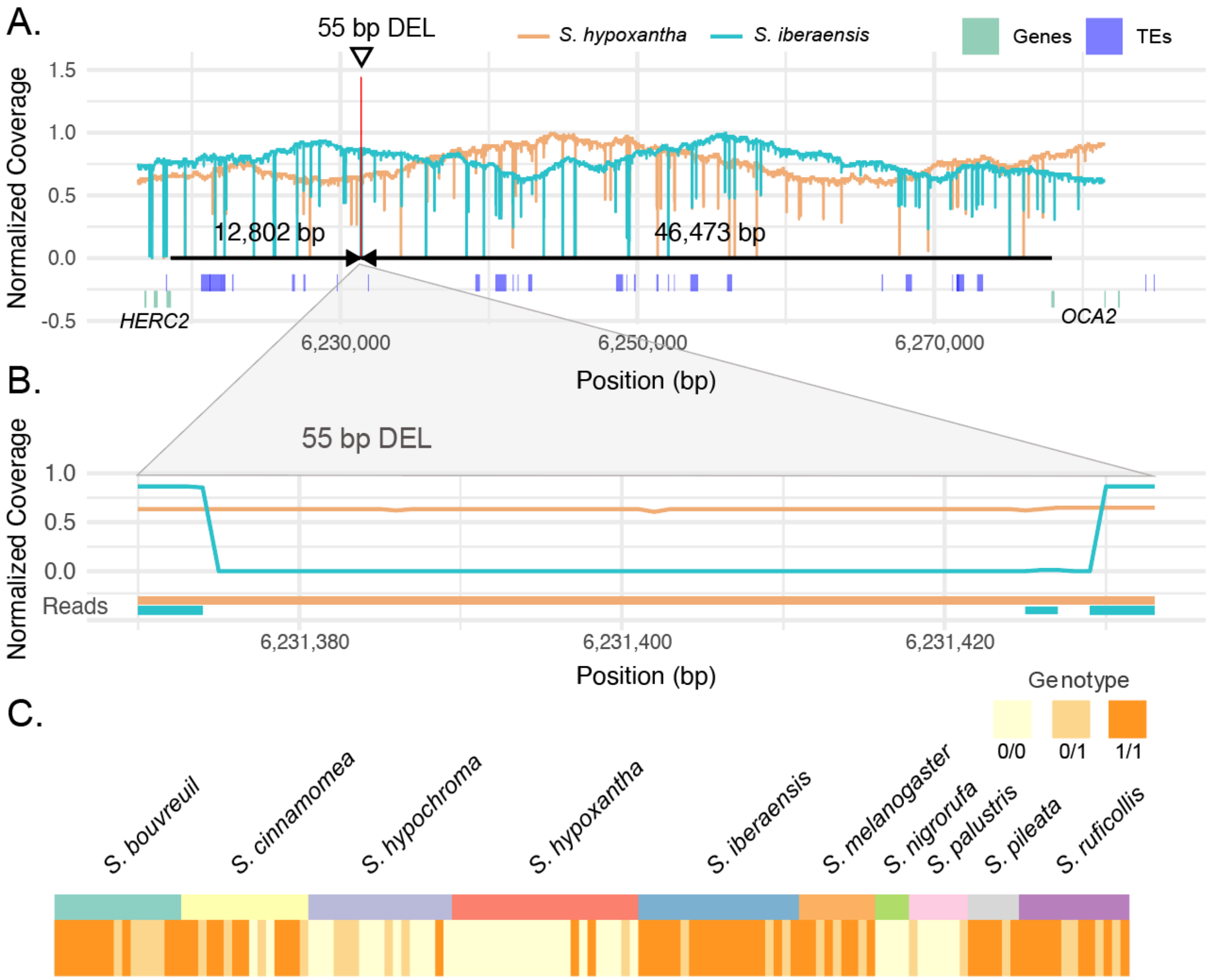

**Figure S28. Genomic context of the 55 bp deletion identified as an outlier in GWAS and $F_{ST}$ scans. A)** Genomic region surrounding the structural variant located at position 6,231,374 on scaffold 10 (red line marked with an arrow). The orange and blue lines represent the normalized coverage for one sample each of *S. hypoxantha* and *S. iberaensis*, respectively, based on long-read sequencing data. The purple tracks indicate transposable elements (TEs), while the green segments represent the genes *HERC2* and *OCA2*, with exons shown as vertical blocks and black arrows indicating the distances to the 55 bp deletion. **B)** Zoomed-in view of the 55 bp deleted region. Normalized coverage is shown by orange and blue lines, while the lines below the plot indicate reads supporting the deletion in one *S. iberaensis* individual. Teal represents *S. hypoxantha*, and orange represents *S. iberaensis*. **C)** Detailed genotypes at the deletion site for 127 individuals obtained using short-read sequencing data mapped to the pangenome. Vertical lines represent individuals grouped by species. Genotypes are color-coded: homozygous reference (0/0) in yellow, heterozygous (0/1) in light orange, and homozygous alternate (1/1; i.e., homozygous for the deletion) in orange.

PAL3372

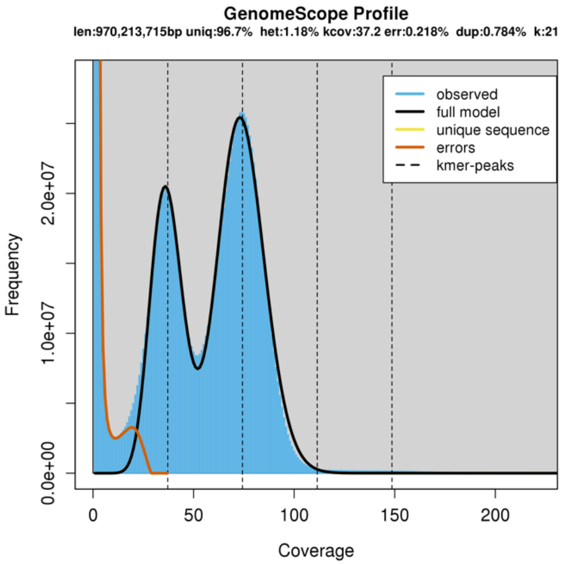

PAL3118

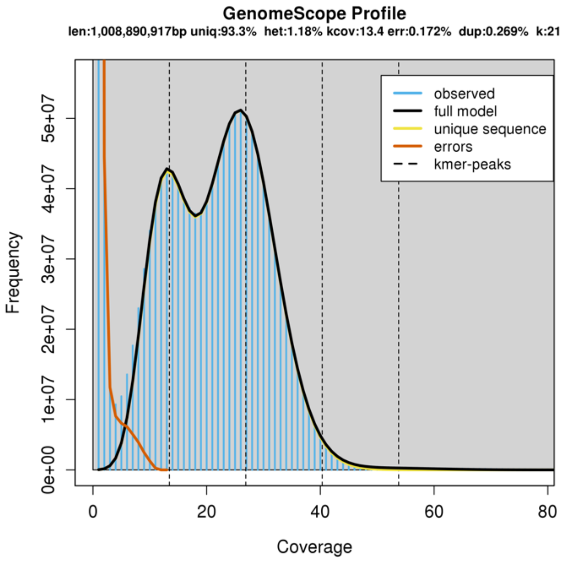

PAL3117

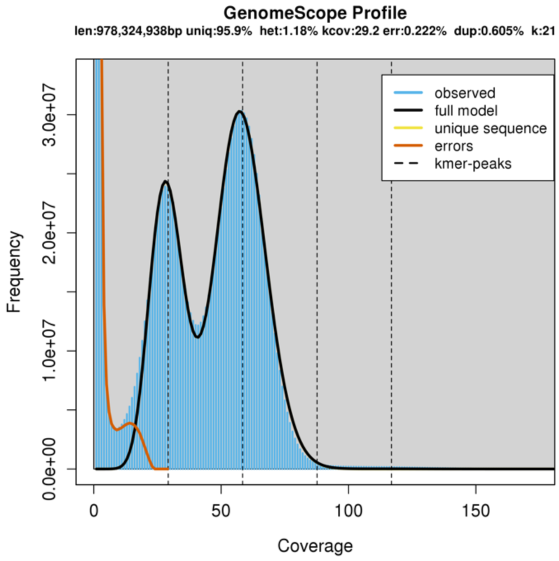

HYPOXB009666

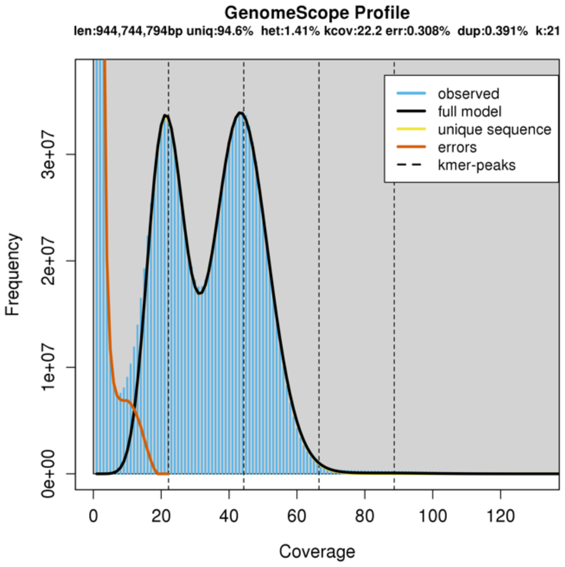

HYPOXB009684

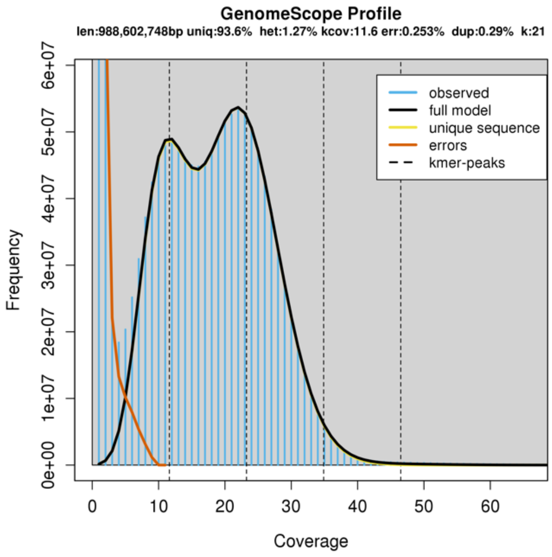

HYPOXB009213

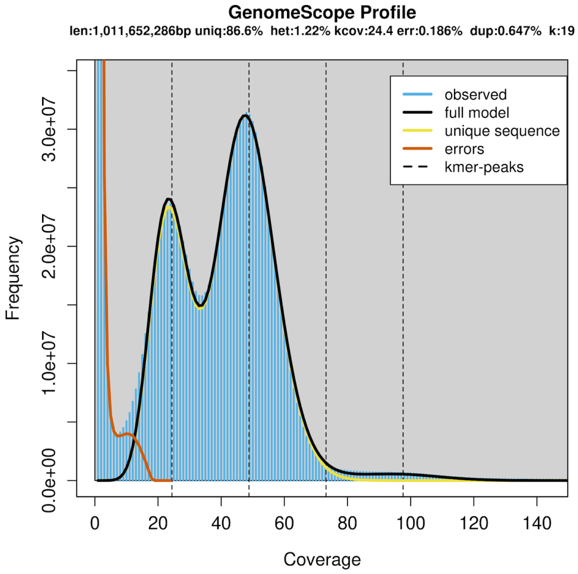

RUF3128

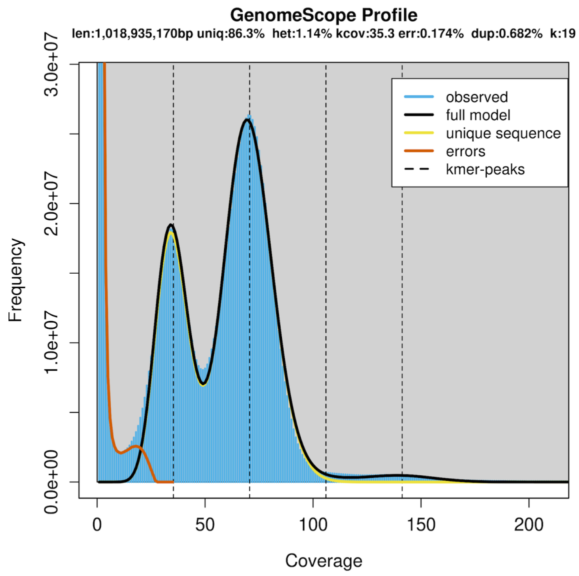

RUF3129

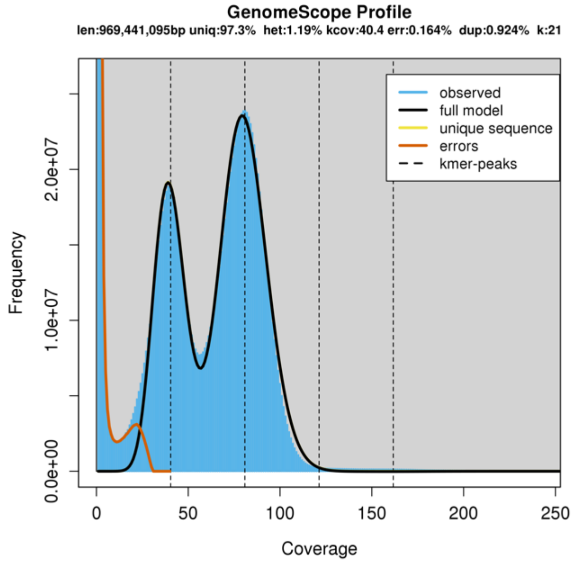

CRO3131

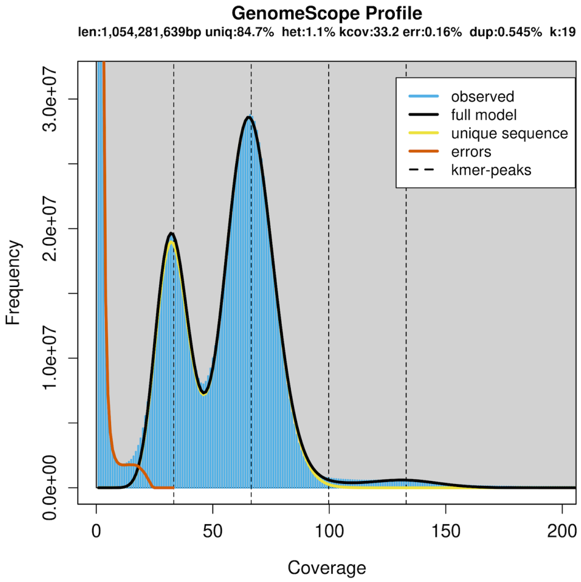

PIL3664

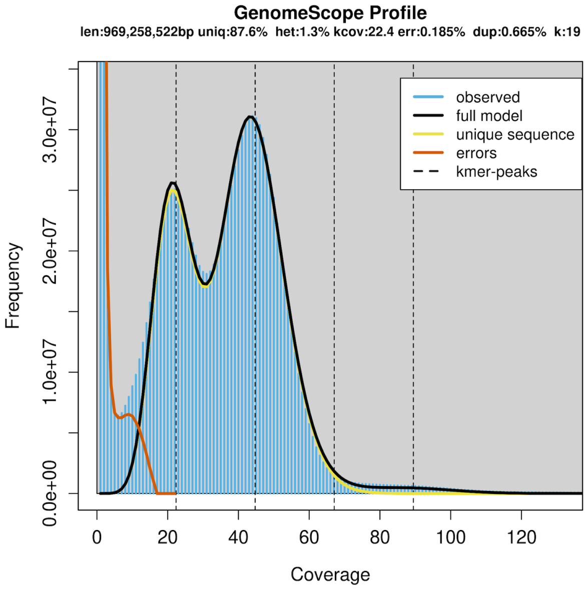

CIN3121

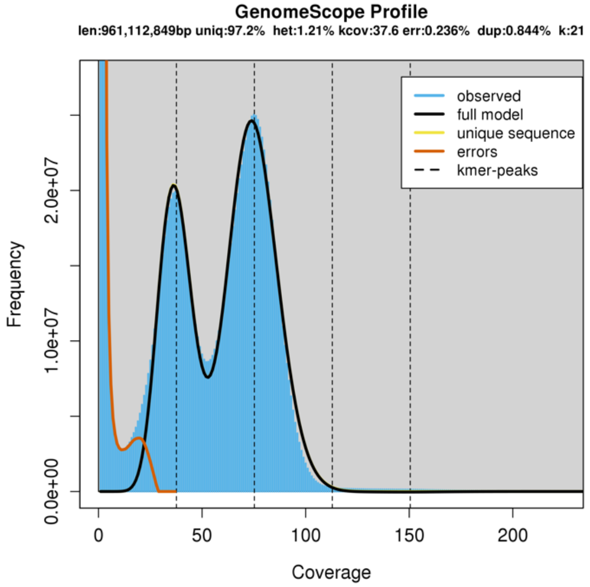

CIN3122

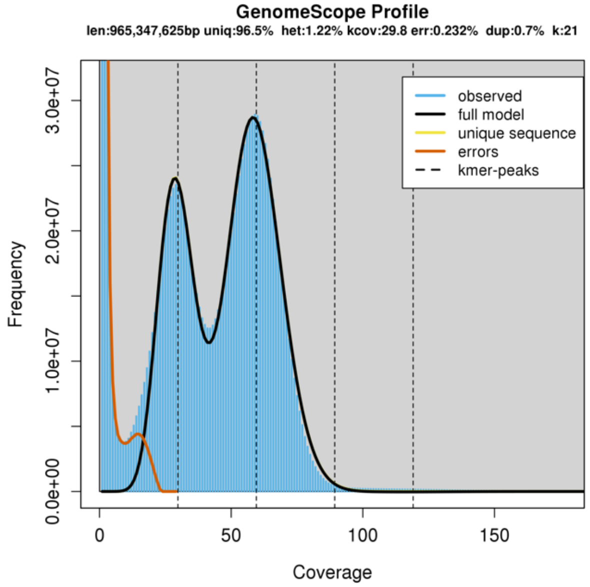

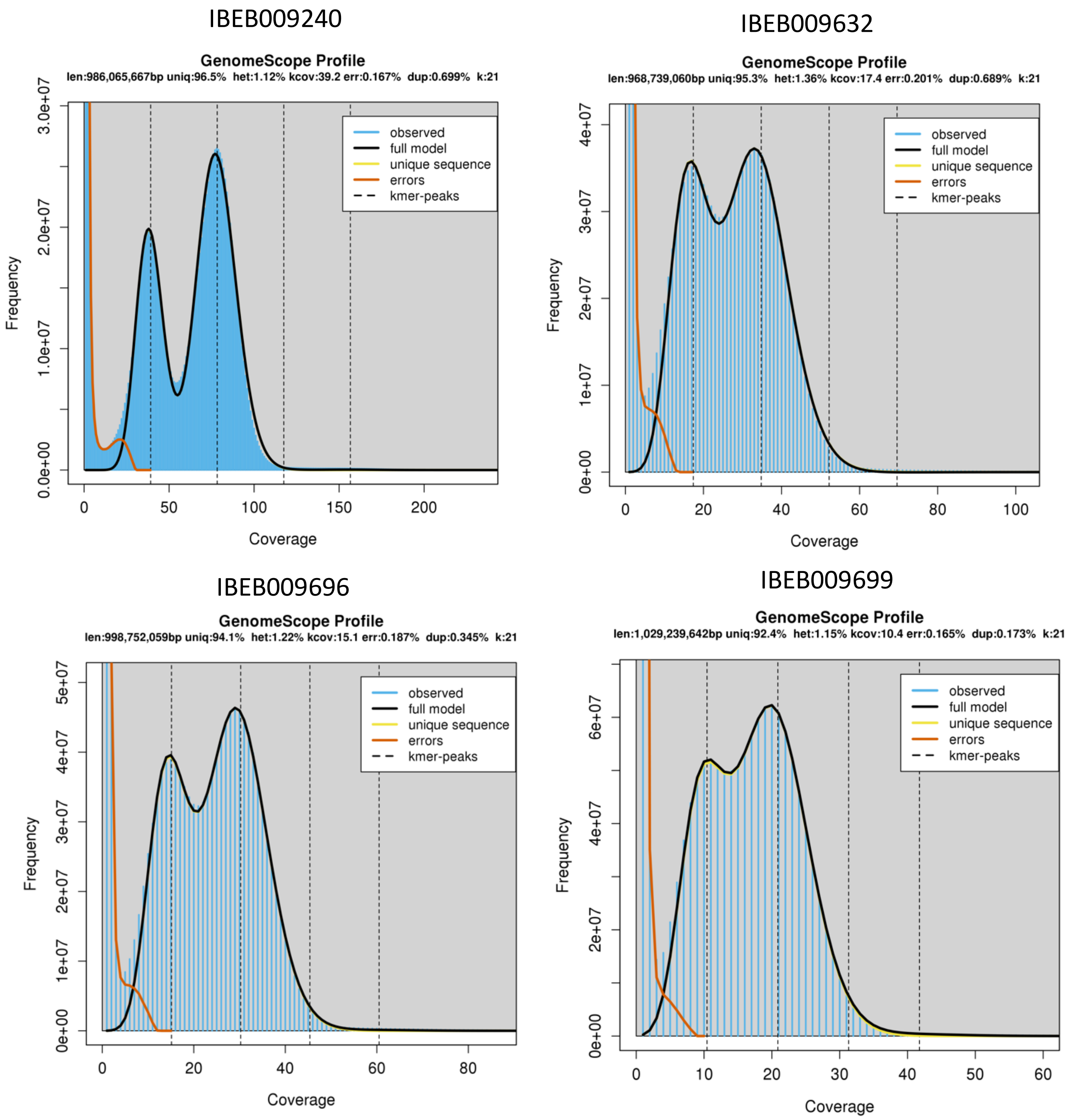

**Figure S29. GenomeScope analysis.** GenomeScope profiles were generated to estimate the genome sizes and heterozygosity of the assemblies, based on k-mer profiles obtained from Jellyfish histograms. These plots provide insights into the overall genome characteristics of each assembly, highlighting the distribution of k-mers and their frequency. In diploid genomes, the plot typically shows two distinct peaks corresponding to the two alleles at each locus: one for the heterozygous portion (with a lower k-mer frequency) and one for the homozygous portion of the genome (with a higher k-mer frequency). The k-mer size was initially set to 21 for all assemblies; however, for four datasets (HYPOXB009213, RUF3128, CRO3131, PIL3664), where convergence was not achieved, the k-mer size was adjusted to 19 to improve the accuracy of the estimates. These profiles offer a visual summary of the genomic complexity and quality of the assemblies.

### CIN3121

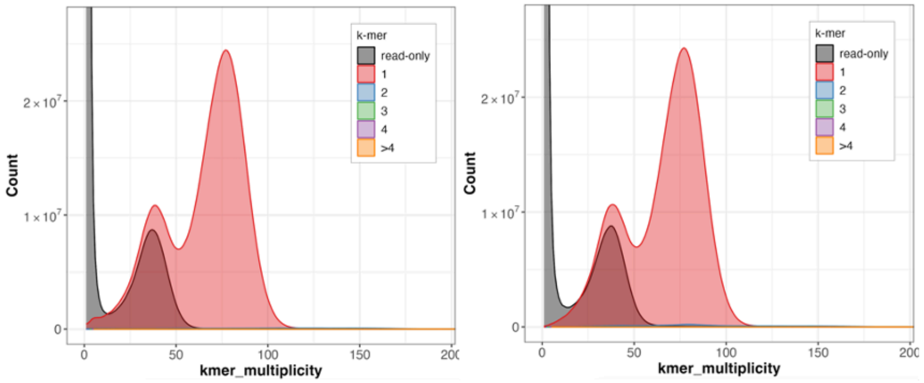

### CIN3122

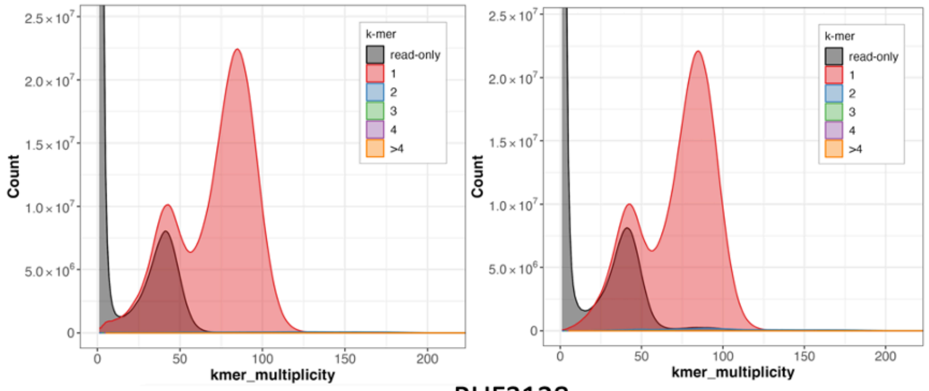

### RUF3128

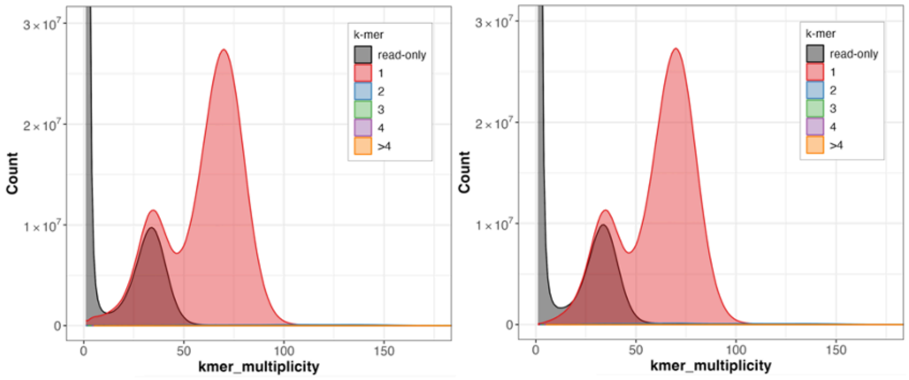

### RUF3129

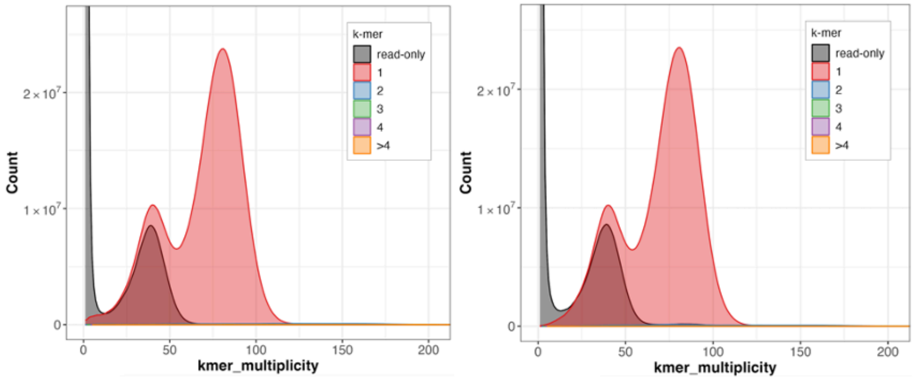

### HYPOXB009213

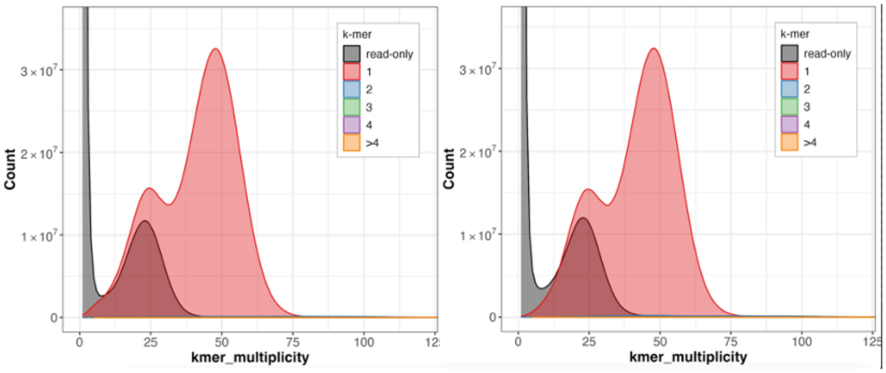

### HYPOXB009666

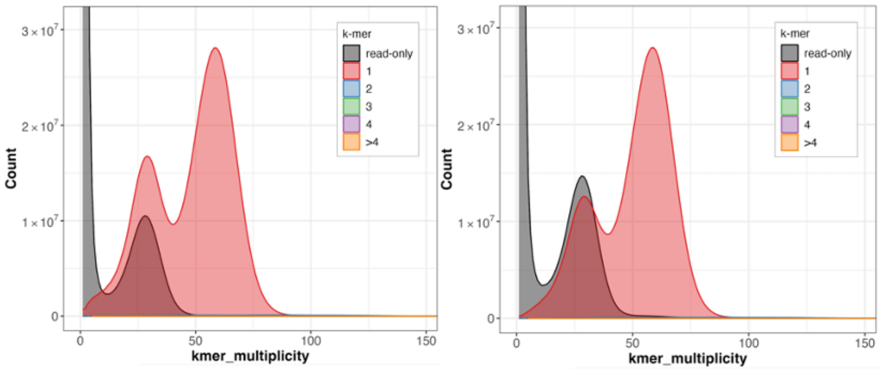

### HYPOXB009684

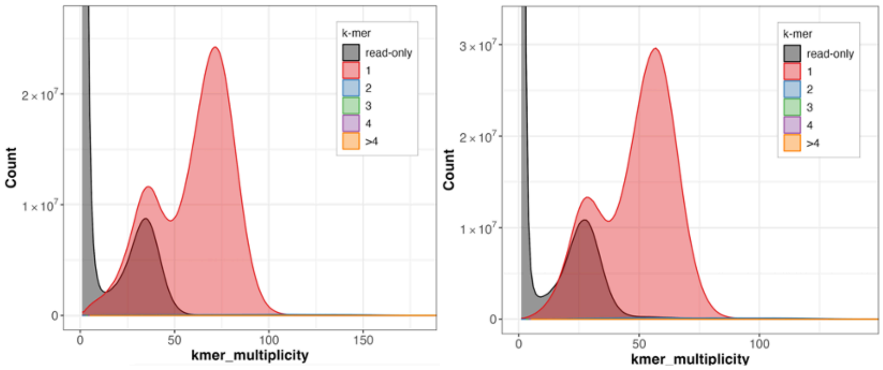

### CRO3131

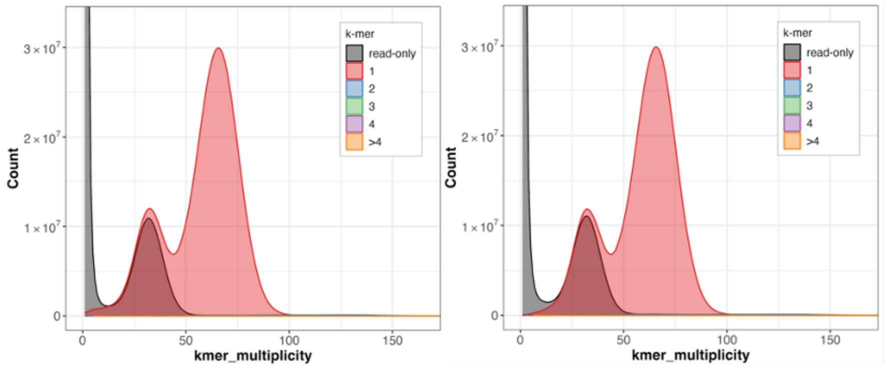

### IBEB009240

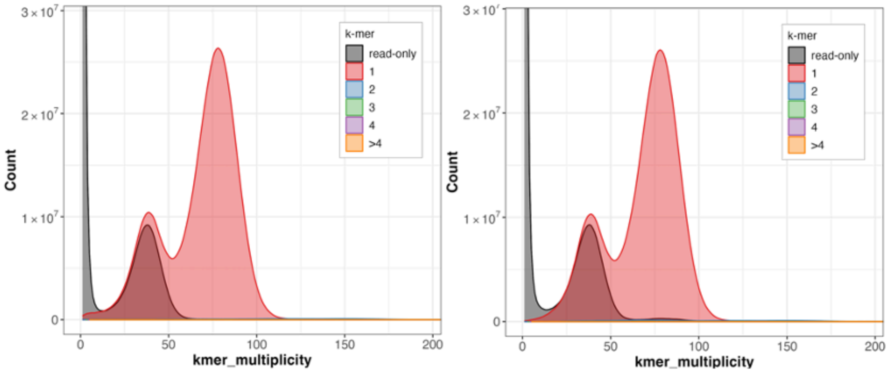

### IBEB009632

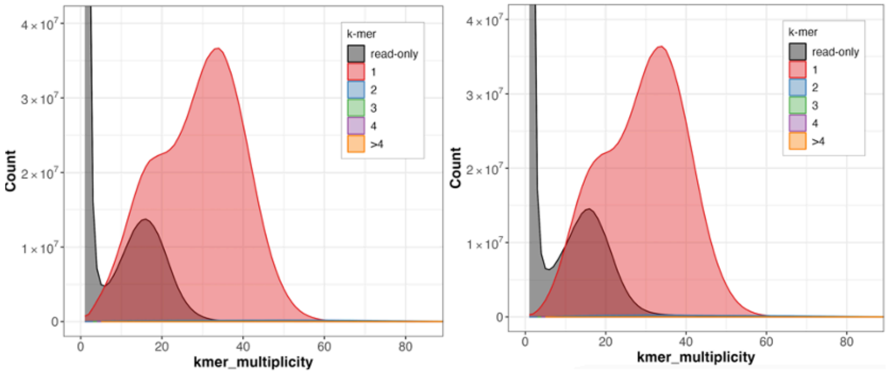

### IBEB009696

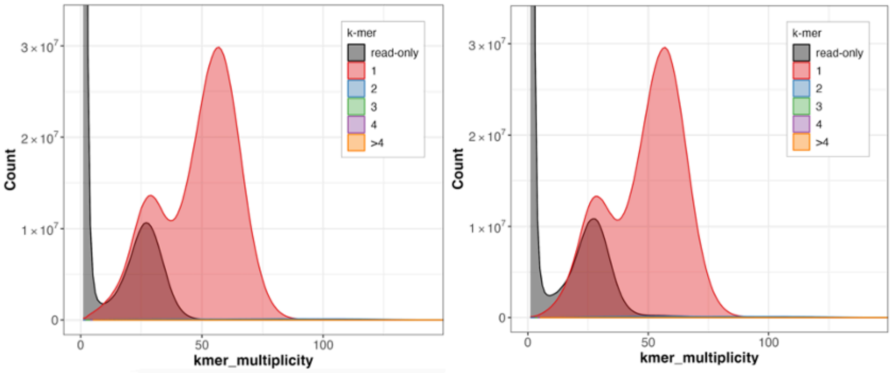

### IBEB009699

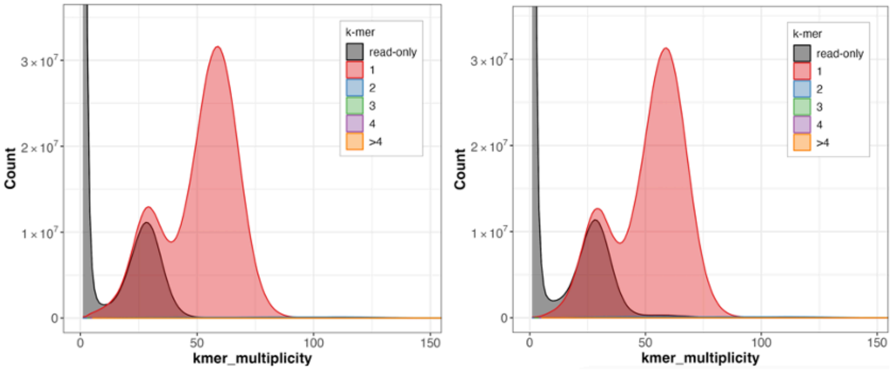

PAL3117

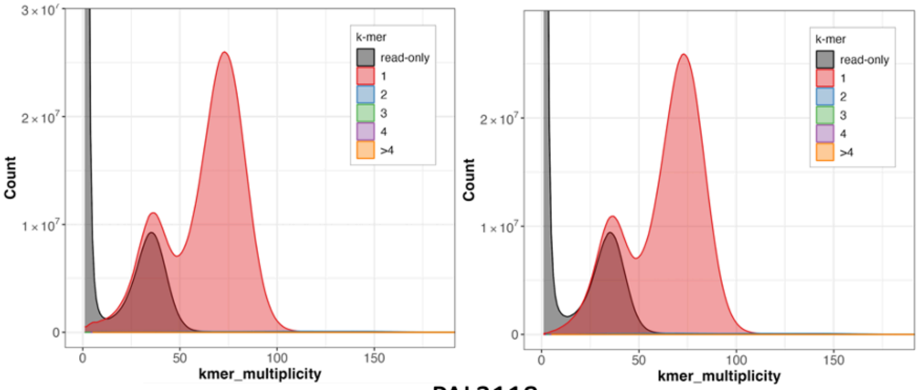

PAL3118

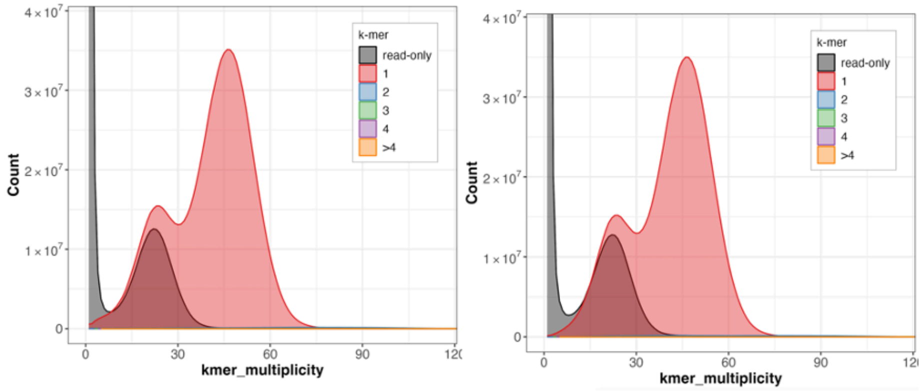

PAL3372

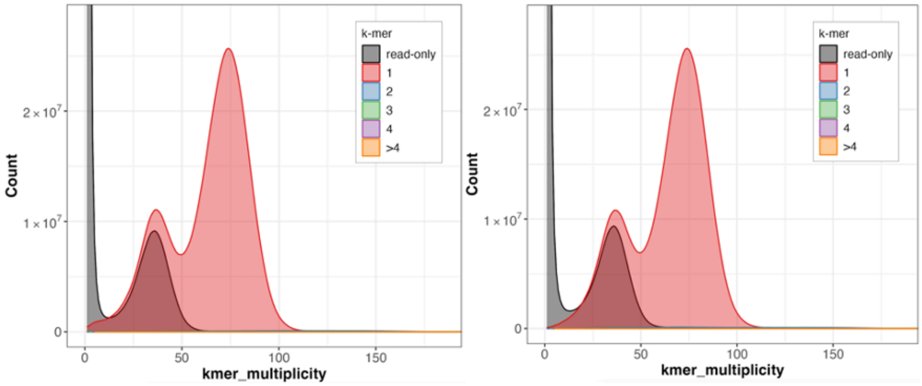

PIL3664

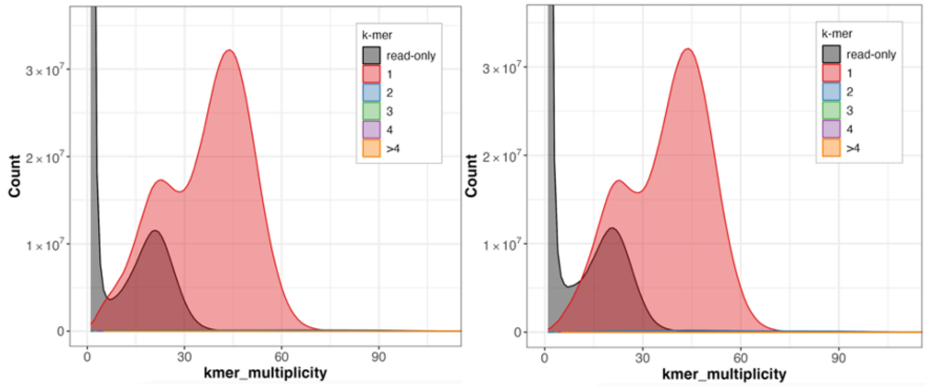

**Figure S30. Merqury analysis of assembly quality.** Merqury plots were generated to assess the quality of the primary (left) and alternate (right) purged assemblies. The plots show the distribution of k-mer multiplicity, with the x-axis representing how often each k-mer appears and the y-axis showing the count of k-mers with a given multiplicity. The two observed peaks correspond to heterozygous and homozygous sites, from left to right. As expected, in each assembly, k-mers are found only once, indicating that for homozygous sites, each allele is represented in either the primary or alternate assembly. The grey "read-only" peak within the heterozygous peak represents k-mers absent in that assembly because they are present in the other.
